# Modeling Patient-Reported Pain Trajectories with Frequent Minimum and Maximum Scores

**DOI:** 10.64898/2026.05.07.721507

**Authors:** Yanxi Liu, Richard E. Harris, Daniel Clauw, Emine Bayman, Andrew Leroux, Martin A. Lindquist, the A2CPS Consortium

**Affiliations:** Department of Biostatistics, Johns Hopkins Bloomberg Schools of Public Health, Baltimore, MD, USA; Department of Anesthesiology and Perioperative Care, School of Medicine, University of California at Irvine, Irvine, CA, USA; Department of Anesthesiology, University of Michigan Medical School, Ann Arbor, MI, USA; Clinical Trials and Data Management Center, Department of Biostatistics, University of Iowa, Iowa City, IA, USA; Department of Biostatistics & Informatics, Colorado School of Public Health, USA

**Keywords:** Chronic Pain, Beta-Binomial, Zero Inflation, Bayesian, Ecological Momentary Assessment, Random Effects, Longitudinal Model

## Abstract

Chronic pain is a widespread public health issue that imposes substantial health, emotional, and economic burdens on individuals and communities. Because pain is subjective and lacks objective biomarkers, it is typically measured using patient-reported scores, often on a numerical scale from zero to ten. Increasingly, pain studies use ecological momentary assessment, with multiple daily assessments over days and across study phases (e.g., a series of baseline and post-intervention assessments). These data frequently show many ratings at the extremes (i.e., at minimum or maximum pain scores), commonly referred to as zero- and one-inflation in the statistical literature, along with considerable within-person variability both within and across days. These phenomena present challenges for statistical analyses, as they violate assumptions of most commonly used statistical techniques (e.g., the normality assumption of linear mixed models). We propose a Bayesian beta-binomial mixed-effects model for modeling potential zero- or one-inflated pain scores while accounting for variability using random effects on the mean and variance parameters across subjects. A simulation study demonstrates that the method accurately estimates model parameters across realistic sample sizes, time points, and zero- and one-inflation levels. An application to data from two longitudinal pain studies demonstrates that the model fits the data better and, when correctly specified, yields accurate uncertainty intervals for longitudinal changes in pain compared to existing models, especially for zero- and one-inflated outcomes. Additionally, the model directly estimates the probability of clinically meaningful pain events. The proposed method provides a powerful statistical framework for studying the patient-reported pain trajectories.

**Summary:** The proposed beta-binomial mixed-effects model accounts for potential zero- and one-inflation in the longitudinal trajectories of bounded pain scores, and subject-specific variability of repeated measures.

## 1 Introduction

Chronic pain, defined as pain persisting for more than three months or co-occurring with another chronic condition, is a major public health concern that reduces well-being and quality of life [1,4,13,28,30,37]. In 2021, 20.9% of U.S. adults (51.6 million) had chronic pain, and 6.9% (17.1 million) had high-impact chronic pain (i.e., pain that severely restricts daily activities) [33]. Despite its prevalence, the mechanisms underlying chronic pain remain poorly understood, underscoring the need for statistical methods that characterize risk factors and pain trajectories.

Pain is typically measured using patient-reported numerical rating scales (e.g., 0 “no pain” to 10 “worst imaginable pain”) and collected through interviews [14], questionnaires [10], or ecological momentary assessment (EMA), which captures assessments over days [17]. These data frequently exhibit many ratings at the extremes (i.e., many 0s and/or 10s), a feature known as zero- and one-inflation (Figures 1–2). Empirically, zero-inflation might be present in population samples with many pain-free subjects. Conversely, one-inflation may be expected in data from trauma patients. Moreover, pain varies within and across days, complicating analysis of repeated measures.

**Figure 1.**
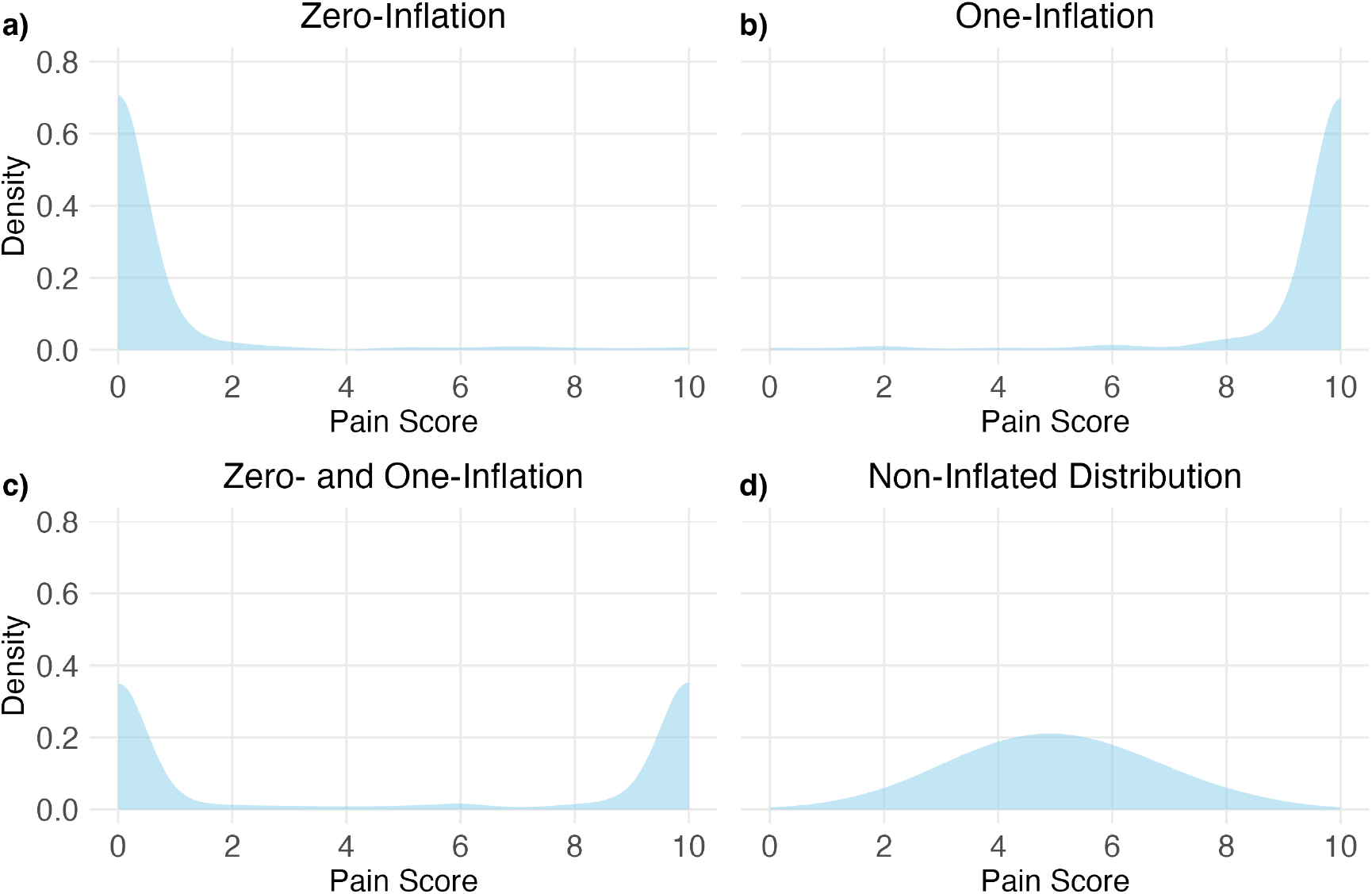
Example plots of a) zero-inflated, b) one-inflated, c) zero- and one-inflated, and d) non-inflated data sampled from the beta-binomial distribution.

**Figure 2.**
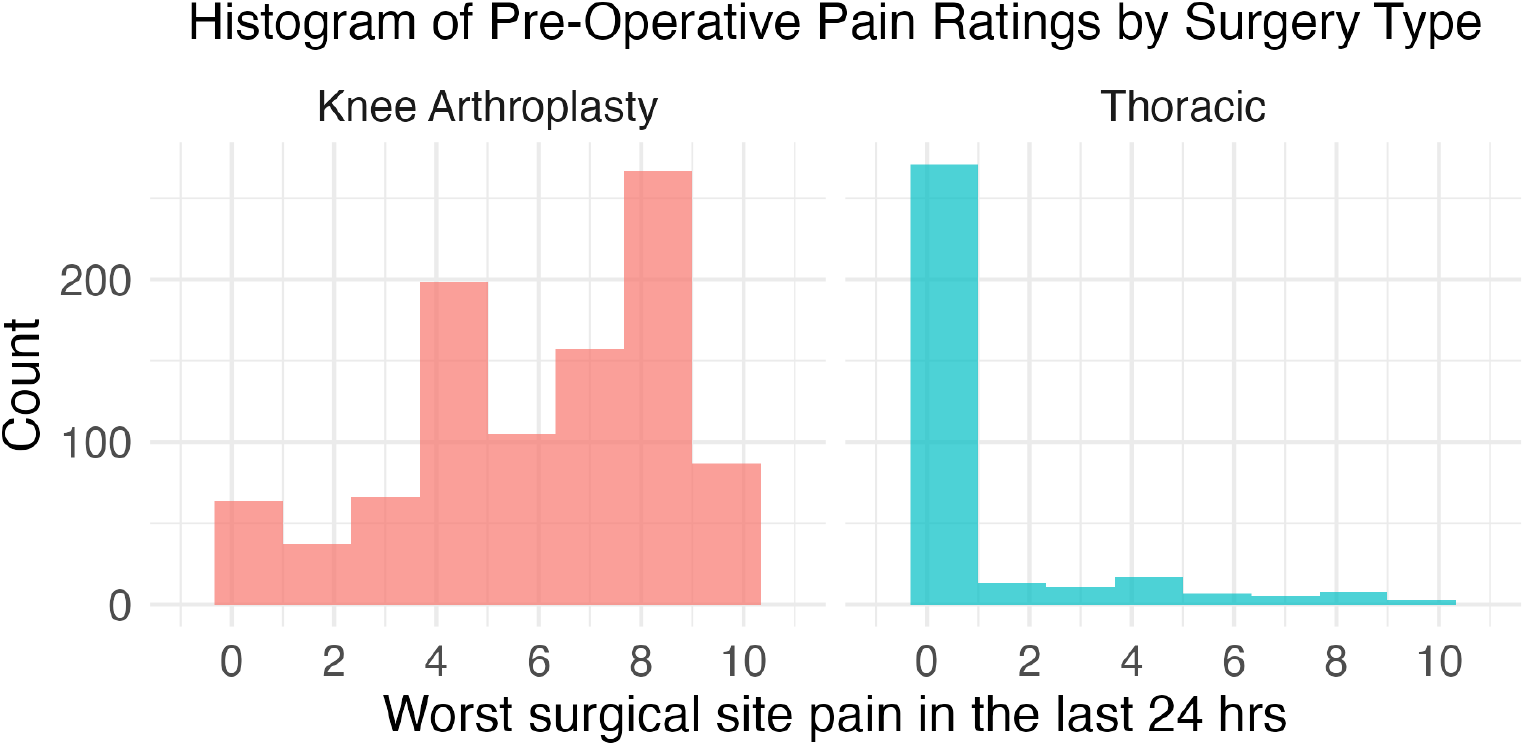
Histogram of worst surgical site pain in the past 24 hours. Baseline data were measured on 1043 total knee arthroplasty (TKA) subjects and 358 thoracic subjects before surgery in the Acute to Chronic Pain Signatures (A2CPS) study (Release 2.1.0; January 2025); worst surgical site pain scores in the past 24 hours were available for 981 TKA and 335 thoracic subjects. A high proportion of thoracic surgery patients reported no pain at the future surgical site, whereas pain scores among TKA patients were more widely distributed.

Many studies have dichotomized scores and analyzed pain using logistic regression [11,38], discarding information, reducing statistical power and estimates’ precision [25]. Linear mixed-effects models (LMMs) preserve the numerical scale [8,22] and accommodate repeated assessments through random effects, but are ill-equipped for bounded-integer outcomes with frequent extreme values and assume a continuous distribution. Other methods (e.g., structural equation modeling and k-means clustering) [10,22] similarly do not adequately address extreme values.

Zero-inflation can arise from structural zeros (true absence of pain), or overdispersion (variability exceeds model expectations). Mixture models like zero-inflated and hurdle models [24,29] target structural zeros and have been used to model pain [18,20,40]. Extensions like zero-inflated beta-binomial and zero/one-inflated beta regression [21,27] further address overdispersion. However, mixture models are most appropriate when zeros arise from distinct generating processes [41] and can challenge interpretation [18,24,40], for example when covariates affect model components differently. In many pain contexts, excess boundary values are more plausibly explained by more variability than expected than by a mixture of “max pain”, “some pain,” and “no-pain” states. We therefore favor models that work for non-inflated data and address boundary behaviors within a single data-generating process.

To overcome the issues in existing models, we propose a beta-binomial mixed-effects (BBME) model for analyzing zero- and/or one-inflated pain scores. The beta-binomial (BB) distribution, with mean and dispersion parameters, naturally accommodates overdispersion and reproduces excess boundary values without invoking structural inflation. We model within- and between-subject variability via random effects, making BBME well-suited to EMA designs. Although BB models have been used across disciplines [15,16,31], to our knowledge they have not been applied to pain.

The model was estimated in a Bayesian framework [7]. Simulations demonstrated accurate estimation for both group differences (e.g., covariate effects) and subject-specific predictions across plausible parameter values at realistic sample sizes. We applied BBME to two longitudinal pain datasets, one with pronounced zero-inflation and one without. Results showed that BBME captures longitudinal changes in pain while appropriately handling boundary values.

## 2 Methods

The BBME model is well suited for pain scores recorded on fixed scales, such as 0-10. This statistical framework readily and directly models pain data with many (or few) reported scores at the low and/or high end of the scale, while also models excess variability that is common in pain data [8,22]. Thus, it can be reasonably applied to model pain in highly heterogeneous pain populations. Random effects specification is flexible and accounts for both between- and within-person variability. Additionally, Bayesian estimation readily provides subject-specific predictions [26,34]. Together, the flexibility of the BBME model generalizes beyond pain research to any bounded-integer outcome, a common feature of EMA data which in general have repeated measures and hierarchical structure, and thus may provide a means for extracting greater insights from the data type.

### 2.1 Beta-Binomial Mixed Effect Model

We begin by introducing the notation used throughout the paper. Suppose we have *I* subjects, each observed at *J* study time points. At each time point, subjects provide *K* repeated assessments. For example, a subject might complete a daily survey for one week at two study points: two weeks before surgery and three months after surgery. In this case, *J* = 2 and *K* = 7. Let *y*_*ijk*_ denote the pain score for subject *i* (*i* = 1, …, *I*) at time point *j* = 1, …,*J*) during the *k*-th assessment *k* = 0, …, *K* − 1).

Because pain scores are recorded on a fixed whole-number scale (e.g., 0-10), rather than making a continuous (e.g. normal) approximation, we model the observed data directly using a BB distribution:

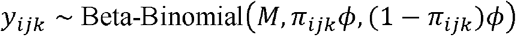

where *M* is the maximum possible score (for the NRS 0-10 scale, *M* = 10), 0< π_*ijk*_ <1 is a location parameter (determining the expected level of pain of subject at time point *j* during the *k*-th assessment), and ϕ> 0 is the dispersion parameter (controlling how variable the pain scores are around the expected level).

In practical terms, larger values of π_*ijk*_ shift the distribution of pain scores toward higher values. The parameter *ϕ* controls how tightly or loosely the data cluster around that mean. One way to better understand this parameter is through an alternative parametrization of the distribution. First, the probability of pain, denoted as *p*_*ijk*_, is drawn from a Beta (π_*ijk*_ *ϕ*, (1 − π_*ijk*_ *)ϕ*) distribution. Then, the observed pain score is drawn from a binomial distribution with probability *p*_*ijk*_. This structure allows the BB distribution to capture extra variability compared to a standard binomial. When *ϕ* is large, the model behaves like an ordinary binomial distribution. When *ϕ* is small, the data is more dispersed, allowing for more clustering at the extremes (i.e., zeros and maximum scores).

Figure 3a-f illustrates how different values of π and *ϕ* can produce zero-inflated, one-inflated, or more evenly distributed outcomes. For example, with a small π and low *ϕ*, most scores will cluster at zero (Figure 3a). Increasing either π or *ϕ* reduces the number of zeros (Figure 3b, c, d), while moderate values of both yield a more symmetric, non-inflated distribution (Figure 3e). This is explored further in Appendix 1.

**Figure 3.**
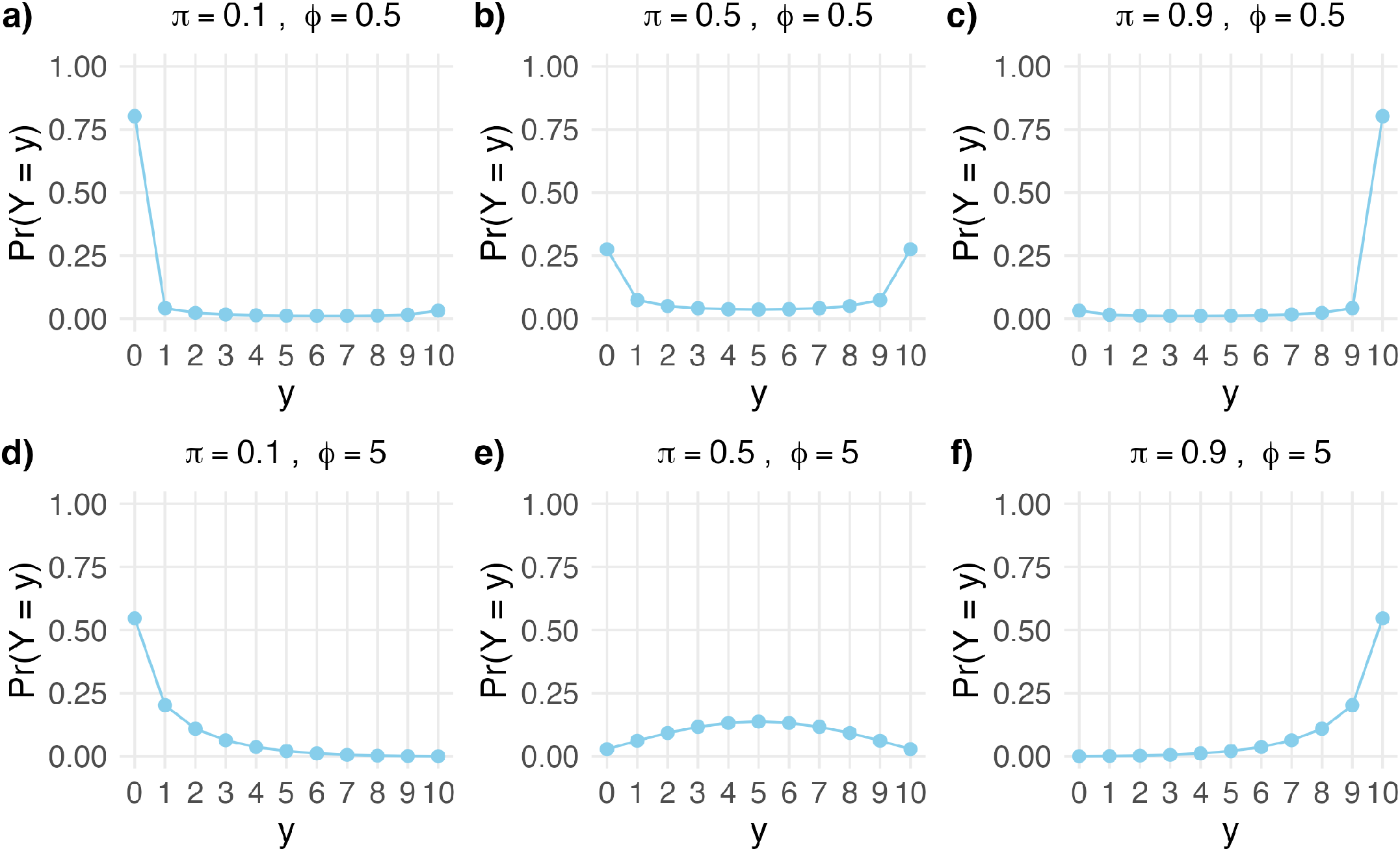
Probability mass functions of the beta-binomial distribution with maximum score *M* = 10 under varying location (*π*) and dispersion (*ϕ*) parameters. By adjusting these parameters, the beta-binomial distribution can generate zero-inflated, one-inflated, and non-inflated distributions.

To allow each subject to have their own pain pattern over time, we link the location parameter π_*ijk*_ to subject-specific random effects using a logit function:

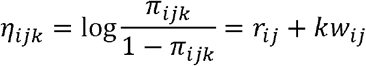

where *k* indexes the within-time-point assessment *k* = 1, …, *K*). Here, **r**_*i*_ =[*r*_*i*1_ *r*_*i*2_ … r_*ij*_]^*t*^ represents a *J* × 1 vector of random intercepts that describe each subject *i*’s initial pain tendency at time point *j*), and *w*_*i*_ is a *J* × 1 vector of random slopes that describe how subject *i*’s pain changes across repeated assessments within each time point). These subject-level random effects are assumed to independently follow a multivariate normal distribution with a 2*J* × 2*J* covariance matrix Σ, which allows us to capture correlations between pain scores at different time points, that is:

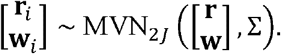

This covariance matrix allows pain patterns at different study time points to be related to one another. The off-diagonal elements represent the relationship between pain-score location parameters across time points. For example, a positive correlation between pre- and post-surgery intercepts implies that subjects with higher baseline pain are also more likely to report higher pain after surgery. Figure 4 demonstrates how varying *r, w*, and *ϕ* generates distributions covering zero-inflation, one-inflation, and non-inflated cases. We refer to this parameter specification on the location parameter as the full model in the following text.

**Figure 4.**
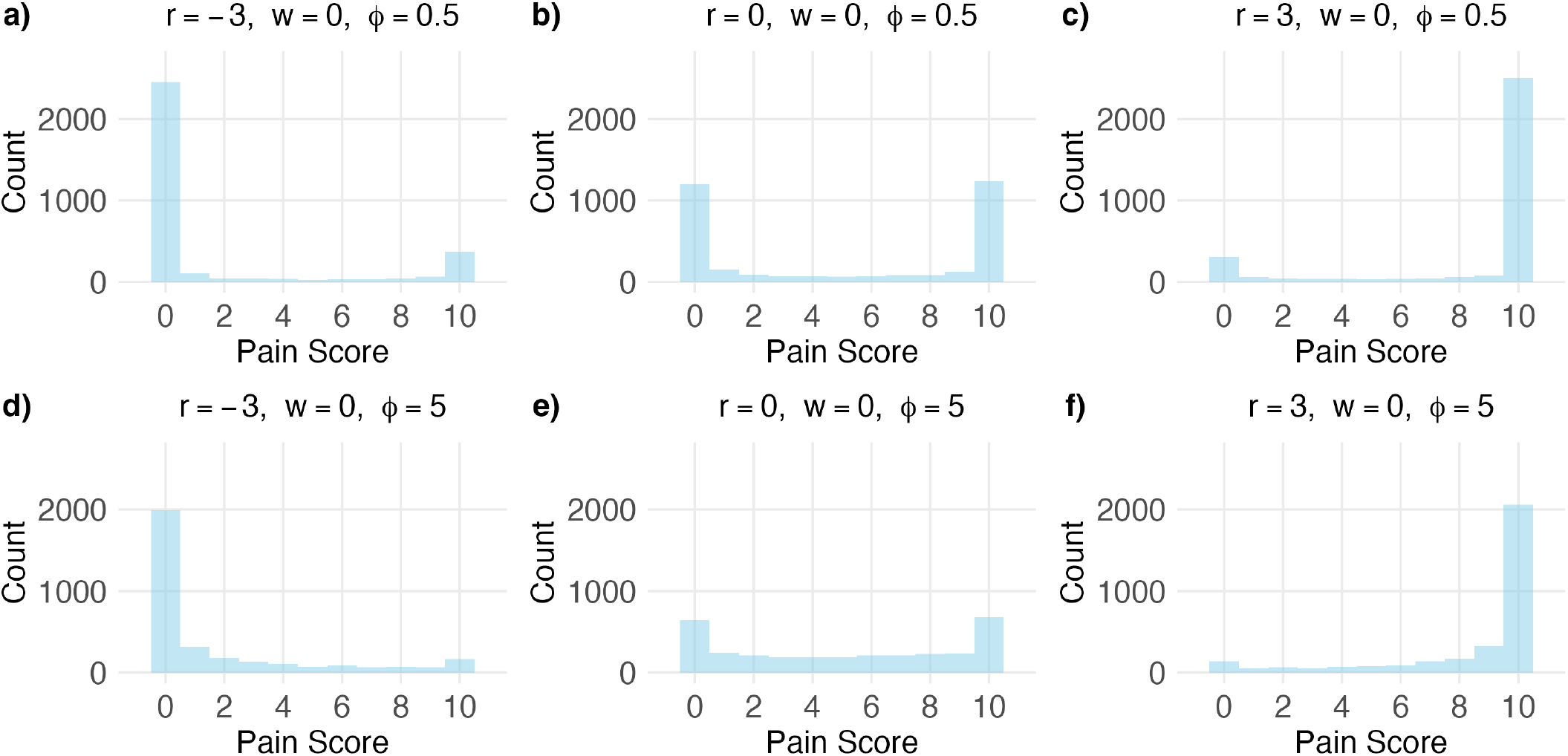
Distribution of randomly generated samples of pain scores with *I* = 500 subjects, *J* = 1 time point, and *K* = 10 scores at each time point per subject. Each pain score follows a beta-binomial distribution with maximum score *M* = 10, varying parameter values *r* and *ϕ*, and fixed *w* = 0. The random effects [*r*_*i*_ *w*_*i*_]^*T*^ are sampled from a multivariate normal distribution with mean [*r w*]^*T*^ and covariance matrix 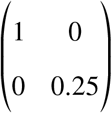.

The dispersion parameter *ϕ* described above is assumed to be the same for all subjects and time points. However, the model can be easily extended. For example, we may allow dispersion to vary by subject *(ϕ*_*i*_), by time point *(ϕ*_*j*_), or by subject and time point *(ϕ*_*ij*_). Conversely, simpler versions of the model can be fit by having only random intercepts. A range of model variations are described in Appendix 2.

The model was estimated in a Bayesian framework using Hamiltonian Monte Carlo implemented in the Stan programming language via the rstan package [7]. For comparison, we also fitted the model using the brms package in R [6] which automates the writing of Stan code using syntax for fitting generalized linear mixed models in R, as well as a frequentist approach implemented using the glmmTMB package in R [5]. Additional details of the model fitting procedure can be found in Appendix 3.

### 2.2 Data Simulation and Model Validity

A series of simulations was conducted to evaluate the validity of the proposed BBME model estimated in Stan. In the zero-inflation simulation, parameter values were chosen to mimic the distribution of the Acute to Chronic Pain Signatures (A2CPS) baseline worst pain scores shown in Figures 2 and S3. Details on the model parameters are provided in Appendix 6.1. This simulation provides an example in which one time point exhibits substantial zero inflation while the other is relatively non-inflated.

In addition, we included a simulation study on one-inflated data in Appendix 6.2, and another using subject-level dispersion in Supplementary Materials (Figure S8). We further assessed the stability of BBME across different parameter settings, including settings corresponding to zero- and one-inflated data, and across different sample sizes (Appendix 7). We also compared the performance of Bayesian and frequentist estimation framework using the same datasets (Appendix 8). Finally, we examined whether BBME might incorrectly interpret random noise as meaningful signal (i.e., false discovery; Appendix 9).

### 2.3 Application to Real Data

We applied BBME to two real-world datasets, one non-inflated and the other zero-inflated pain scores. We do not include a one-inflated application here. Nevertheless, the ability to accommodate both zero- and one-inflation is an important feature of BBME and broadens the potential applications of the model. For example, a high proportion of very high or maximum pain scores may be expected in post-surgery surveys or activity-based assessments in A2CPS.

More generally, this model accommodates bounded outcomes with excess boundary values. Whether the excess occurs at the minimum or maximum depends on how the outcome is defined. For example, some surveys ask subjects to rate their health today on a Visual Analogue Scale (VAS) of 0 to 100, with 100 being best health and 0 being worst imaginable health [12]. This is the reverse of the usual 0-10 pain scale, where higher values indicate worse pain. As a result, zero inflation in one setting may correspond to one-inflation in another, assuming no transformation of the scale is performed. For reference, simulation studies and stability analyses for one-inflated data can be found in Appendix 6.2 and 7.

#### 2.3.1 FM Study

The fibromyalgia (FM) study was a randomized controlled trial of milnacipran versus placebo in 147 FM patients [19]. The study included a 2-week baseline observational period before taking either milnacipran or placebo, followed by a 12-week follow up period. Pain intensity was originally measured on an integer scale between 0 and 132. In the original study, pain scores were divided by 6.6 to create a 0 to 20 scale for comparison with the Gracely Box Scale (GBS), but in our analysis we used the raw 0 to 132 scores. During both the baseline and follow-up periods, subjects reported pain in real time using a Palm-based electronic diary, averaging 3.4 ratings per day at random intervals.

As shown in Figure 5, this dataset contains neither a high proportion of zero scores nor a high proportion of maximum scores (i.e. 132). It therefore serves as an example of fitting the model to non-inflated data.

**Figure 5.**
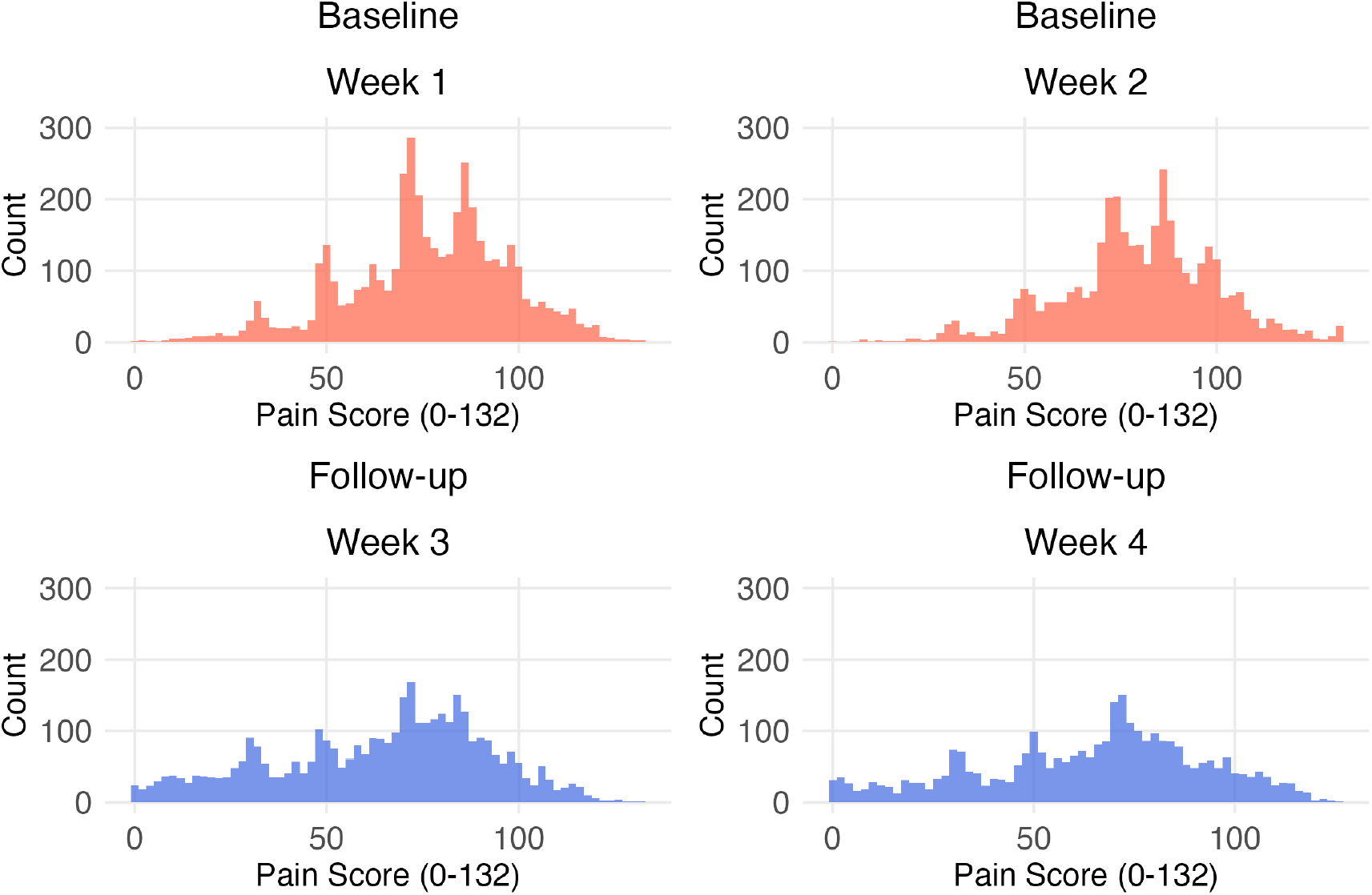
Histogram of pain scores from 147 subjects in the fibromyalgia (FM) study. Pain was recorded on a 0 to 132 scale multiple times per day during the 2-week baseline period and the 12-week follow-up after randomization to milnacipran or placebo. For modeling purposes, we focus on four weeks of data: the 2 baseline weeks (weeks 1-2) and the final two weeks of the follow-up period (weeks 3-4).

For analysis, we fitted a BB model using pain data from the 2-week baseline and the last two weeks of the follow-up period. We grouped the days into four time points: days 0-7 of the baseline as week 1, days 8-14 of the baseline as week 2, days 0-7 of the follow-up as week 3, and days 8-14 of the follow-up as week 4. The resulting dataset consisted of a sample of *I* = 147 subjects, *J* = 4 time points, and *K* = 45 pain assessments per time point (week). Within each time point, between 5 and 45 assessments were completed by each subject. We then fitted the full model with a common dispersion parameter on the dataset, with implementation details in Appendices 3 and 10.1. Our goal was to study differences in pain intensity across weeks, especially between baseline and follow-up, and to characterize how pain changed within each week while accommodating between-subject variation. To this end, we quantified both population-level and subject-specific trajectories and demonstrated how posterior samples can be used to visualize clinically relevant quantities, such as the probability that the observed pain score is above 100.

For comparison, we also fitted an LMM using the R package lme4, with the same hierarchical structure but without a dispersion parameter; details on the LMM specification are in Appendix 10.1.

#### 2.3.2 A2CPS Cuff Pressure Pain

The Acute to Chronic Pain Signatures (A2CPS) program investigates patients before and after surgery to identify predictors of the transition from acute to chronic pain [3,36]. The study focuses on individuals undergoing total knee arthroplasty (TKA) or thoracic surgery. Pain is assessed pre- and post-surgery using multiple approaches, including patient-reported worst surgical-site pain measured on a 0-10 numerical rating scale, cuff pain assessed during functional magnetic resonance imaging (fMRI) and quantitative sensory testing (QST), and movement-related pain measured during functional testing. As of January 2025, pre-surgical baseline data were available for 1043 TKA subjects and 358 thoracic subjects (Release 2.1.0). Baseline worst surgical site pain ratings (Figure 2) contained 2.6% zeros and 8.3% tens in the TKA cohort (little zero- or ten-inflation), compared with 70.7% zeros and 0.8% tens in the thoracic cohort (substantial zero-inflation).

In this study, we focused on cuff pressure pain measured during the fMRI protocol [35]. Pain was assessed on a 0-10 scale in increments of 0.5. Cuff pressure was applied using a Hokanson E20 AG101 Rapid Inflation Cuff System during fMRI sessions. Subjects with high blood pressure, prior lower-limb vascular surgery, lower limb vascular dysfunction, or peripheral neuropathy were excluded from cuff pain testing. For eligible subjects, the cuff was applied to the non-surgical limb in eligible TKA patients and the non-dominant limb on eligible thoracic patients. Before scanning, a baseline cuff pressure between 80-300 mmHg was applied to achieve a target pain rating of 4 out of 10, with 3.5-5 considered an acceptable pain range. We refer to this as the pain4 pressure.

During the first cuff pain scan, denoted CUFF1, the pain4 pressure was maintained for six minutes, after which the cuff was deflated. Subjects reported cuff pain during three intervals: beginning (first two minutes), middle (next two minutes), and end (final two minutes). A second cuff pain scan session, denoted CUFF2, was then conducted using a constant 120 mmHg pressure. This session was discontinued if the subject could not tolerate this pressure level. Subjects again rated cuff pain at the same three intervals (beginning, middle, end). Finally, a resting-state fMRI scan, denoted REST2, was performed, and subjects again rated cuff pain at the same three intervals (beginning, middle, end).

For this study, we focused on pain ratings measured during CUFF1 and REST2 scans. Ratings were doubled and rescaled to integers on a 0-20 scale for consistency. The resulting dataset consisted of a sample of I = 495 subjects, *J* = 2 time points, and *K* = 3 pain assessments per time point. Within each time point, subjects completed between 1 to 3 assessments. The number of completed assessments at each time point can be found in table S5. After data processing, we fitted the full model with common dispersion parameter, adjusting for covariates; implementation details are provided in Appendices 3 and 10.2. As in the FM analysis, we also fitted an LMM using R package lme4 with the same hierarchical structure for comparison, except that the LMM does not include a dispersion parameter; LMM syntax used is in Appendix 10.2.

As shown in Figure 6, the cuff pressure pain dataset contains a high proportion of zero pain ratings, especially during REST2 across both cohorts. These features make the A2CPS Cuff Pressure Pain data a perfect example of where we expect the BBME model more appropriately models the observed data as compared to alternative approaches.

**Figure 6.**
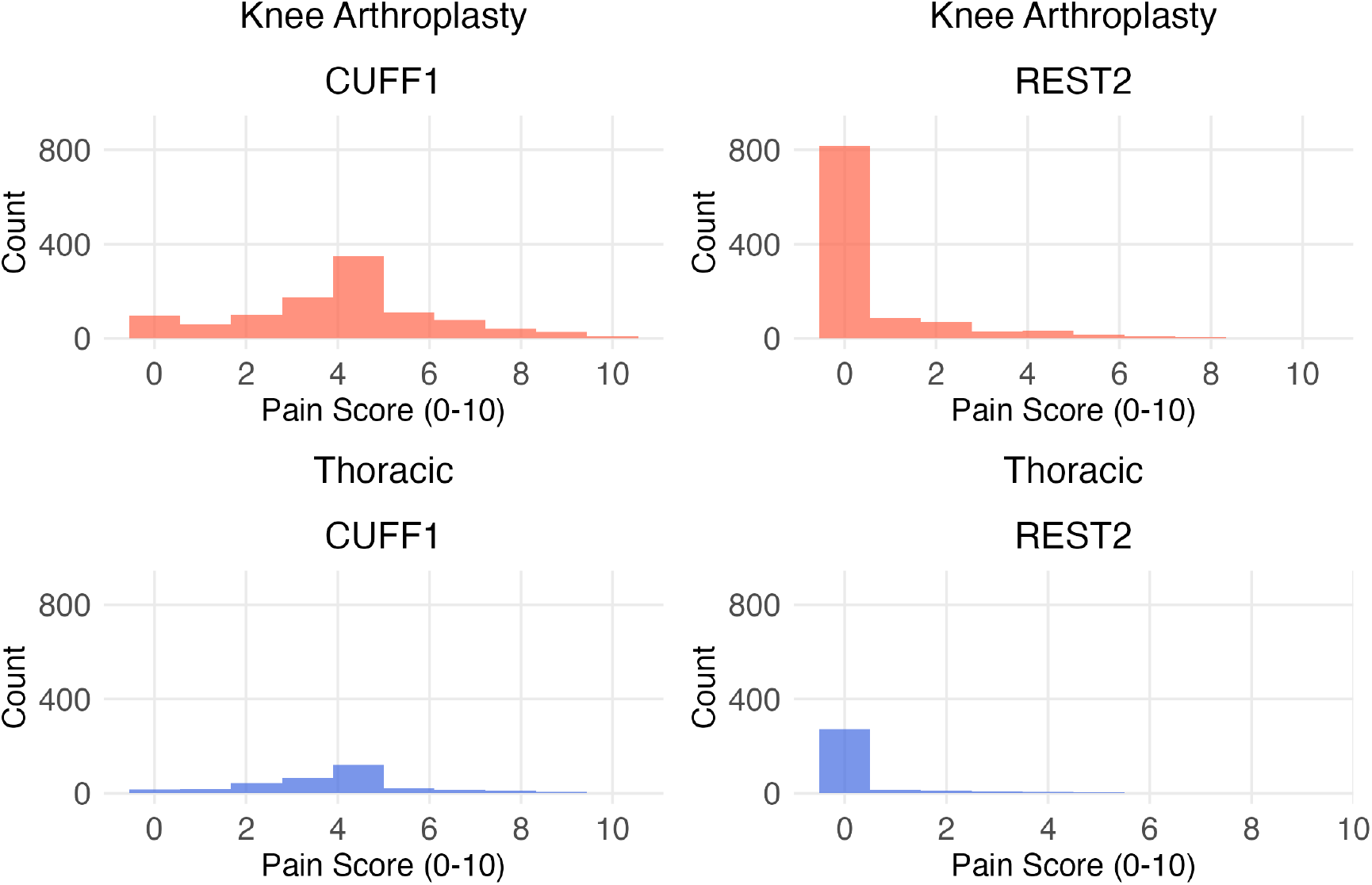
Histogram of cuff pressure pain ratings from 495 subjects in the A2CPS study (385 total knee arthroplasty (TKA) subjects, 110 thoracic subjects). Each subject rated their pain three times under a subject-specific baseline cuff pressure during three 2-minute time intervals of a 6-min fMRI scan (CUFF1). The same procedure was repeated during a resting session (REST2) to obtain three pain scores per scan. The histogram shows the three pain scores per subject at each session, for TKA and thoracic subjects separately.

## 3 Results

### 3.1 Simulation Results

Table 1 summarizes results from the simulations described in Section 2.2. Posterior means for all parameters were close to the true values, and the corresponding 95% credible intervals included the true values in all cases. In addition, the model recovered subject-level parameters, as discussed in detail in Appendix 6.1.

**Table 1:** True and posterior estimates for simulated data. Data were generated for 500 subjects observed at 2 time points. Each subject had a 15% chance having data at only one time point. At each time point, 4-10 pain scores were randomly sampled for each subject. Dispersion varies by timepoint. The model was fit in Stan using 4 parallel chains with 3000 burn-in iterations followed by 6000 sampling iterations, saving every second draw (thinning factor of 2). Effective sample size (ESS) reflects the size of independent posterior samples of each parameter. All parameters converged with 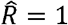.

| Parameter | True | Posterior mean | SD | Posterior median | 2.5% quantile | 97.5% quantile | ESS |
| --- | --- | --- | --- | --- | --- | --- | --- |
| $r_1$ | -3.2 | -3.31 | 0.13 | -3.31 | -3.56 | -3.07 | 1665 |
| $r_2$ | 0.6 | 0.66 | 0.06 | 0.66 | 0.54 | 0.79 | 4812 |
| $w_1$ | 0.0 | 0.02 | 0.04 | 0.02 | -0.05 | 0.09 | 2727 |
| $w_2$ | 0.0 | 0.00 | 0.03 | 0.00 | -0.05 | 0.05 | 4849 |
| $\sigma_{r1}$ | 1.5 | 1.52 | 0.12 | 1.52 | 1.30 | 1.77 | 962 |
| $\sigma_{r2}$ | 1.0 | 1.04 | 0.06 | 1.04 | 0.93 | 1.16 | 2772 |
| $\sigma_{w1}$ | 0.6 | 0.55 | 0.04 | 0.55 | 0.48 | 0.63 | 2117 |
| $\sigma_{w2}$ | 0.5 | 0.53 | 0.03 | 0.52 | 0.48 | 0.58 | 3898 |
| $\rho_{r1,r2}$ | 0.5 | 0.40 | 0.08 | 0.41 | 0.25 | 0.55 | 1558 |
| $\rho_{r1,w1}$ | 0.3 | 0.44 | 0.08 | 0.44 | 0.28 | 0.60 | 1171 |
| $\rho_{r1,w2}$ | 0.0 | 0.13 | 0.08 | 0.13 | -0.03 | 0.27 | 2539 |
| $\rho_{r2,w1}$ | -0.3 | -0.21 | 0.07 | -0.21 | -0.34 | -0.07 | 3131 |
| $\rho_{r2,w2}$ | 0.0 | 0.03 | 0.07 | 0.03 | -0.10 | 0.17 | 4103 |
| $\rho_{w1,w2}$ | 0.0 | 0.09 | 0.07 | 0.09 | -0.04 | 0.22 | 3704 |
| $\phi_1$ | 5.0 | 5.59 | 0.45 | 5.56 | 4.77 | 6.55 | 3185 |
| $\phi_2$ | 10.0 | 10.03 | 0.76 | 9.99 | 8.68 | 11.65 | 4233 |

For comparison, model results for the one-inflation simulation can be found in Appendix 6.2, whereas results for the full model with subject-level dispersion are presented in the Supplementary Material (Table S1, Figures S8-S12).

### 3.2 Application to Real Data

#### 3.2.1 FM Study

##### 3.2.1.1 Population-Level Changes Across Study Weeks

FM analysis results are summarized in Table 2. On average, pain levels were higher during the two-week baseline period than during the two-week follow-up period. Treating each week as a separate timepoint, the posterior means of the fixed intercept were 0.25 (95% CrI: 0.15 to 0.36) in baseline week 1 and 0.46 (95% CrI: 0.36 to 0.57) in baseline week 2, but decreased to −0.16 (95% CrI: −0.33 to 0.02) in follow-up week 3 and −0.09 (95% CrI: −0.27 to 0.09) in follow-up week 4. Thus, there was clearer evidence of higher average pain during the baseline weeks than during the follow-up weeks, where the pain score was on average 74 (out of 132) at the start of week 1 and 81 at the start of week 2 in baseline weeks, and dropped to 61 in follow-up week 3 and 63 in follow-up week 4.

**Table 2:**
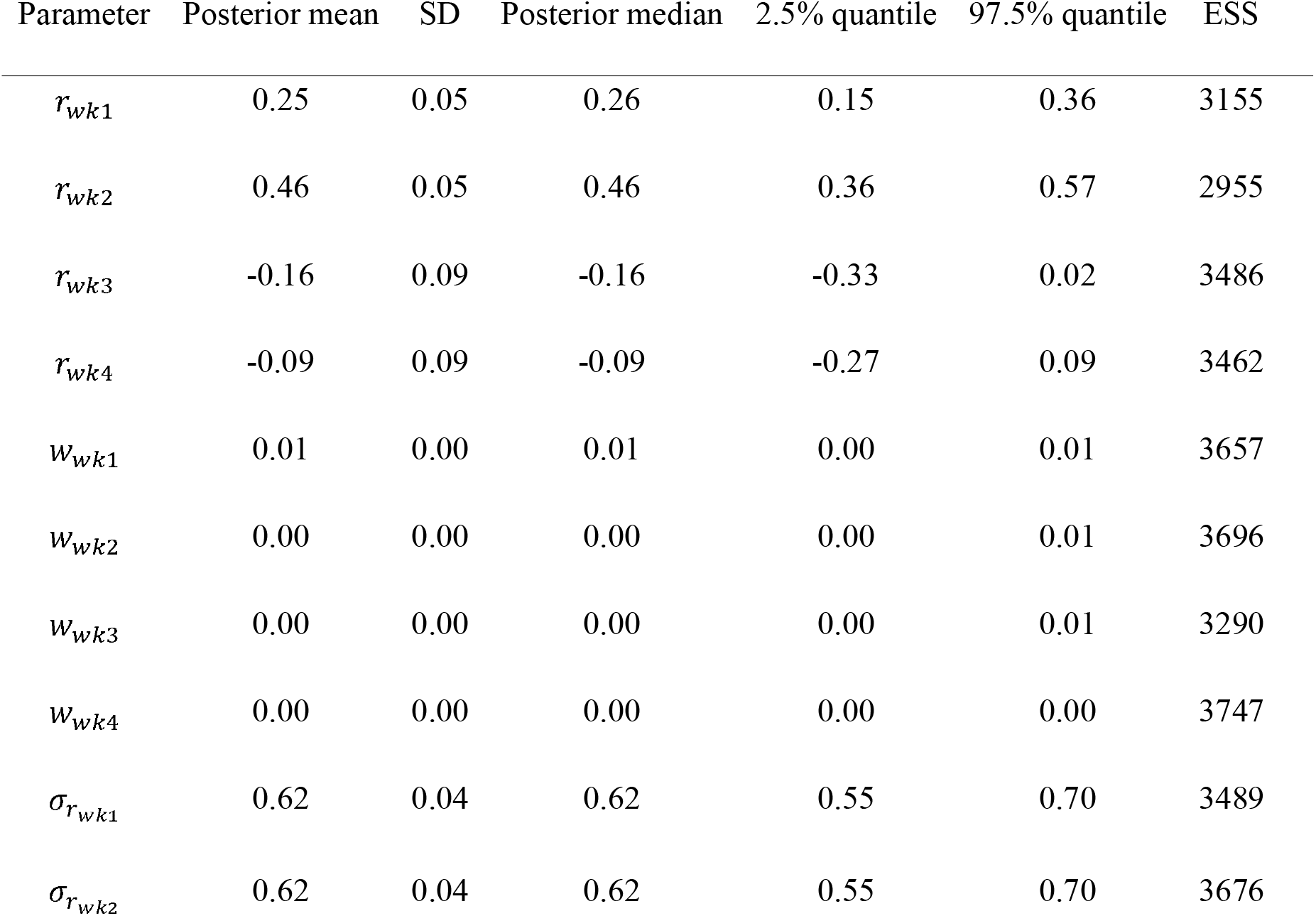
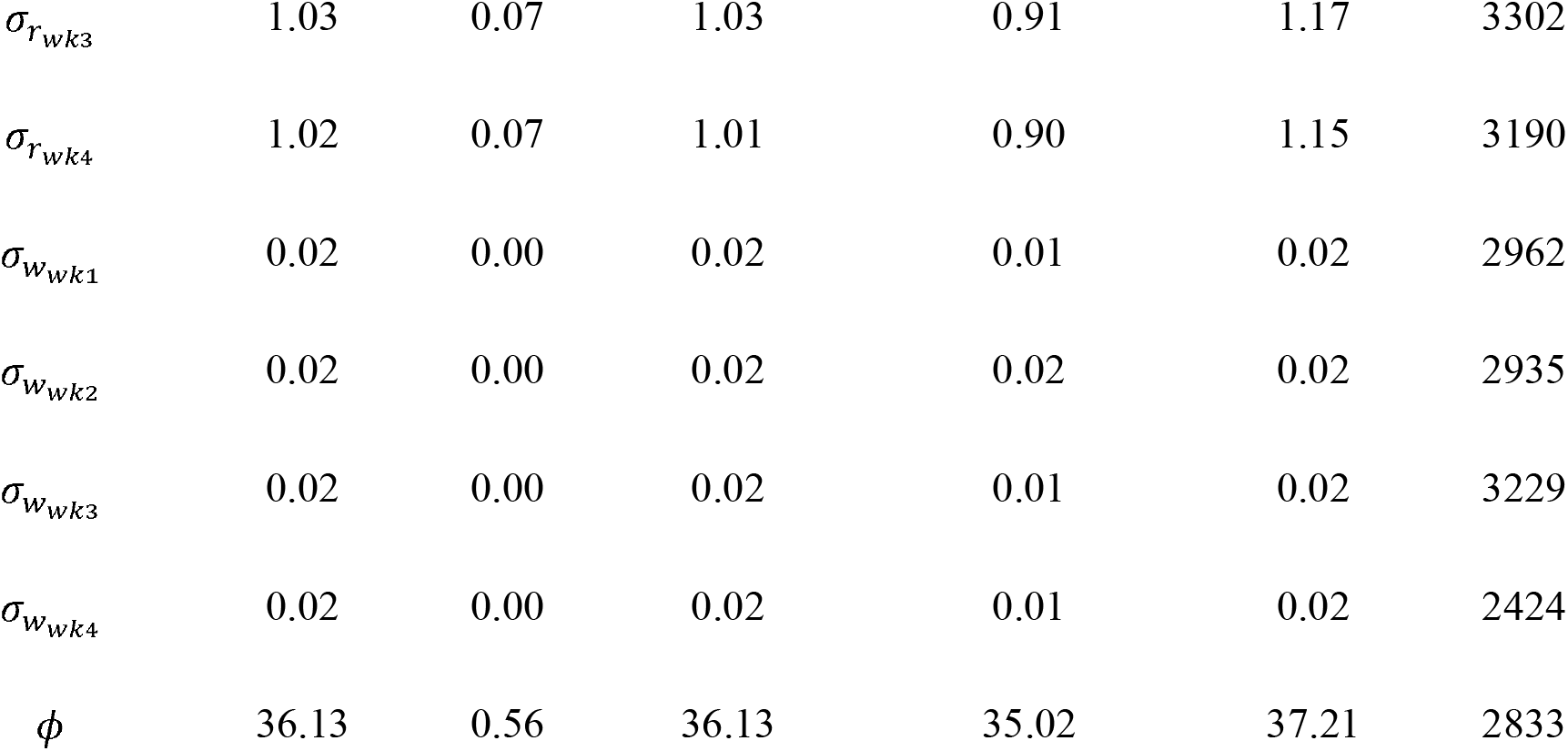
Posterior estimates in the FM study, including mean of random intercepts (*r*), mean of random slopes (*w*), standard deviations of random effects (*σ*), and dispersion (*ϕ*). All parameters converged with 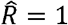. All correlation parameters are shown in the Figure S14(a).

Figure S14(a) shows the correlations among the random effects. Random intercepts for the two baseline weeks were positively correlated, as were the random intercepts at the two follow-up weeks, suggesting that subjects with higher initial pain in the week before also tended to have higher initial pain in the next week. In contrast, the random intercept and random slope of baseline week 1 were negatively correlated, suggesting that subjects with higher initial pain scores tended to experience decreases in pain over the first week. However, since within-week changes was small (for example, posterior mean of the fixed slope in week 1 was 0.01 (95% CrI 0.00 to 0.01)), subjects with high initial level of pain tended to still have high pain at the end of the week. Based on posterior mean random effects, 49.6% of the subjects (73 out of 147) had initial pain higher than average level in baseline week 1; among them, 71.2% (52 out of 73) still had pain higher than average at the end of week 1.

##### 3.2.1.2 Subject-Specific Pain Trajectories

Figure 7 shows the observed and estimated pain trajectories for three subjects: a) a randomly selected subject, b) a subject with low pain, and c) a subject with high pain. These examples illustrate that BBME captures both between- and within-week changes in pain. Predicted trajectories from BBME and LMM were broadly similar and aligned closely with the observed scores.

**Figure 7.**
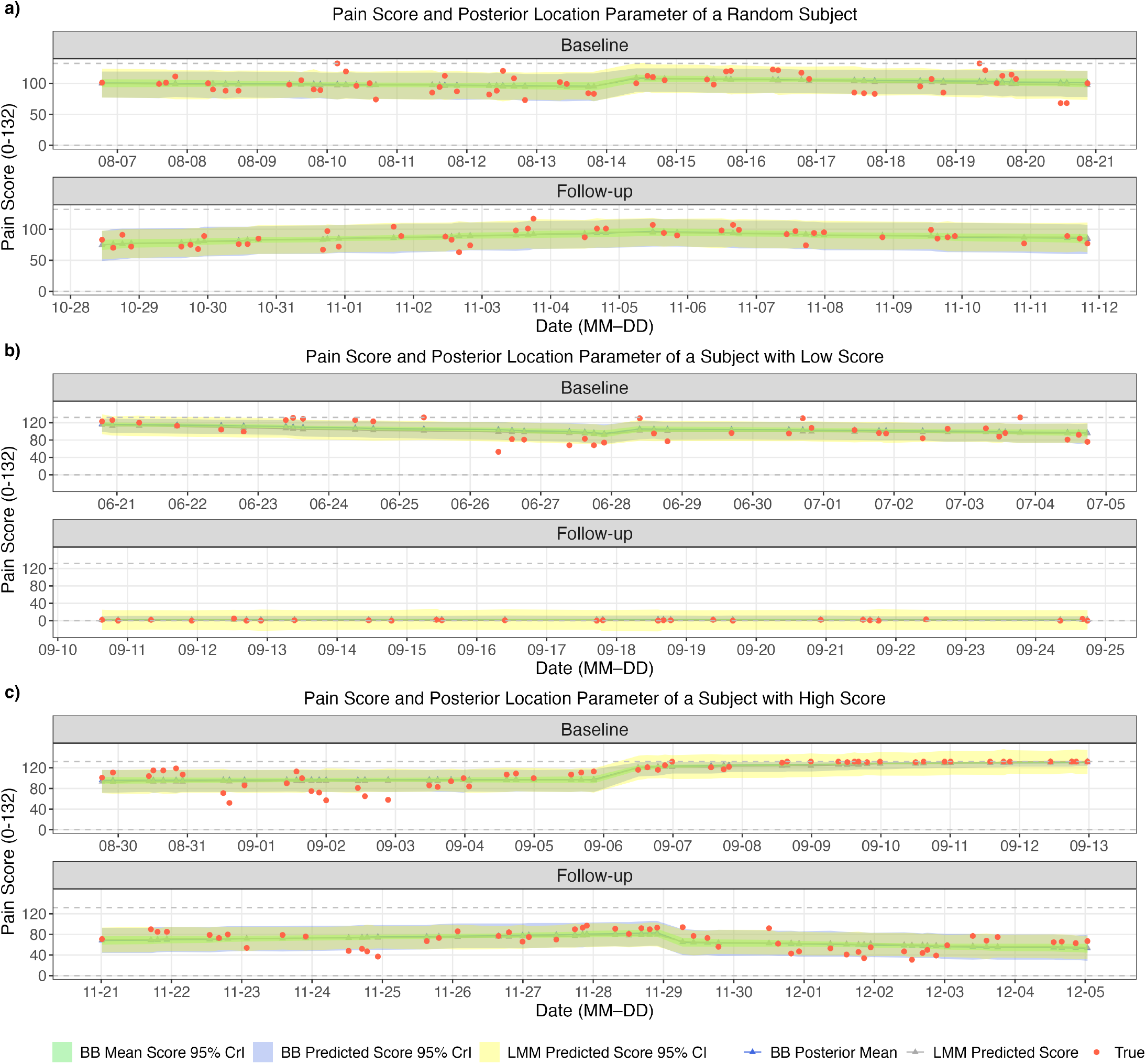
Trajectory of observed pain scores and posterior mean location parameters for a) a random subject in the fibromyalgia (FM) study, b) a subject with low score, and c) a subject with high score. Pain was assessed on a numerical rating scale on 0 to 132 (shown in gray horizontal dash lines). Pain scores were grouped into four time points: two weeks in baseline and two weeks in follow-up. The trajectory captures trends both within and between weeks. The random subject in a) showed a lower average pain score in the follow-up weeks compared to the baseline weeks.

For the randomly selected subject, uncertainty intervals from BBME and LMM were similar (Figure 7a). However, for subjects with consistently low or high pain, LMM produced invalid uncertainty intervals when the predicted scores approached the boundaries of the scale (Figure 7b, c). Specifically, the upper or lower bounds of some LMM intervals fell outside the valid range of 0 to 132. In contrast, BBME always respected the bounded scale. While most LMM point predictions remained within range, approximately 10% of intervals were outside the allowable range during each follow-up week, compared with about 3% during each baseline week. These results suggest that even in a dataset that is approximately non-inflated, BBME provides more reliable subject-level uncertainty quantification near the boundaries of the scale.

##### 3.2.1.3 Comparison with Linear Mixed-Effects Models

In the FM dataset, where pain scores were approximately non-inflated and symmetric (Figure 5), BBME achieved a similar model fit as LMM, with an in-sample log-likelihood of −56,979.52 evaluated at posterior mean location and dispersion parameters, compared to in-sample log-likelihood −58,711.51 for the LMM fit using lme4 package; however, these comparisons do not account for differences in model complexity. The predicted pain scores were also similar between the two models (Figure S15). Although the parameter estimates were not directly comparable because the two models were formulated differently, they showed similar overall trends (Table S3). For example, in LMM, the mean intercept in week 1 was 73.89 (SE = 1.54), higher than the mean intercept in week 3 (63.14, SE = 2.37), indicating that average pain scores were higher during baseline than during follow-up. The correlation between the intercept and slope in week 1 was −0.38, consistent with the negative association seen in BBME (Figure S14).

When treatment assignment was added as a covariate, both the LMM and BBME showed no clear evidence of treatment difference in mean pain scores between the treatment and placebo groups (Table S6-7), although the parameters differ in interpretation under the two modeling frameworks. Overall, despite violating the usual distributional assumptions, LMM produced similar predicted outcomes and led to broadly consistent conclusions with those form BBME for the FM data. This suggests that for relatively non-inflated pain data, standard and BBME-based analyses may lead to similar substantive conclusions, although BBME remains better aligned with the bounded nature of the outcome.

##### 3.2.1.4 Posterior Probabilities for Clinically Relevant Pain Thresholds

The BBME framework also allows direct probability statements about clinically relevant events. For example, on the 0 to 132 pain scale, the probability of having pain score above 100 at the given assessment can be calculated by computing the proportion of posterior predictive pain scores larger than that threshold. The posterior probability that a randomly selected subject’s pain score exceeded 100 at the last baseline assessment was 55.6%, whereas the posterior probability that the score exceeded 100 at the first test in follow-up week 4 was 38.5%. Figure 8 shows the full trajectory of this probability over time. These results illustrate how posterior samples from BBME can be used to answer clinically interpretable questions about whether a patient is likely to exceed a threshold of concern.

**Figure 8.**
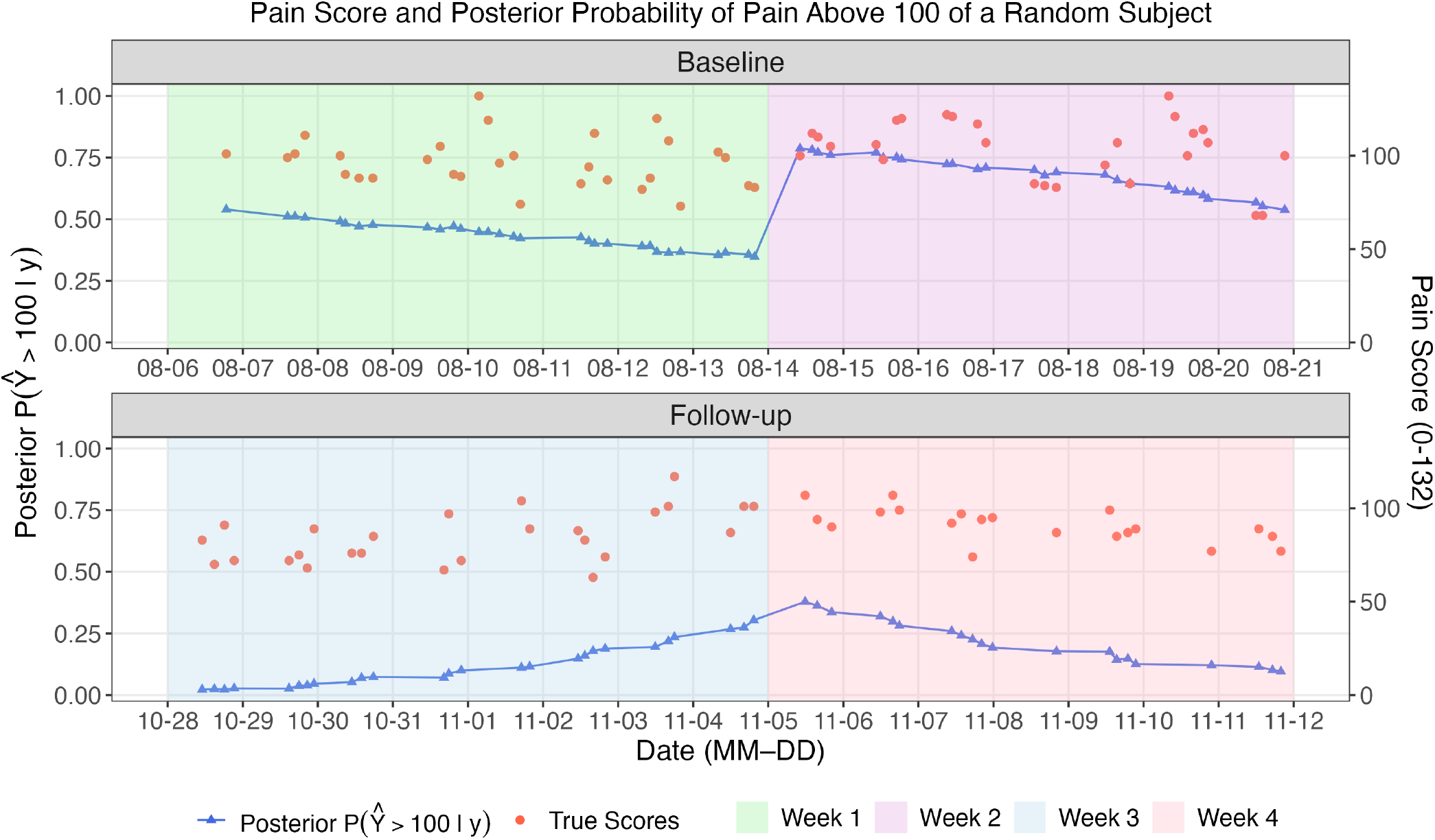
Trajectory of observed pain scores and posterior probability of pain score higher than 100 for a randomly selected subject in the FM study. This subject showed a lower probability of having pain score above 100 in the follow-up weeks compared to in baseline weeks.

#### 3.2.2 A2CPS Cuff Pressure Pain

##### 3.2.2.1 Population-Level Differences Between CUFF1 and REST2

Results for the cuff-pain analysis are summarized in Table 3. On average, pain during CUFF1 was higher than during REST2. The mean random intercept in CUFF1 was −0.86 (95% CrI −1.03 to −0.71), corresponding to a more symmetric distribution of pain scores. In contrast, the mean of random intercepts in REST2 was −6.65 (95% CrI −7.50 to −5.91), corresponding to average pain score at 0.02 (on a scale of 0 to 20). The mean random slope was positive in CUFF1 but negative in REST2, indicating that pain tended to increase during the 6-minute cuff session and gradually decrease during the 6-minute rest session. The population level mean pain scores are shown in Figure S17.

**Table 3:** Posterior estimates for cuff pressure pain in the A2CPS study, including mean of random intercepts (*r*), mean of random slopes (*w*), standard deviations of random effects (*σ*), and dispersion (*ϕ*). All model parameters converged with 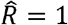.

| Parameter | Posterior mean | SD | Posterior median | 2.5% quantile | 97.5% quantile | ESS |
| --- | --- | --- | --- | --- | --- | --- |
| $r_{\text{CUFF1}}$ | -0.86 | 0.08 | -0.86 | -1.03 | -0.71 | 1093 |
| $r_{\text{REST2}}$ | -6.65 | 0.41 | -6.63 | -7.50 | -5.91 | 1000 |
| $w_{\text{CUFF1}}$ | 0.33 | 0.03 | 0.33 | 0.28 | 0.39 | 2474 |
| $w_{\text{REST2}}$ | -0.65 | 0.25 | -0.64 | -1.16 | -0.17 | 297 |
| $\sigma_{r_{\text{CUFF1}}}$ | 0.99 | 0.05 | 0.99 | 0.90 | 1.09 | 3715 |
| $\sigma_{r_{\text{REST2}}}$ | 4.06 | 0.34 | 4.05 | 3.43 | 4.76 | 921 |
| $\sigma_{w_{\text{CUFF1}}}$ | 0.45 | 0.03 | 0.45 | 0.39 | 0.50 | 3239 |
| $\sigma_{w_{\text{REST2}}}$ | 0.75 | 0.13 | 0.74 | 0.54 | 1.04 | 333 |
| $\rho_{r_{\text{CUFF1}}, r_{\text{REST2}}}$ | 0.27 | 0.07 | 0.27 | 0.12 | 0.40 | 4086 |
| $\rho_{r_{\text{CUFF1}}, w_{\text{CUFF1}}}$ | -0.25 | 0.07 | -0.25 | -0.39 | -0.11 | 3297 |
| $\rho_{r_{\text{CUFF1}}, w_{\text{REST2}}}$ | -0.21 | 0.12 | -0.21 | -0.44 | 0.04 | 1200 |
| $\rho_{r_{\text{REST2}}, w_{\text{CUFF1}}}$ | 0.07 | 0.08 | 0.07 | -0.09 | 0.22 | 4172 |
| $\rho_{r_{\text{REST2}}, w_{\text{REST2}}}$ | 0.45 | 0.19 | 0.47 | 0.00 | 0.74 | 328 |
| $\rho_{w_{\text{CUFF1}}, w_{\text{REST2}}}$ | 0.28 | 0.11 | 0.28 | 0.04 | 0.49 | 1660 |
| $\beta_{TKA}$ | 0.02 | 0.08 | 0.01 | -0.14 | 0.21 | 751 |
| $\phi$ | 1004.11 | 8350.57 | 284.58 | 101.37 | 4514.07 | 6276 |

We also observed a mild positive correlation of 0.27 (95% CrI 0.12 to 0.40) between the random intercepts of CUFF1 and REST2 location parameters, suggesting that subjects who reported a higher pain score at the start of the cuff session were more likely to report higher pain at the start of the rest session. The estimated common dispersion parameter was low in this dataset. The posterior mean covariate effect for TKA subjects (*β*_TKA_) was 0.02 (95% CrI −0.14 to 0.21), implying an odds ratio of 1.02 (95% CrI 0.87 to 1.23) for the location parameter compared to thoracic subjects. This suggests no meaningful difference in pain scores between the two cohorts.

##### 3.2.2.2 Subject-Specific Pain Trajectories and Interval Validity

Random effects in both BBME and LMM allow estimation of subject-specific pain trajectories. However, the two models differed substantially in the validity of predicted scores and uncertainty intervals near the lower boundary of the scale. As shown in Figure 9, when the observed score was 0, the 95% credible intervals of BBME-predicted scores were much tighter, whereas the 95% prediction intervals from the LMM were considerably wider and sometimes extended below zero, which is not a valid value on the 0-20 pain scale.

**Figure 9.**
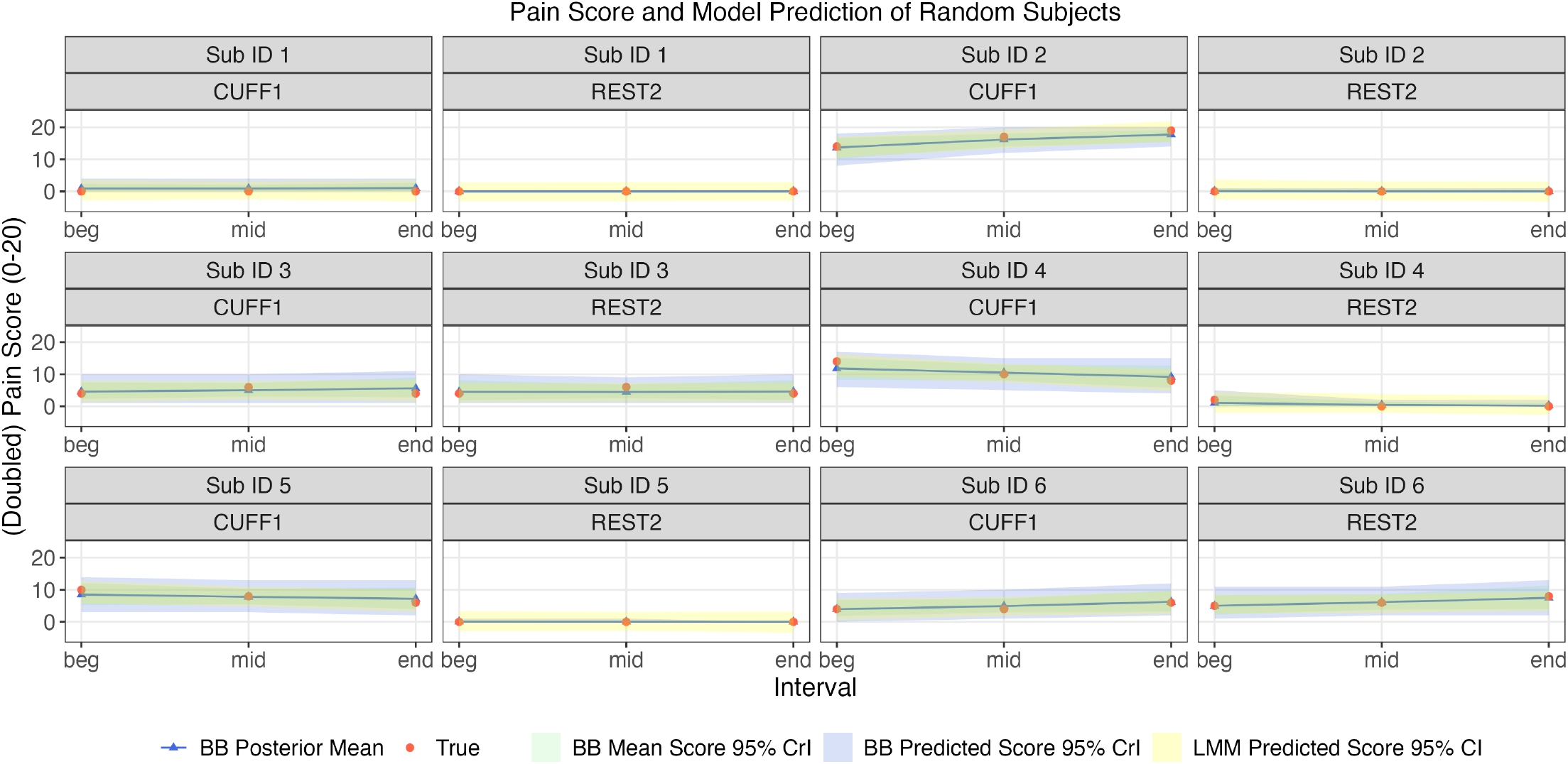
Trajectory of pain scores of six randomly selected subjects. Temporary ID 1 to 6 were assigned to the subjects in A2CPS Cuff Pain dataset. The 95% prediction intervals for predicted score of LMMs were computed using predictInterval() function by merTools R package [**?**]. 95% intervals by LMMs were much wider for true score equals to zero, and extended below zero in some subjects.

Using the definition of an invalid predicted score as one below 0 or above 20, and an invalid interval as one with either endpoint outside [0, 20], all predicted scores and intervals from BBME remained valid for CUFF1. However, when using LMM, for thoracic subjects during REST2, 22.6%, 19.8%, and 36.8% of predicted scores were invalid at the beginning, mid, and end of the session, respectively. In addition, 88.7%, 88.7%, and 90.6% of the corresponding LMM intervals were invalid at those three locations in the REST session. Invalid scores and intervals were also observed for TKA subjects (Supplementary Table S5).

These results show that LMM can produce implausible subject-level predictions and uncertainty intervals when the outcome lies near the boundary. In contrast, BBME always respects the bounded outcome scale.

##### 3.2.2.3 Comparison with Linear Mixed-Effects Model Parameters

For this zero-inflated dataset, BBME achieved a log-likelihood of −3,106 evaluated at posterior mean location and dispersion parameters, compared to log-likelihood −6,408 for the LMM, indicating a substantially greater improvement in model fit than observed in the FM data.

Despite the poorer fit of the LMM, its parameter estimates showed trends broadly consistent with those from BBME (Table S4). For example, the correlation between the intercepts at CUFF1 and REST2 was 0.30 (Table S4), indicating a mild positive association, similar to that estimated from BBME. The mean intercept at CUFF1 was 6.3 (95% CI: 5.76 to 6.84), in contrast to the mean intercept of 0.89 (95% CI: 0.41 to 1.37) in REST2, which also showed that the initial pain score at CUFF1 was higher than REST2 on average. TKA subjects reported 0.44 (95% CI: −0.08 to 0.96) higher pain than thoracic subjects on average, showing no significant differences between the two cohorts.

Although both models led to similar qualitative conclusions, their parameter estimates differ in interpretation. In BBME, *r*_*cuff*1_ = −0.86 implied that the expected pain levels for thoracic subjects at the start of CUFF1 was expit(−0.86) = 5.95 on the 0-20 scale. In contrast, *r*_*cuff*1_ = 6.3 in LMM represented that thoracic subjects on average reported pain score of 6.3 at the start of CUFF1. Similarly, *w*_*cuff*1_ = 0.33 implied that the odds of the location parameter at mid CUFF1 were exp(0.33) = 1.39 times the odds at the beginning of CUFF1, whereas *w*_*cuff*1_ = 1.39 in the LMM meant that the predicted pain score increased by 1.39 points on average from the beginning to the middle of CUFF1.

Overall, BBME and LMM produced parameter estimates with similar directional trends, but BBME provided a substantially better fit and a more appropriate framework for bounded, zero-inflated pain scores.

##### 3.2.2.4 Recovery of the Observed Score Distribution

To further compare the two models, we generated two resampled datasets with *I* = 495 subjects, *J* = 2 time points, and *K* = 3 pain assessments per time point: one using the posterior mean parameters from the BBME model with common dispersion (i.e. *r, w, σ, ρ, ϕ*), and the other using the parameter estimates from the LMM. Because the resampled datasets did not include missingness, the effective sample size was larger than in the observed data.

As shown in Figure 10, the dataset resampled from BBME closely resembled the true cuff pain data. For example, in one such dataset, the proportions of zeros at the beginning, middle, and end of REST2 session for TKA subjects were 72.5%, 75.6%, and 77.9%, respectively, close to the corresponding proportions of 74.2%, 73.6%, and 75.8% in the observed data.

**Figure 10.**
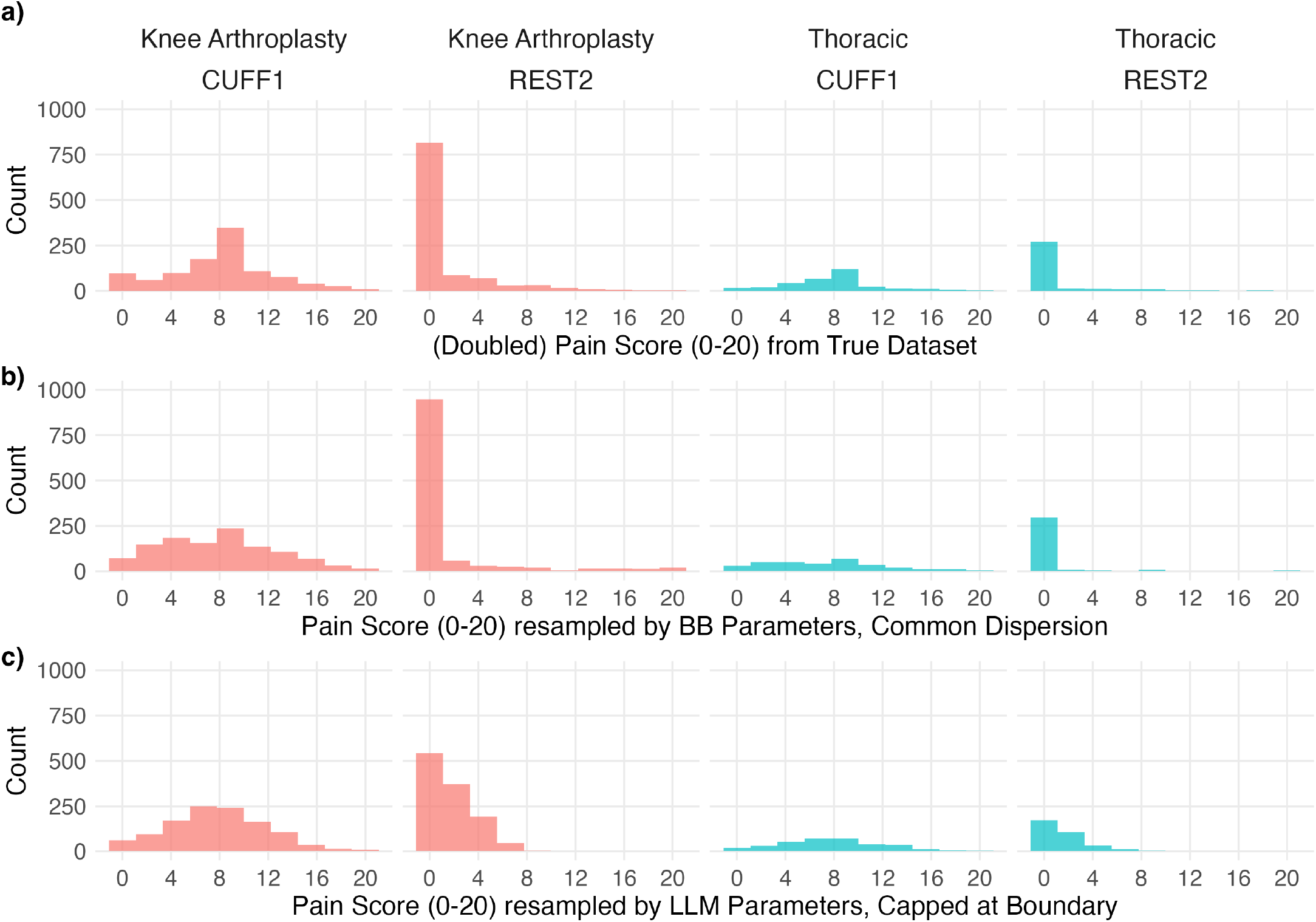
Distribution of a) true scores in observed A2CPS Cuff Pain dataset, and b) one randomly resampled dataset using BBME-estimated population parameters, and c) one randomly resampled dataset using LMM-estimated population parameters, with scores capped at [0, 20].

Because the LMM assumes an unbounded continuous outcome, the LMM-resampled dataset included invalid values below 0 and above 20 (Figure S18). We therefore truncated values below 0 to 0, and values above 20 to 20. Even after truncation, resampled LMM data did not resemble the observed distribution. For example, the proportions of zeros at the beginning, middle, and end of REST2 for TKA subjects were 27.8%, 25.7%, and 29.4% respectively, much lower than those observed in the true dataset. Thus, although the LMM produced parameter estimates with qualitatively similar trends, it failed to capture the strong zero inflation and overall score distribution.

##### 3.2.2.5 Posterior Probabilities for Clinically Relevant Comparisons

The BBME framework also supports direct probability statements about clinically relevant events. Using the same approach as in the FM study, we calculated posterior probabilities for clinically meaningful events. For example, we estimated the posterior probability that a subject’s pain score was below 6 on the scale of 0 to 20 for several randomly sampled subjects (Figure 11(a)). Temporary IDs 1 to 6 were assigned to these subjects. Figure 11(a) shows, for example, that subject 4 had a slightly higher posterior probability of pain below 6 at the end of CUFF1 than at the beginning. In contrast, subject 6 showed a lower probability of pain below 6 at the end of CUFF1 compared to that at the beginning of CUFF1.

**Figure 11.**
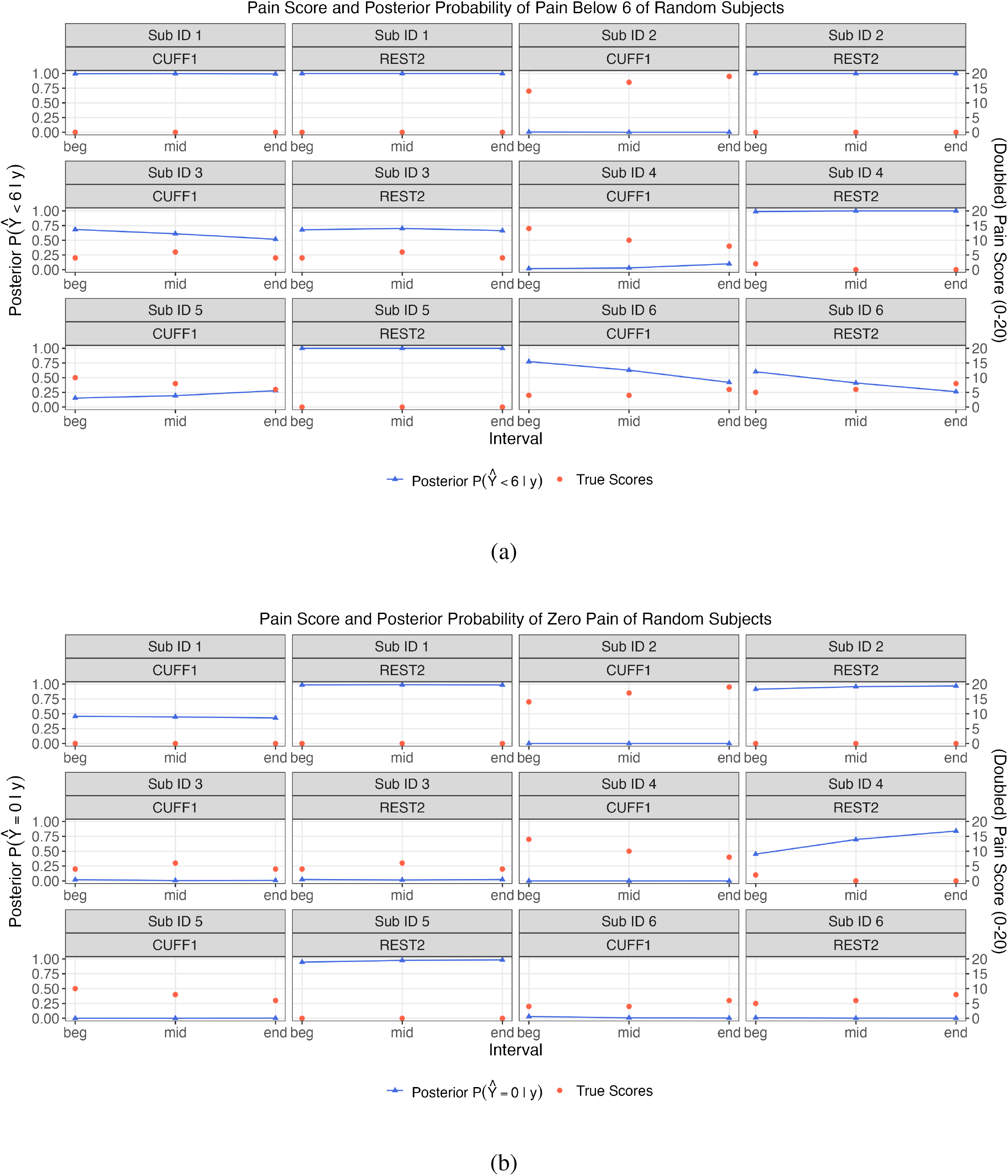
Trajectory of pain scores and posterior probability of score a) below 6 and b) being 0 of randomly sampled subjects in A2CPS Cuff Pain dataset.

A similar procedure can be used to estimate the probability of zero pain (Figure 11(b)). This quantity cannot be directly computed under LMM, which assumes a continuous distribution on pain scores and therefore cannot provide probability estimates to any exact value, such as zero.

In addition, posterior samples allow direct comparison of the two sessions. For example, for subject 1, the posterior probability that pain at the end of CUFF1 was higher than pain at the end of REST2 was 56.6%, whereas the corresponding probability for subject 5 was 99.7%. This probability is calculated as the proportion of posterior samples, among the 8000 saved iterations, in which the predicted score at the end of CUFF1 is greater than the predicted score at the end of REST2.

These examples illustrate how posterior samples from BBME can be used not only for estimation, but also to answer clinically interpretable questions about the likelihood of specific pain states or changes over time both for each subject and the whole sample.

#### 3.2.3 Comparison of the models on two real data

Looking at the two real-data applications, LMMs can provide reasonable estimates on the mean structure of pain (Table S3-4, S6-7, Figure S14). However, they are mis-specified for bounded-integer outcomes. Increasingly, researchers are interested in not just average values. This mis-specification limits the ability of LMMs to accurately characterize other aspects of pain score distribution.

First, LMMs assume a continuous distribution on pain and therefore do not compute probabilities of specific values, for example Pr(*Pain* = 0), which is often of primary clinical interest. In contrast, BBME is suitable for discrete outcomes and can naturally provide these probabilities (Figure 11). Bayesian framework further facilitates estimation of these probabilities, such as zero pain, mild pain, or even comparing pain across time points (Figures 8 and 11).

Second, distribution of bounded integer outcomes is not only characterized by the mean but also the variance. In binomial-type outcomes, variance is a function of mean, and additional variability can arise from correlation among underlying Bernoulli components (e.g. 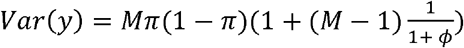 for BB outcome *y*, in Appendix 1), leading to overdispersion and the subsequent increased probability mass at boundaries. In contrast, LMMs impose a variance structure that is not linked to mean and thus does not account for the intrinsic mean-variance relationship and the extra variability induced by overdispersion. As a result, they may mischaracterize variability estimates. This is demonstrated by the much tighter uncertainty estimates of subject-specific trajectories by BBME in Figures 7 and 9.

Third, BBME is more suitable for studying the whole distribution of pain score. The binomial-based model with an additional dispersion parameter takes care of both the bounded scale of the outcome and the inflation of boundary values. This is evident in resampled datasets given population estimates (Figure 10). Moreover, LMM-based resampling or prediction may produce values outside valid range, potentially requiring truncation and distorting inference (Figures 7, 9, and 10).

Together, these limitations indicate that LMMs are not well suited for capturing the distributional characteristics of bounded discrete outcomes, even though they can approximate the mean well.

## 4 Discussion

This work introduces a Bayesian beta-binomial mixed-effects (BBME) model for longitudinal pain scores measured on bounded integer scales (e.g., 0-10). The model treats each score as arising from a beta-binomial process, with a latent probability modeled by a beta distribution and the observed score modeled by a binomial distribution conditional on that probability. This formulation naturally accommodates overdispersion and boundary behavior (excess 0s and maximum scores) without separate zero- or one-inflation components.

BBME is appropriate for pain scales used in clinical research. Numerical rating scales are bounded and discrete, yet are often analyzed with Gaussian models such as linear mixed-effects models. That mismatch can distort inference, overestimate uncertainty for boundary scores, and obscure meaningful features. By contrast, BBME respects the bounded-integer nature of pain scores and provides subject-specific predictions via random effects. By modeling random effects in the mean component and allowing correlation across time points, BBME addresses clinically relevant questions such as whether average pain differs from baseline to follow-up, whether patients with higher baseline pain recover differently, and whether reported pain scores tend to change over time even if the mean pain remains unchanged.

The Bayesian formulation produces full posterior distributions for all parameters and predictive quantities. This allows coherent uncertainty quantification, principled pooling of information across subjects and time, and probability statements on clinically interpretable questions (Figures 8 and 11). Such statements are difficult to obtain from standard Gaussian models, but follow directly from the Bayesian BBME framework.

Simulation studies showed that BBME recovered population-level and subject-level quantities with near-nominal coverage under realistic sample sizes. Precision improved as either the number of subjects or repeated assessments increased, offering guidance for study design (Appendix 7). Applications to both inflated and non-inflated datasets suggested that BBME could serve as a default model for bounded-integer pain outcomes. Moreover, in contexts where clinically meaningful pain cutoffs exist, BBME allows for estimation and inference on thresholded scores while avoiding information loss associated with thresholding the data before modelling.

### Clinical relevance

From a clinical perspective, zero inflation occurs when many subjects report no pain, whereas one inflation occurs when many report maximal pain. Zero-inflation may be expected when subjects are not under active pain-inducing conditions, such as resting pain among pre-surgery thoracic patients, control subjects before quantitative sensory testing, or general population surveys. In a nationally representative US EQ-5D survey, approximately half of respondents reported no pain [12].

Maximum pain is more likely in severe clinical settings, such as early post-surgical EMA or among subjects presenting for ambulatory surgery after trauma. A study of chronic pain patients undergoing nononcologic surgery reported mean pre-operative worst pain ratings above 9 on a 0-10 scale, with similarly high values post-operatively [2]. A study of knee replacement patients reported that about 30% experienced the most severe pain category pre-operatively [32]. However, pain-control protocols may reduce the frequency of maximum scores.

Datasets containing both zeros and maximum scores can arise in more complex settings, such as long EMA follow-up after surgery or studies involving heterogeneous populations that include healthy controls and patients with pain conditions. For example, in a study of orthopaedic foot and ankle surgery, a majority of subjects reported zero pain pre-operatively and at 6-weeks post-operatively, whereas maximum pain scores were more common 3-days post-operatively [9]. Here, a single flexible model is particularly useful.

### Comparison with alternative approaches

Several alternative approaches are used in the pain literature. LMMs preserve the numerical scale but assume unbounded continuous outcomes with constant variance [8,22], making them poorly suited to zero/one-inflation and heteroskedasticity. Ordinal models respect discreteness but treat scores as ordered categories with thresholds [11,38], which can be difficult when many observations occur at the boundaries. Multinomial mixed models could avoid thresholding, but they require more parameters than BBME and may be impractical for more densely measured pain ratings (e.g. 0 - 132).

Mixture models are appropriate when extreme values arise from distinct data-generating processes (e.g., structural zeros). However, they are justified only when data truly arise from two distinct generating processes [41]. Otherwise, a large number of zeros may simply reflect a low mean rather than a structural-zero state. In many pain settings, it is more natural to view pain as lying on a continuum, with zero representing the lower bound of the same underlying process rather than a separate biological state. Within BBME, the latent probability *p* ∈ (0, 1) can be interpreted as a propensity for reporting any pain score on the scale. When *p* is near zero, the probability of reporting zero pain increases, and when *p* is near one, the probability of reporting maximum pain increases.

This interpretation fits contemporary biopsychosocial theories of pain, which conceptualize pain as a graded, dynamically modulated experience shaped by sensory, affective, cognitive, and contextual influences. The within-subject variability observed in EMA data further supports this view, as subjects frequently move between zero and nonzero ratings over time. Under this framework, boundary clustering can arise naturally from heterogeneity in latent pain.

Mixture models can be difficult to interpret when covariates affect model components differentially [24]. This problem has appeared in pain studies, where coefficients are reported separately for zero-inflation and count components, complicating interpretation [18,40]. The problem would be further exacerbated if an additional mixture component was introduced to handle maximum pain.

For these reasons, mixture models may be unnecessary when overdispersion can adequately account for extreme values. That said, BBME is not intended to resolve whether pain is best understood as a single continuum or as a mixture of “pain” and “no pain” states. Zero-inflated models remain appropriate when there is genuine reason to believe that both structural and sampling zeros are present. Our goal is to present BBME as a flexible framework that accommodates non-inflated, zero-and/or-one-inflated data.

### Real-data applications

In two real-data applications, BBME compared favorably with LMM. At the subject level, BBME generated valid predicted scores and uncertainty intervals that respected the bounded scale, whereas LMM intervals were wider and sometimes extended below zero. For the zero-inflated cuff-pain dataset [3,35,36], BBME better reproduced the observed distribution of pain scores, especially the large proportion of zeros. BBME reported probabilities of pain events while continuous models were limited in this aspect. Taken together, these applications indicate the utility of BBME on capturing the full distributional characteristics of bounded discrete outcomes.

### Practical considerations, strengths, and limitations

BBME is appropriate when the outcome is a bounded integer, there is evidence of excess boundary values or heterogeneity across the outcome, and subject-specific predictions are of interest. Because it performs well with non-inflated data, analysts need not switch models based on the outcome distribution. In practice, we recommend starting with a logit link for the mean, random intercepts across time points, and random slopes, and a common dispersion, with sensitivity analyses that allow dispersion to vary by subject or time point if substantial heterogeneity is suspected (Appendix 2). Weakly informative priors on fixed effects and hierarchical priors on standard deviations are often sufficient, while horseshoe priors may help when the number of covariates is large (Appendix 4).

Model fitting should include convergence diagnostics (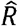, trace plots, e.g. Figure S4, S9), posterior predictive checks, and information criteria such as LOOIC [39]. In our simulation, Hamiltonian Monte Carlo using either Stan or brms performed comparably well [6,7], whereas glmmTMB [5] was faster but under-covered some parameters. Stan generally scaled better in runtime as the number of assessments increased, although at higher memory cost, while brms used relatively stable RAM at the cost of longer runtime (Figure A7).

#### BBME has several strengths

It aligns with the measurement scale, generates boundary mass without structural mixtures, accommodates irregular EMA data, supports subject-level prediction, and performed well across multiple inflation regimes. Clinically, it also supports subject-specific trajectory estimation, uncertainty intervals, and posterior probability statements about meaningful changes in pain over time.

The limitations are also important. The logit link can be difficult to estimate when the true location parameter is close to 0 or 1, though the impact on predicted probabilities was limited in our simulations (Figure S5-6, S11-12). Alternative links may help mitigate this problem. More complex dispersion structures may be needed when heterogeneity is strong, at the cost of additional computational complexity and convergence monitoring. The current model also treats repeated assessments within a time point as conditionally independent given random effects and covariates. If stronger serial dependence is present, autoregressive terms may be necessary. Finally, we treated missingness as ignorable. In the EMA setting, compliance is often informative, so joint modeling of missingness may be required if missing-not-at-random mechanisms are plausible. Future work should address model selection among BBME variants, shrinkage-based feature selection for high-dimensional covariates, and dependence structures beyond a linear within-time trend.

## Conclusion

The beta-binomial mixed-effects model offers a flexible, scale-respecting, unified framework for analyzing patient-reported pain on bounded-integer scales, especially in EMA and multi-phase study designs. It captures extreme values and overdispersion without structural mixtures, delivers subject-level inference, and provides coherent uncertainty quantification through Bayesian posterior summaries. We view BBME as a default for longitudinal pain outcomes and a foundation for specialized models when needed.

## Acknowledgements

Data were provided [in part] by the A2CPS Consortium funded by the National Institutes of Health (NIH) Common Fund, which is managed by the Office of the Director (OD)/ Office of Strategic Coordination (OSC). Consortium components and their associated funding sources include Clinical Coordinating Center (U24NS112873), Data Integration and Resource Center (U54DA049110), Omics Data Generation Centers (U54DA049116, U54DA049115, U54DA049113), Multi-site Clinical Center 1 (MCC1) (UM1NS112874), and Multi-site Clinical Center 2 (MCC2) (UM1NS118922).

Code to reproduce the data from the simulation study is available upon request to the corresponding author Yanxi Liu. A2CPS data can be accessed on https://nda.nih.gov/edit_collection.html?id=5121. Data on Fibromyalgia study is available upon request to authors Daniel Clauw and Richard E. Harris.

The A2CPS Consortium includes: David Chesla, Danneka Cooper, Mark Delano, Nina Duong, Geoffrey Lam, John Mahajan, Todd Mulderink, Opal Tafe, Amber (DeGraaf) White, Brian Winner, Denise Wittenbach, Yong Zhou, James Ford, Sydney Shohan, Michael Sun, Tor Wager, Katerina Zorina-Lichtenwalter, Briha Ansari, Brian Caffo, Vince Calhoun, Ciprian Crainiceanu, Luda Diatchenko, Farzad Farahani, Kara (Schoenberg) Friedman, Adi Gherman, Andre Hackman, Kasper Hansen, Khaled Hasan, Hongkai Ji, Micah Johnson, Martin Lindquist, Marc Parisien, Ingo Ruczinski, Patrick Sadil, Stephani Sutherland, Margaret Taub, Scott Zeger, Erik Westlund, Seth Berke, Yanxi Liu, Bella Satpathy-Horton, Zheng Ren, Raed Alnajjar, Miguel Alvelo-Rivera, Michael Charters, Olivia Dionisio, Mohan Kulkarni, Sarah Meehan, Lillian (Anna) Nefcy, Kasia Nowak, Ikenna Okereke, Allison Pekar, Angela Broski (Polanco), Andrew Popoff, Lara Zador, Kortney Barrett, Emine Bayman, Giovanni Berardi, Chris Coffey, Michele Costigan, Dana L Dailey, Dixie J Ecklund, Laura Frey Law, Candy Hodges, Trevis Huff, Adam Janowski, Zackary Lemka, Vincent A Magnotta, Leigh Nadel, Tina Neill-Hudson, Lynn A Rasmussen, Kathleen Sluka, Maggie Spencer, Carol G T Vance, Ezgi Yarasir, Emma Griebenow, Samantha Lariosa, Nondas Leloudas, Hristo Piponov, Pottumarthi Prasad, Maryann Regner, Shanah Salter, Pablo Vicente, Vivien Wang, Anna Lokshin, Panshak Dakup, Jon Jacobs, Wei-Jun Qian, John Burns, Asokumar Buvanendran, Josh Jacobs, Angel Ledesma, Michael Liptay, Silvia Marroquin, Robert McCarthy, Zaki Mehkri, Laura Quigley, Jacey Schickel, Christopher Sica, Chelsey Thomas, James Carson, Vennela Gajjala, Karla Gendler, Ari B Kahn, Donald Koehler, Nathaniel Mendoza, Lissa Pearson, Hedda Prochaska, Pat Scherer, Joshua Urrutia, Matthew Vaughn, Kumari Adams, Elizabeth Goetz, Nivya Kolli, Oliver Fiehn, Tobias Kind, Aravind Athivirahham, Aishat Bakare, Tessa Balach, Derrick Brown, Darren Bryan, Laura Cin, Amy Durkin, Xiaodong Guo, Olivia Keaveny, Hue Luu, Maria Lucia Madariaga, Ashley Mayo, Carla Tisdale, Sara Wallace, Miracle Anderson, Anelizze Castro-Martinez, Katie Fisch, Abbas Hakim, Christopher Hermosillo, Kristen Jepsen, Louise Laurent, Cayla Mason, Tyler Ostrander, Chloe Reeves, Enamul Bhuiyan, Michael Flannery, Hagai Ganin, Rachel Gorre, Muge Karaman, Qingfei Luo, Ping-Shou Zhong, Joe Zhou, Ambar Akhlas, Madhumitha Balaji, Rachel Bergmans, Emby Black, Chad Brummett, Andrew Chang, Joe Chue, Daniel Clauw, Courtney Cole, Douglas Colquhoun, Elizabeth Dailey, Mary Donahue, Kendall Dubois, Maximillian Egan, Sarah Fazal, Stephan Frangakis, Lars Fritsch, DeAnna Hanewald, Steven Harte, Emre Berk Hayir, Esmeralda Hidalgo-Lopez, Eric Ichesco, Jimmy Jagan, Chelsea Kaplan, Sachin Kheterpal, Tony Larkin, Remy Lobo, Maggie Makar, Scott Peltier, Samantha Pianga (Law), Jennifer Pierce, Thomas Prince, Andrew Schrepf, Kathy Scott, Anik Sinha, Michael Sipe, Alexis Stanczuk, Alicia Suydam, Andrew Urquhart, Jennifer Waljee, Noah Waller, Sydney Whack, David Williams, Melanie Wong, Guohao (Arthur) Zhu, Laura Cox, Timothy Howard, Carl Langefeld, Michael Olivier, Sobha Puppala, Ellen Quillen, Arisbeth Reyes, Benlian Wang, Kip Zimmerman, Oluwaseyi “Seyi” Akintoroye, Ramtilak Gattu, Sophie George, Tanja Jovanovic, Shaurel Valbrun, Sterling Winters, Yang Xuan.

## References

[1] Barnett K, Mercer SW, Norbury M, Watt G, Wyke S, Guthrie B. Epidemiology of multimorbidity and implications for health care, research, and medical education: a cross-sectional study. The Lancet 2012;380:37–43.

[2] Barreveld AM, Correll DJ, Liu X, Max B, McGowan JA, Shovel L, Wasan AD, Nedeljkovic SS. Ketamine Decreases Postoperative Pain Scores in Patients Taking Opioids for Chronic Pain: Results of a Prospective, Randomized, Double-Blind Study. Pain Med 2013;14:925–934.

[3] Berardi G, Frey-Law L, Sluka KA, Bayman EO, Coffey CS, Ecklund D, Vance CGT, Dailey DL, Burns J, Buvanendran A, McCarthy RJ, Jacobs J, Zhou XJ, Wixson R, Balach T, Brummett CM, Clauw D, Colquhoun D, Harte SE, Harris RE, Williams DA, Chang AC, Waljee J, Fisch KM, Jepsen K, Laurent LC, Olivier M, Langefeld CD, Howard TD, Fiehn O, Jacobs JM, Dakup P, Qian W, Swensen AC, Lokshin A, Lindquist M, Caffo BS, Crainiceanu C, Zeger S, Kahn A, Wager T, Taub M, Ford J, Sutherland SP, Wandner LD. Multi-Site Observational Study to Assess Biomarkers for Susceptibility or Resilience to Chronic Pain: The Acute to Chronic Pain Signatures (A2CPS) Study Protocol. Frontiers in Medicine 2022;Volume 9 - 2022.

[4] Breivik H, Collett B, Ventafridda V, Cohen R, Gallacher D. Survey of chronic pain in Europe: Prevalence, impact on daily life, and treatment. European Journal of Pain 2006;10:287.

[5] Brooks M, Kristensen K, van Benthem K, Magnusson A, Berg C, Nielsen A, Skaug H, Mächler M, Bolker B. glmmTMB balances speed and flexibility among packages for zero-inflated generalized linear mixed modeling. The R Journal 2017;9:378–400.

[6] Bürkner P. brms: An R Package for Bayesian Multilevel Models Using Stan. J Stat Soft 2017;80:1.

[7] Carpenter B, Gelman A, Hoffman MD, Lee D, Goodrich B, Betancourt M, Brubaker M, Guo J, Li P, Riddell A. Stan: A probabilistic programming language. Journal of statistical software 2017;76:1–32.

[8] Chapman CR, Davis J, Donaldson GW, Naylor J, Winchester D. Postoperative Pain Trajectories in Chronic Pain Patients Undergoing Surgery: The Effects of Chronic Opioid Pharmacotherapy on Acute Pain. The Journal of Pain 2011;12:1240–1246.

[9] Chou LB, Wagner D, Witten DM, Gabriel J. Martinez-Diaz, Brook NS, Toussaint M, Carroll IR. Postoperative Pain Following Foot and Ankle Surgery: A Prospective Study. Foot Ankle Int 2008;29:1063–1068.

[10] Cormier S, Lavigne GL, Choinière M, Rainville P. Expectations predict chronic pain treatment outcomes. Pain 2016;157.

[11] Cox TC, Huntington CR, Blair LJ, Prasad T, Lincourt AE, Heniford BT, Augenstein VA. Predictive modeling for chronic pain after ventral hernia repair. The American Journal of Surgery 2016;212:501–510.

[12] Craig BM, Pickard AS, Lubetkin EI. Health problems are more common, but less severe when measured using newer EQ-5D versions. J Clin Epidemiol 2014;67:93–99.

[13] Dominick CH, Blyth FM, Nicholas MK. Unpacking the burden: Understanding the relationships between chronic pain and comorbidity in the general population. Pain 2012;153:293–304.

[14] Dorris H, Oh J, Jacobson N. Wearable Movement Data as a Potential Digital Biomarker for Chronic Pain: An Investigation Using Deep Learning. Physical Activity and Health 2024;8:83–92.

[15] Ennis DM, Bi J. The Beta-Binomial Model: Accounting for Inter-Trial Variation in Replicated Difference and Preference Tests. J Sens Stud 1998;13:389–412.

[16] Gange SJ, Muñoz A, Sáez M, Alonso J. Use of the Beta-Binomial Distribution to Model the Effect of Policy Changes on Appropriateness of Hospital Stays. Journal of the Royal Statistical Society.Series C (Applied Statistics) 1996;45:371–382.

[17] Garcia-Palacios A, Herrero R, Belmonte MA, Castilla D, Guixeres J, Molinari G, Baños RM, Botella C. Ecological momentary assessment for chronic pain in fibromyalgia using a smartphone: A randomized crossover study. EJP 2014;18:862–872.

[18] Goulet JL, Buta E, Bathulapalli H, Gueorguieva R, Brandt CA. Statistical Models for the Analysis of Zero-Inflated Pain Intensity Numeric Rating Scale Data. The Journal of Pain 2017;18:340–348.

[19] Harris RE, Williams DA, McLean SA, Sen A, Hufford M, Gendreau RM, Gracely RH, Clauw DJ. Characterization and consequences of pain variability in individuals with fibromyalgia. Arthritis & Rheumatism 2005;52:3670–3674.

[20] Hogans BB, Siaton BC, Sorkin JD. Pain when it “counts”: hurdle analysis of clinical pain ratings improves data model performance. PAIN Reports 2025;10.

[21] Jeffrey MA, Wang W, Nelson S. Estimating overall exposure effects for zero-inflated regression models with application to dental caries. Stat Methods Med Res 2014;23:257–278.

[22] Kannampallil T, Galanter WL, Falck S, Gaunt MJ, Gibbons RD, McNutt R, Odwazny R, Schiff G, Vaida AJ, Wilkie DJ, Lambert BL. Characterizing the pain score trajectories of hospitalized adult medical and surgical patients: a retrospective cohort study. Pain 2016;157.

[23] Knowles JE, Frederick C. merTools: Tools for Analyzing Mixed Effect Regression Models., 2025.

[24] Lambert D. Zero-Inflated Poisson Regression, with an Application to Defects in Manufacturing. Technometrics 1992;34:1–14.

[25] Leroux A, Crainiceanu C, Zeger S, Taub M, Ansari B, Wager TD, Bayman E, Coffey C, Langefeld C, McCarthy R, Tsodikov A, Brummet C, Clauw DJ, Edwards RR, Lindquist MA AC. Statistical modeling of acute and chronic pain patient-reported outcomes obtained from ecological momentary assessment. Pain 2024;165.

[26] Leroux A, Xiao L, Crainiceanu C, Checkley W. Dynamic prediction in functional concurrent regression with an application to child growth. Statistics in Medicine 2018;37:1376–1388.

[27] Liu F, Kong Y. zoib: An R Package for Bayesian Inference for Beta Regression and Zero/One Inflated Beta Regression. The R Journal 2015;7:34–51.

[28] Macfarlane GJ, Barnish MS, Jones GT. Persons with chronic widespread pain experience excess mortality: longitudinal results from UK Biobank and meta-analysis. Ann Rheum Dis 2017;76:1815–1822.

[29] Mullahy J. Specification and testing of some modified count data models. J Econ 1986;33:341–365.

[30] Niv D, Kreitler S. Pain and Quality of Life. Pain Practice 2001;1:150–161.

[31] Parker RMA, Tilling K, Terrera GM, Barrett JK. Modeling Risk Factors for Intraindividual Variability: A Mixed-Effects Beta-Binomial Model Applied to Cognitive Function in Older People in the English Longitudinal Study of Ageing. Am J Epidemiol 2024;193:159–169.

[32] Reynolds LW, Hoo RK, Brill RJ, North J, Recker DP, Verburg KM. The COX-2 Specific Inhibitor, Valdecoxib, Is An Effective, Opioid-Sparing Analgesic in Patients Undergoing Total Knee Arthroplasty. J Pain Symptom Manage 2003;25:133–141.

[33] Rikard SM, Strahan AE, Schmit KM, Guy GP. Chronic Pain Among Adults - United States, 2019-2021. MMWR Morb Mortal Wkly Rep 2023;72:379–385.

[34] Rizopoulos D. Dynamic Predictions and Prospective Accuracy in Joint Models for Longitudinal and Time-to-Event Data. Biometrics 2011;67:819–829.

[35] Sadil P, Arfanakis K, Bhuiyan EH, Caffo B, Calhoun VD, Clauw DJ, DeLano MC, Ford JC, Gattu R, Guo X, Harris RE, Ichesco E, Johnson MA, Jung H, Kahn AB, Kaplan CM, Leloudas N, Lindquist MA, Luo Q, Mulderink TA, Peltier SJ, Prasad PV, Sica C, Urrutia J, Vance CG, Wager TD, Xuan Y, Zhou XJ, Zhou Y, Shu DC, The Acute to Chronic Pain Signatures Consortium. Image Processing in the Acute to Chronic Pain Signatures (A2CPS) Project. bioRxiv 2024:2024.12.19.627509.

[36] Sluka KA, Wager TD, Sutherland SP, Labosky PA, Balach T, Bayman EO, Berardi G, Brummett CM, Burns J, Buvanendran A, Caffo B, Calhoun VD, Clauw D, Chang A, Coffey CS, Dailey DL, Ecklund D, Fiehn O, Fisch KM, Frey Law LA, Harris RE, Harte SE, Howard TD, Jacobs J, Jacobs JM, Jepsen K, Johnston N, Langefeld CD, Laurent LC, Lenzi R, Lindquist MA, Lokshin A, Kahn A, McCarthy RJ, Olivier M, Porter L, Qian W, Sankar CA, Satterlee J, Swensen AC, Vance CGT, Waljee J, Wandner LD, Williams DA, Wixson RL, Zhou XJ, the AC. Predicting chronic postsurgical pain: current evidence and a novel program to develop predictive biomarker signatures. Pain 2023;164.

[37] Smith D, Wilkie R, Uthman O, Jordan JL, McBeth J. Chronic Pain and Mortality: A Systematic Review. PLOS ONE 2014;9:e99048.

[38] Traeger AC, Henschke N, Hübscher M, Williams CM, Kamper SJ, Maher CG, Moseley GL, McAuley JH. Estimating the Risk of Chronic Pain: Development and Validation of a Prognostic Model (PICKUP) for Patients with Acute Low Back Pain. PLOS Medicine 2016;13:e1002019.

[39] Vehtari A, Gelman A, Gabry J. Practical Bayesian model evaluation using leave-one-out cross-validation and WAIC. Statistics and Computing 2017;27:1413–1432.

[40] Vittori A, Cascella M, Di Gennaro P, Marchetti G, Francia E, Mascilini I, Tarquini R, Innamorato MA, Petrucci E, Marinangeli F, Coluccia S, Picardo SG. Advanced statistical approaches for predicting pain after pediatric thoracotomy: a cross-sectional study using zero-inflated and Poisson models. Journal of Anesthesia, Analgesia and Critical Care 2024;4:53.

[41] Warton DI. Many zeros does not mean zero inflation: comparing the goodness-of-fit of parametric models to multivariate abundance data. Environmetrics 2005;16:275–289.

