## Supplementary Materials for "Modeling Patient-Reported Pain Trajectories with Frequent Minimum and Maximum Scores"

**Supplementary Tables** **27**

**Supplementary Figures** **34**

### Appendix

#### Appendix 1: An alternative parametrization of the beta-binomial (BB) distribution

Assume that each pain score $y_{ijk}$ follows a BB distribution:

$$y_{ijk} \sim \text{Beta-Binomial}\left(M, \pi_{ijk}\phi, (1 - \pi_{ijk})\phi\right)$$

where $M$ is the maximum possible pain score, $\pi_{ijk}$ is a location parameter, and $\phi > 0$ is the dispersion parameter.

The intuition behind these parameters can be better understood through an alternative parametrization of the distribution. Conditional on a latent probability $p_{ijk}$, the pain score follows a binomial distribution, $y_{ijk}|p_{ijk} \sim \text{Bin}(M, p_{ijk})$, and the latent probability is modelled as $p_{ijk} \sim \text{Beta}(\alpha, \beta)$, with $\alpha = \pi_{ijk}\phi, \beta = (1 - \pi_{ijk})\phi$. The marginal mean and variance of $y_{ijk}$ is $\mathbb{E}[y_{ijk}] = M\pi_{ijk}$ and $Var[y_{ijk}] = M\pi_{ijk}(1 - \pi_{ijk})(1 + (M - 1)\frac{1}{1+\phi})$. The marginal variance can be interpreted as the binomial variance inflated by a multiplicative factor $1 + (M - 1)\frac{1}{1+\phi}$. As $\phi$ decreases, this factor increases, producing greater dispersion. As $\phi$ approaches $\infty$, the factor approaches 1, recovering a standard binomial distribution.

Figure 3 illustrates that the BB distribution can generate zero-inflated, one-inflated, and non-inflated outcomes depending on the choice of $\pi$ and $\phi$. The corresponding beta distributions on the latent probability $p$ are shown in Figure A1. For example, when $\pi = 0.1, \phi = 0.5$, the latent probability follows a beta distribution with parameters $\alpha = 0.05$ and $\beta = 0.45$, which is highly dispersed with mass near zero, resulting in many zero outcomes in the BB distribution. Increasing either $\phi$ or $\pi$ reduces the proportion of zeros. For instance, when $\pi = 0.1, \phi = 5$, $p \sim \text{Beta}(0.5, 0.45)$ which is less dispersed and skewed, resulting in fewer zeros. When $\pi = 0.5, \phi = 5$, the BB distribution closely resembles a symmetric binomial distribution, corresponding to a non-inflated distribution.

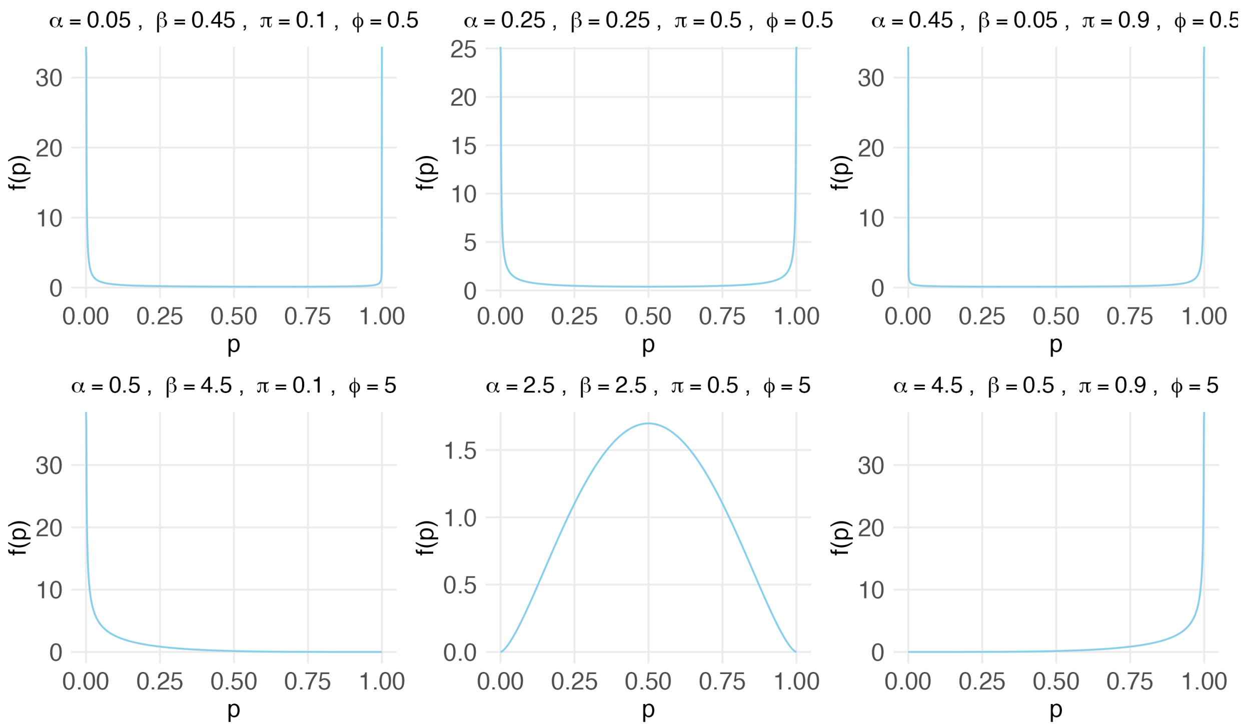

Figure A1: Probability density functions of the Beta distribution under varying $\alpha$ and $\beta$ parameters. The beta-binomial distribution can be understood as a two-step process where the random variable follows a binomial distribution with latent probability $p$, and $p$ itself follows a Beta distribution, thereby accommodating more data variability. We re-parametrize the Beta distribution with a location parameter $\pi$ and a dispersion parameter $\phi$. Together, $\pi$ and $\phi$ controls the amount of zero-inflation and one-inflation.

#### Appendix 2: Specifications of Model Variants

Full model with subject-level dispersion:

$$y_{ijk} \sim BB(M, \pi_{ijk}, \phi_i)$$

$$\eta_{ijk} = \log \frac{\pi_{ijk}}{1-\pi_{ijk}} = r_{ij} + k w_{ij} \qquad \begin{bmatrix} \mathbf{r}_i \\ \mathbf{w}_i \end{bmatrix} \sim \mathrm{MVN}_{2J}\left( \begin{bmatrix} \mathbf{r} \\ \mathbf{w} \end{bmatrix}, \Sigma \right)$$

$$\xi_i = \log \phi_i \qquad \xi_i \sim N(\xi, \sigma_\xi)$$

Full model with timepoint-specific dispersion:

$$y_{ijk} \sim BB(M, \pi_{ijk}, \phi_j)$$

$$\eta_{ijk} = \log \frac{\pi_{ijk}}{1-\pi_{ijk}} = r_{ij} + k w_{ij} \qquad \begin{bmatrix} \mathbf{r}_i \\ \mathbf{w}_i \end{bmatrix} \sim \mathrm{MVN}_{2J}\left( \begin{bmatrix} \mathbf{r} \\ \mathbf{w} \end{bmatrix}, \Sigma \right)$$

Covariate model with subject-level dispersion:

$$y_{ijk} \sim BB(M, \pi_{ijk}, \phi_i)$$

$$\eta_{ijk} = \log \frac{\pi_{ijk}}{1-\pi_{ijk}} = \mathbf{X}\boldsymbol{\beta} + r_{ij} + k w_{ij} \qquad \begin{bmatrix} \mathbf{r}_i \\ \mathbf{w}_i \end{bmatrix} \sim \mathrm{MVN}_{2J}\left( \begin{bmatrix} \mathbf{r} \\ \mathbf{w} \end{bmatrix}, \Sigma \right)$$

$$\xi_i + \mathbf{Z}\boldsymbol{\alpha} = \log \phi_i \qquad \xi_i \sim N(\xi, \sigma_\xi)$$

No slope model with common dispersion:

$$y_{ijk} \sim BB(M, \pi_{ijk}, \phi)$$

$$\eta_{ijk} = \log \frac{\pi_{ijk}}{1-\pi_{ijk}} = r_{ij} \qquad \mathbf{r}_i \sim \mathrm{MVN}_J\left(\mathbf{r}, \Sigma_{\mathbf{r}}\right)$$

Mixed intercept-only model with common dispersion:

$$y_{ijk} \sim BB(M, \pi_{ijk}, \phi)$$
$$\eta_{ijk} = \log \frac{\pi_{ijk}}{1 - \pi_{ijk}} = r_i \qquad\qquad r_i \sim N(r, \sigma_r)$$

Fixed intercept-only model with common dispersion

$$y_{ijk} \sim BB(M, \pi_{ijk}, \phi)$$
$$\eta_{ijk} = \log \frac{\pi_{ijk}}{1 - \pi_{ijk}} = r$$

#### Appendix 3: Inference in Bayesian Framework and Frequentist Approach

Inference was performed using the full model with timepoint-specific dispersion parameter. Results for the full model with subject-level dispersion parameter can be found in Supplementary Table S1, Figure S8-S12.

##### Stan

The primary focus of this paper is on estimating the model parameters in a Bayesian framework. Posterior sampling is done using Hamiltonian Monte Carlo implemented in Stan [3]. Hamiltonian Monte Carlo is a sampling algorithm that is able to efficiently explore complex probability distributions. The parameters of interest include the mean and covariance matrix of the random effects in $\eta_{ijk}$, the linear predictors of location parameters, and the dispersion parameter $\phi$. To improve estimation, we first decompose the covariance matrix of the random effects as $\Sigma = S\Omega S$, where $S$ is the diagonal matrix of standard deviations of the random effects, and $\Omega$ is a correlation matrix with $\Omega_{pp} = 1$ and $\Omega_{pq} = \rho_{pq}$ $(p \neq q)$. We then assign priors

$$\begin{aligned}
\begin{bmatrix} \mathbf{r} \\ \mathbf{w} \end{bmatrix} &\sim \text{MVN}_{2J}\left( \begin{bmatrix} 0 \\ 0 \\ \vdots \\ 0 \end{bmatrix}, \begin{bmatrix} s^2 & 0 & \cdots & 0 \\ 0 & s^2 & \cdots & 0 \\ \vdots & \vdots & \ddots & \vdots \\ 0 & 0 & \cdots & s^2 \end{bmatrix} \right) \\
\text{Diag}(S) &\sim \text{Half-Cauchy}(0, c) \\
\Omega &\sim \text{LKJ}(1) \\
\phi_j &\sim \text{Half-Cauchy}(0, c_\phi)
\end{aligned}$$

Standard deviations and correlation of random effects are both of interest, which leads us to choose the combination of the Half-Cauchy prior for standard deviations and the Lewandowski-Kurowicka-Joe (LKJ) prior for correlation between random effects. This prior setup separates uncertainty about standard deviations and correlations, avoiding the limitations of using a single inverse-Wishart prior. Smaller values of $s$ and $c$ impose stronger prior information, while larger values yield weakly informative priors. The LKJ(1) prior corresponds to a uniform distribution over all valid correlation matrices. The dispersion parameter follows a half cauchy prior. Alternatively, we can use a normal prior on $\log(\phi)$.

Each dataset was fit using four parallel chains, each run with a burn-in of 2000 iterations followed by

4000 sample iterations. Here the burn-in works to warm-up or stabilize the model, and the sample iterations are recorded as actual results. To reduce autocorrelation between consecutive samples and to ensure a sufficiently large effective sample size (ESS) of each parameter, every second sample was saved (thinning factor of 2). ESS reflects the size of independent posterior samples of each parameter; intuitively it is the actual posterior sample size divided by an autocorrelation penalty. We can increase ESS by running longer chains or increasing the thining factor. When all $N$ samples are independent, ESS equals $N$; conversely when the $N$ samples become more autocorrelated, ESS becomes smaller. All models were fit using the default control parameters in Stan. Convergence was monitored for each parameter using traceplots and Gelman-Rubin diagnostics $\hat{R}$, with a threshold of $\hat{R} < 1.05$ indicating convergence [4]. Details on priors for covariates and subject-level dispersion parameters are provided in Appendix 4.

##### brms

Alternatively, the model can also be fit using the brms package in R [2], which provides a user-friendly interface by expressing random effects in the familiar lme4 syntax:

Y | trials(M) ~ 0 + Time + Test_id:Time + (0 + Time + Test_id:Time | ID),

phi ~ 1

The underlying estimation is performed in Stan, but brms simplifies model specification and is easier for users with mixed-effects modeling experience. While brms is convenient for applied users, direct coding in Stan remains more flexible for extending the model structure. In Appendix 8, we compared the performance and runtime of brms and Stan.

##### Frequentist Approach

For comparison, the beta-binomial mixed-effects model (BBME) can also be fit using a frequentist approach, which relies on maximizing the marginal likelihood. This involves iteratively optimizing the likelihood function to obtain maximum likelihood estimates of the parameters. To align with the lme4 syntax used in brms, we used the glmmTMB package in R [1].

##### Appendix 3.1: brms syntax in full model with subject-level dispersion

Y | trials(M) ~ 0 + Time + Test_id:Time + (0 + Time + Test_id:Time | ID),

phi ~ 1 + (1 | ID)

**Appendix 3.2: lmer syntax in full model**

```
Y ~ 0 + Time + Test_id:Time + (0 + Time + Test_id:Time | ID)
```

#### Appendix 4: Prior specifications for model variations

Priors on subject-level dispersion: For $\log(\phi_i) = \xi_i$

$$\xi_i \sim N(\xi, \sigma_\xi)$$
$$\xi \sim N(0, s)$$
$$\sigma_\xi \sim \text{Half-Cauchy}(0, c)$$

Priors when covariates are included:

Let $X$ be the matrix of covariates and $\boldsymbol{\beta}$ be the vector of coefficients of size $P$. We impose a normal prior on $\boldsymbol{\beta}$:

$$\beta_p \sim N(0, \sigma_\beta^2) \quad \text{for } p = 1, \ldots, P$$

When number of covariates is large and some feature selection is needed, we impose a horseshoe prior on $\boldsymbol{\beta}$:

$$\beta_p | \tau, \lambda_p \sim N(0, \lambda_p^2 \tau^2) \quad \text{for } p = 1, \ldots, P$$
$$\lambda_p \sim \text{Half-Cauchy}(0, \lambda_0)$$
$$\tau \sim \text{Half-Cauchy}(0, \tau_0)$$

where $\lambda_p$ is a local shrinkage parameter on each coefficient. $\tau$ is a global shrinkage parameter on all coefficients.

#### Appendix 5: Data generating processes

##### Appendix 5.1: Data generating process in testing model validity

We generated a random sample of data with $I = 500$ subjects, $J = 2$ time points, and $K = 10$ possible assessments per time point. There is 15% chance that each subject was randomly assigned to only one time point, and 85% chance each subject were assigned to both time points. At each time point, subjects completed a random number of pain assessments, with at least four randomly chosen from 10 possible assessments numbered from 0 through 9.

Each pain score $y_{ijk}$ was sampled from a BB distribution with maximum score $M = 10$, location parameter $\pi_{ijk}$. The dispersion parameter varies by timepoint, setting $\phi_{j=1} = 5$ and $\phi_{j=2} = 10$. The location parameter $\pi_{ijk}$ was linked to subject-level random effects through a logit link, where the vector of random effects, $\begin{bmatrix} r_{i1} & r_{i2} & w_{i1} & w_{i2} \end{bmatrix}^T$ was sampled independently from a multivariate normal distribution with mean $(-3.2, 0.6, 0, 0)$, and a $4 \times 4$ covariance matrix $\Sigma = S\Omega S$ where $S = \text{Diag}\begin{bmatrix} 1.5 & 1 & 0.6 & 0.5 \end{bmatrix}$ and $\Omega = \begin{bmatrix} 1 & 0.5 & 0.3 & 0 \\ 0.5 & 1 & -0.3 & 0 \\ 0.3 & -0.3 & 1 & 0 \\ 0 & 0 & 0 & 1 \end{bmatrix}$. This covariance induces a mild positive correlation (0.5) between $r_{i1}$ and $r_{i2}$, a weaker positive correlation (0.3) between $r_{i1}$ and $w_{i1}$, and a weak negative correlation (-0.3) between $r_{i2}$ and $w_{i1}$.

##### Appendix 5.2: Data generating process in model comparison, and null-noise model analysis

We generated a random sample of data with $I = 500$ subjects, $J = 2$ time points, and $K = 10$ possible assessments per time point. There is 15% chance that each subject was randomly assigned to only one time point, and 85% chance each subject were assigned to both time points. At each time point, subjects completed a random number of pain assessments, with at least four randomly chosen from 10 possible assessments numbered from 0 through 9.

Each pain score $y_{ijk}$ was sampled from a BB distribution with maximum score $M = 10$, location parameter $\pi_{ijk}$, and dispersion factor $\phi = 1$. The location parameter $\pi_{ijk}$ was linked to subject-level random effects through a logit link, where the vector of random effects, $\begin{bmatrix} r_{i1} & r_{i2} & w_{i1} & w_{i2} \end{bmatrix}^T$ was sampled independently from a multivariate normal distribution with mean $(-1, 1, 0, 0)$, and a $4 \times 4$ covariance matrix

$$\Sigma = S\Omega S \text{ where } S = \text{Diag}\begin{bmatrix}1 & 2 & 0.6 & 0.5\end{bmatrix} \text{ and } \Omega = \begin{bmatrix}1 & 0.5 & 0.3 & 0\\ 0.5 & 1 & -0.3 & 0\\ 0.3 & -0.3 & 1 & 0\\ 0 & 0 & 0 & 1\end{bmatrix}.$$

**Appendix 5.3: Data generating process in stability test**

Appendix 7 illustrated stability test for simulated sample of varying number of subjects ($I$), $J = 2$ time points, and varying number of assessments ($K$) using 500 different seeds. There is 15% chance that each subject was randomly assigned to only one time point, and 85% chance to both time points. Each subject would be randomly assigned at least 40% of the assessments per time point. To showcase one representative parameter combination, each pain score $y_{ijk}$ was sampled from a BB distribution with maximum score $M = 10$, location parameter $\pi_{ijk}$, and dispersion factor $\phi = 1$. The location parameter $\pi_{ijk}$ was linked to subject-level random effects through a logit link, where the random effects follow

$$\begin{bmatrix}r_{i1}\\ r_{i2}\\ w_{i1}\\ w_{i2}\end{bmatrix} \sim \text{MVN}\left(\begin{bmatrix}-3\\ 0\\ 0\\ 0\end{bmatrix}, \Sigma\right), \text{ with } \Sigma = S\Omega S,\ S = \text{Diag}\begin{bmatrix}1 & 1 & 0.5 & 0.5\end{bmatrix},\ \Omega = \begin{bmatrix}1 & 0.5 & 0.5 & 0\\ 0.5 & 1 & 0 & 0.5\\ 0.5 & 0 & 1 & 0\\ 0 & 0.5 & 0 & 1\end{bmatrix}, \text{ and}$$

dispersion parameter $\phi = 1$.

#### Appendix 6: Simulation Study and Model Performance

##### Appendix 6.1: Zero-Inflation Simulation

BBME model performance was checked on zero-inflated data. We generated data for $I = 500$ subjects, each observed at $J = 2$ time points with $K = 10$ pain assessments per time point. To mimic real-world conditions where subjects do not always complete all assessments, subjects were randomly assigned to one or both time points, with 15% contributing data at only one time point and 85% contributing at both time points. Within each time point, each subjects contributed data for between four and ten assessments. This structure corresponds to a missing completely at random (MCAR) mechanism. Details on the simulated model parameters are provided in Appendix 5.1. Supplementary Figure S1 shows histograms of datasets randomly sampled under this design, and Figure S2 shows pain trajectories of randomly selected subjects.

The parameter values in this simulation were chosen to mimic the distribution of the Acute to Chronic Pain Signatures (A2CPS) baseline worst pain score shown in Figure S3. In particular, the distribution of pain scores at the initial assessment within each timepoint was selected to resemble the baseline distributions observed in the total knee arthroplasty (TKA) and thoracic cohorts (Figure S3). The dispersion parameter varies by time point. In a representative simulated dataset, pain scores at time point $j = 1, k = 0$ contained approximately 74.2% zeros and 0% tens, whereas pain scores at time point $j = 2, k = 0$ contained approximately 2.5% zeros and 11.9% tens. Exact percentages varied randomly across simulated datasets. Thus, this simulation provides an example in which one time point exhibits substantial zero inflation while the other is relatively non-inflated. The simulated dataset was then fit in Stan using the procedure described in Appendix 3. Model performance was evaluated by comparing true parameter values with posterior means, and by assessing coverage of the resulting 95% credible intervals.

All model parameters converged with $\hat{R} = 1$ (Table 1, Figure S4). Table 1 shows that the posterior means for all model parameters were close to the true value.

Figure A2 shows scatter plots comparing the true and estimated values of the individual pain score specific location parameters ($\pi_{ijk}$), linear predictors ($\eta_{ijk}$), and random intercepts ($r_{ij}$) and slopes ($w_{ij}$). Most subject-level parameter estimates were close to the identity line, indicating little systematic bias. Some boundary effects were observed, with posterior mean estimates showing slight bias at extreme values of $\eta_{ijk}$ and the random effects. However, these differences had minimal practical impact on the estimated $\pi_{ijk}$ due to the behavior of the inverse logit function, as shown by the estimates of $\pi_{ijk}$ scatter around the identity line in Figure A2. In addition, approximately 95% of the true $\pi_{ijk}$ and $\eta_{ijk}$ are contained within their respective 95% credible intervals, both overall and consistently across all score levels from 0 to 10. Specifically, the coverage is 95.5% at score 0 (no pain) and 94.1% at score 10 (maximum pain). The detailed percentages for each score level are provided in table S2. Also, as shown in Supplementary Figure S5 and S6, the true and posterior mean probability mass function curves are closely aligned both for randomly chosen subjects and for those with largest difference between true and posterior linear predictor $\eta_{ijk}$. This suggests that the model accurately recovered the underlying distributions of individual pain scores, even when extreme parameter values on the link scale introduced small biases. From a clinical perspective, this means the model still captured each subject's overall pain pattern well.

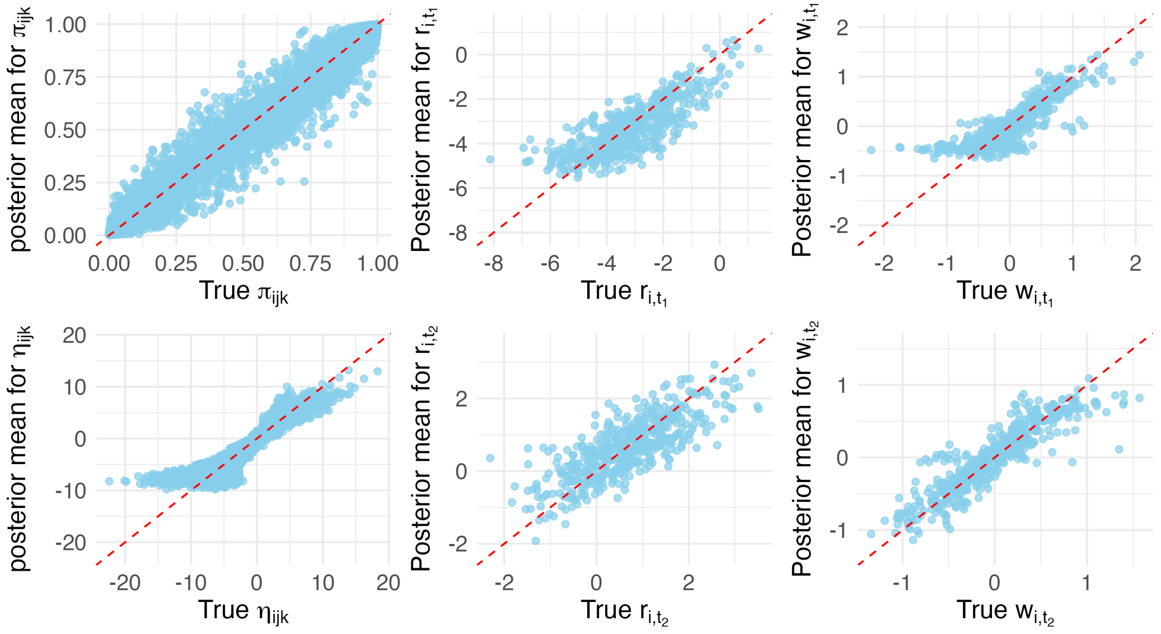

Figure A2: Simulation results of random effects in full BBME with common dispersion for zero-inflation simulation study in Appendix 6.1. Scatter plots show posterior means of subject-specific location parameters, linear predictors, random intercepts and slopes against corresponding true values. The red line indicates the identity line.

The slight bias of estimated $\eta_{ijk}$ arises from the logit transformation $\eta_{ijk} = log(\pi_{ijk}/(1-\pi_{ijk}))$, which goes to $-\infty$ as $\pi_{ijk} \to 0$ and goes to $\infty$ as $\pi_{ijk} \to 1$. Small changes on $\pi_{ijk}$ near the boundaries corresponds to big changes on the logit scale. Consequently, although we observe some bias in the posterior mean of $\eta_{ijk}$, posterior means of $\pi_{ijk}$ remain largely unbiased, and the 95% credible intervals of $\pi_{ijk}$ and $\pi_{ijk}$ were able to contain the true values, suggesting that the model provides reliable estimates even near the boundaries.

**Appendix 6.2: One-Inflation Simulation**

BBME model performance was also checked on one-inflated data. We generated data using the same procedure described in Appendix 5.2, besides changing mean of random effects to $\begin{pmatrix} r_1 & r_2 & w_1 & w_2 \end{pmatrix}^T = \begin{pmatrix} 0 & 3 & 0 & 0 \end{pmatrix}^T$, and common dispersion parameter $\phi = 10$. The distribution of the simulated data is shown in Figure A3. We then fit the simulated dataset in Stan using the procedure described in Appendix 3. Parameter estimates are summarized in Table A1. All parameters converged with $\hat{R} = 1$ (Table A1, Figure S7). Posterior means for all the parameters were very close to the true values, and corresponding 95% credible intervals included the true values as well. Figure A4 shows the posterior subject-level random effects are also close to their true values.

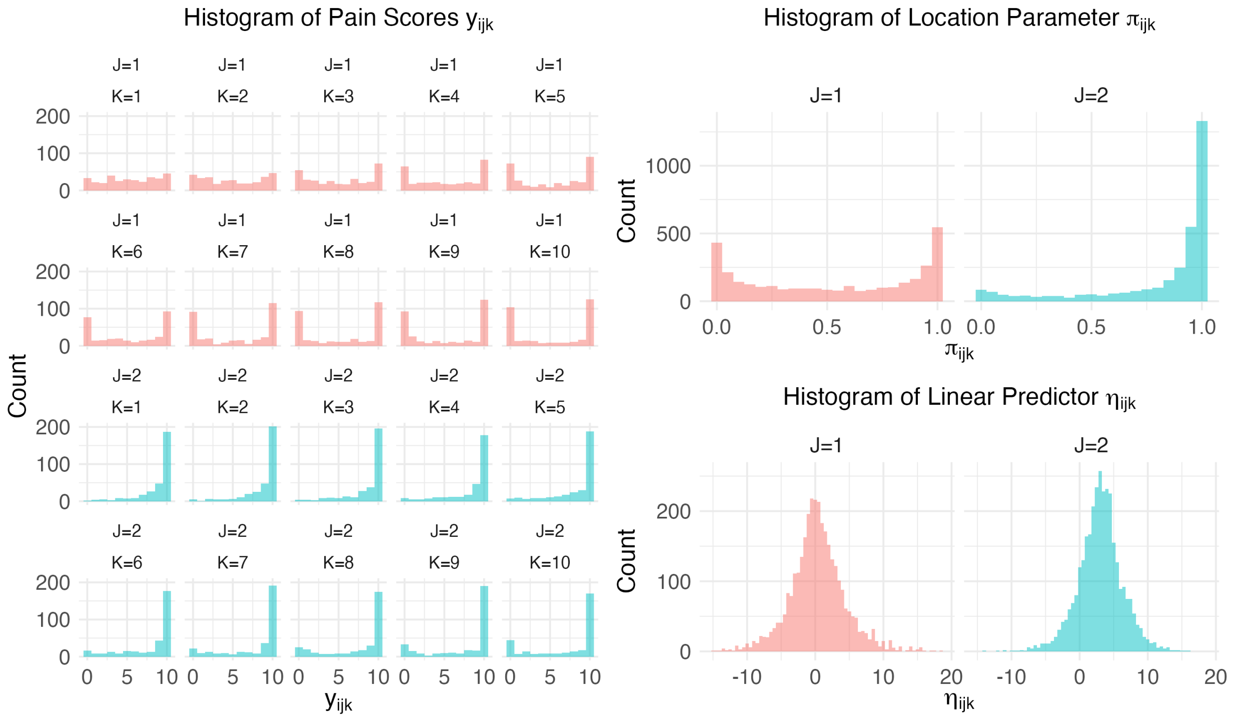

Figure A3: Histograms of simulated one-inflated data. Pain scores, location parameters ($\pi_{ijk}$), and linear predictors ($\eta_{ijk}$) from one simulated dataset of 500 subjects, up to 2 time points, and 4-10 pain scores per time point.

Table A1: True and posterior estimates of simulated data that is one-inflated. Data were generated for 500 subjects, each observed at up to 2 time points with 4-10 pain scores per time points. There is a 15% chance subjects only have one time point. The model was fit using 4 parallel chains with 2000 burn-in iterations followed by 4000 iterations, applying a thinning factor of 2. All model parameters converged with $\hat{R} = 1$

| Parameter | True | Posterior mean | SD | Posterior median | 2.5% quantile | 97.5% quantile | ESS |
|---|---|---|---|---|---|---|---|
| $r_1$ | 0.0 | 0.08 | 0.08 | 0.08 | -0.08 | 0.25 | 6996 |
| $r_2$ | 3.0 | 2.96 | 0.12 | 2.96 | 2.73 | 3.21 | 5915 |
| $w_1$ | 0.0 | 0.04 | 0.03 | 0.04 | -0.02 | 0.10 | 6867 |
| $w_2$ | 0.0 | 0.01 | 0.03 | 0.01 | -0.05 | 0.07 | 5598 |
| $\sigma_{r1}$ | 1.5 | 1.56 | 0.08 | 1.55 | 1.40 | 1.72 | 5027 |
| $\sigma_{r2}$ | 2.0 | 1.89 | 0.11 | 1.88 | 1.67 | 2.12 | 4208 |
| $\sigma_{w1}$ | 0.6 | 0.61 | 0.03 | 0.61 | 0.55 | 0.68 | 5168 |
| $\sigma_{w2}$ | 0.5 | 0.51 | 0.03 | 0.51 | 0.46 | 0.58 | 4242 |
| $\rho_{r1,r2}$ | 0.5 | 0.47 | 0.06 | 0.47 | 0.35 | 0.58 | 4005 |
| $\rho_{r1,w1}$ | 0.3 | 0.39 | 0.06 | 0.39 | 0.27 | 0.49 | 4980 |
| $\rho_{r1,w2}$ | 0.0 | 0.04 | 0.07 | 0.04 | -0.10 | 0.18 | 5056 |
| $\rho_{r2,w1}$ | -0.3 | -0.28 | 0.06 | -0.29 | -0.41 | -0.16 | 5179 |
| $\rho_{r2,w2}$ | 0.0 | -0.06 | 0.07 | -0.07 | -0.20 | 0.07 | 5040 |
| $\rho_{w1,w2}$ | 0.0 | -0.05 | 0.07 | -0.05 | -0.18 | 0.08 | 6338 |
| $\phi$ | 10.0 | 9.69 | 0.57 | 9.67 | 8.62 | 10.87 | 5765 |

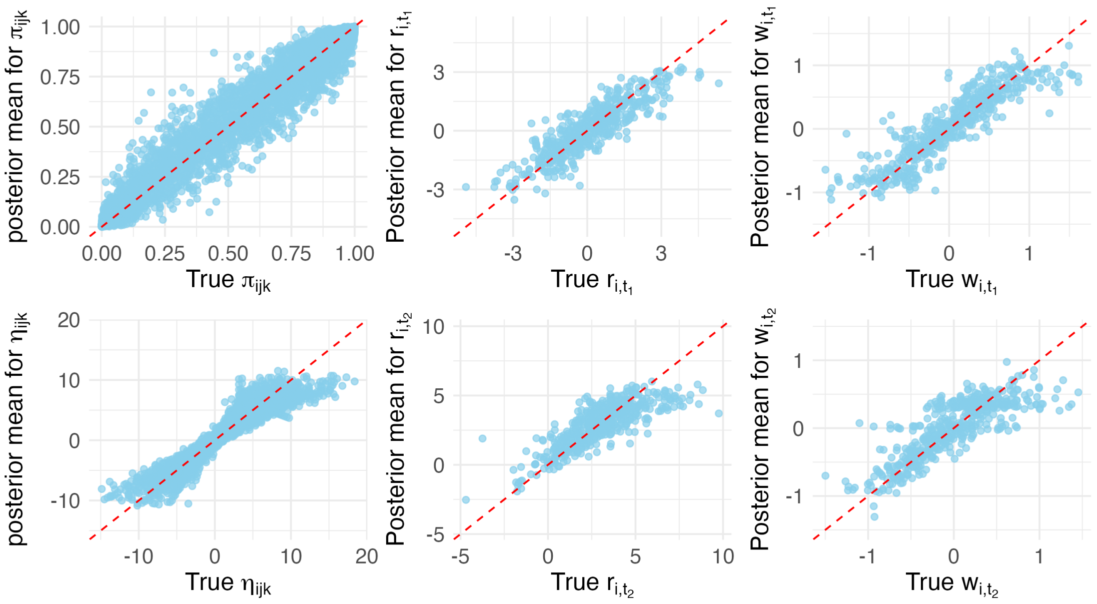

Figure A4: Simulation results of the beta-binomial mixed-effects model (BBME) on one-inflated data. Scatter plots show posterior means of subject-specific location parameters, linear predictors, random intercepts and slopes against corresponding true values. The red line indicates the identity line.

#### Appendix 7: Stability Test

We performed a stability analysis to assess the performance of the BBME model under different parameter settings and sample sizes. The model is designed to flexibly model zero or one-inflated pain scores across multiple time points. To investigate how model performance and stability are influenced by different conditions, we varied the following parameters: (1) mean ($r_t$) of the random intercepts ($r_{i,t}$) were set to $-3$, 0, and 3, corresponding to the location parameters in zero-inflated, non-inflated, and one-inflated distributions, respectively; (2) correlations among random effects ($r_{i,t}$ and $w_{i,t}$) were set to either 0 or 0.5; (3) dispersion was set to 1 or 10; (4) the number of subjects ($I$) were set to 200, 500, and 1000; (5) the number of time points ($J$) were set to be 1, 2, and 3; (6) the number of pain scores ($K$) at each time points were set to 4, 7, and 12. The time points and assessments available for each subject were randomized. We fixed the maximum of the pain scores ($M$) to be 10, standard deviations of the random intercepts ($r_{i,t}$) to be 1, and standard deviations of the random slopes ($w_{i,t}$)) to be 0.5. For each simulation, we randomly generated 500 datasets on each combination of parameters using the random seeds 1 to 500. Each dataset was fit using four parallel chains, each run with a burn-in of 2000 iterations followed by 6000 sample iterations, and the default control parameters in Stan. For each model parameter, we calculated the mean of posterior mean and 95% credible intervals across model fits. Coverage probabilities were then computed to assess how often the 95% intervals contained the true parameter values.

Figure A5 shows coverage probabilities for simulated sample of one representative combination of parameters; details of data generation can be found in Appendix 5.3. Under these parameters, pain scores at time point $j = 1$ are more zero-inflated while scores at time point $j = 2$ are more symmetric: take $k = 7$ as example, a random dataset can contain approximately 75% zeros and 5% tens, while pain scores at time point $j = 2$ contain about 30% zeros and 40% tens. As shown in Figure A5, coverage probabilities of all parameters are close to 95% in all sample sizes. However, the width of the credible intervals decreases as the sample size increases, either by adding more subjects or by increasing the number of pain scores per time point. When the total number of subjects is small ($I = 200$), increasing the number of pain scores per time points from $K = 4$ to $K = 12$ effectively reduces the width of the credible intervals.

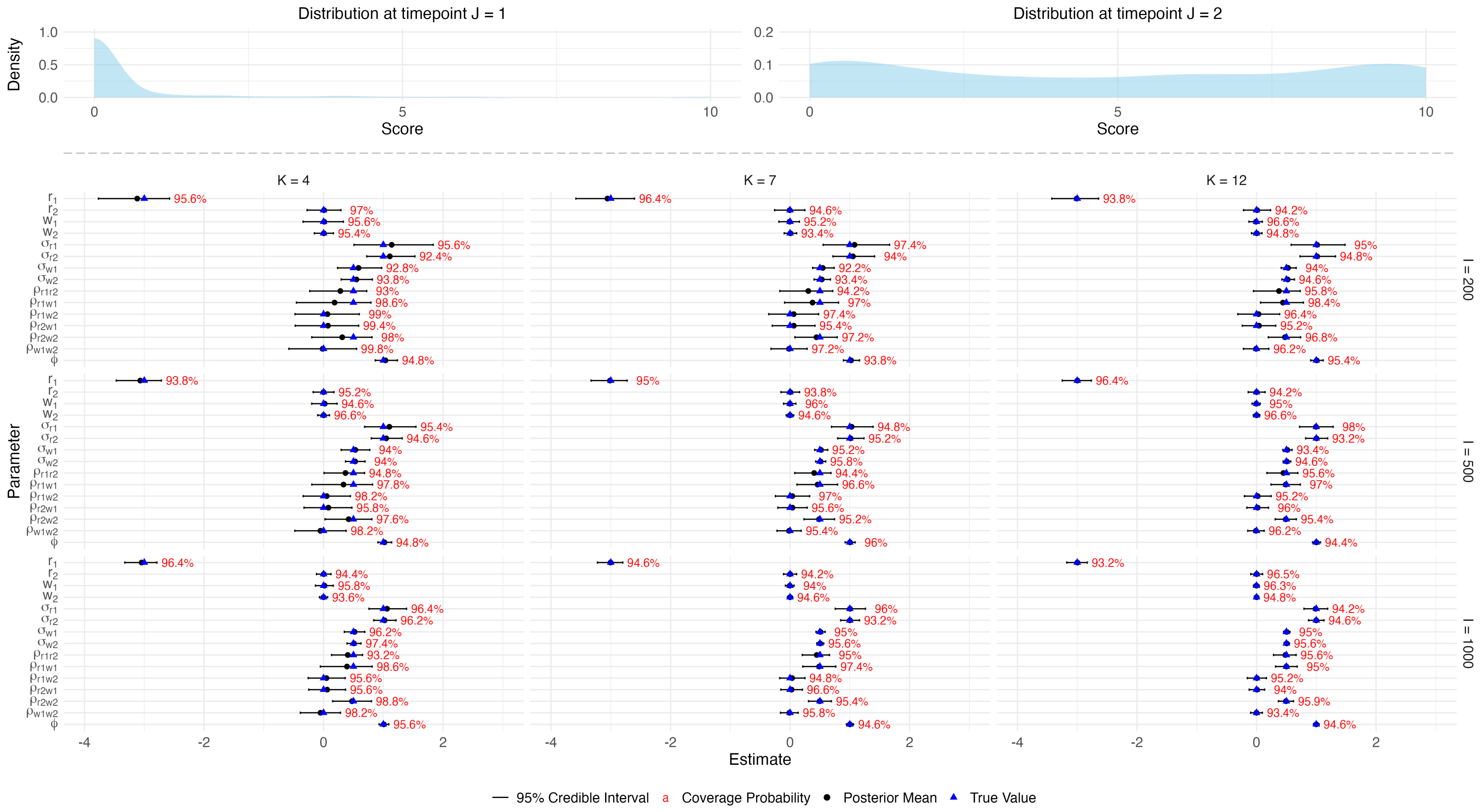

Figure A5: Mean of posterior mean, 95% credible intervals, and coverage probabilities from simulations with varying number of subjects ($I$) and assessments per time point ($K$). All beta-binomial pain scores are sampled under the same parameter settings, details in Appendix 5.1. Black lines indicate 95% credible intervals, blue triangles represent true parameter values, and red numbers denote the coverage probabilities.

#### Appendix 8: Model Comparison between Bayesian and Frequentist Approach

We compared the performance of BBME under Bayesian and Frequentist frameworks. Datasets were generated using 500 random seeds following the setup described in Appendix 5.2. Each dataset was fit using Stan, brms, and glmmTMB in R. For the Stan and brms models, we used the same set of default control parameters. We ran four parallel chains, each with a burn-in of 2000 chains, followed by 4000 chains and a thinning factor of 2. The glmmTMB model was fit using the default optimizer, nlminb, which handles both unconstrained and box-constrained optimization problems. Each model is run on a maximum of 1000 major iterations and a maximum of 1000 evaluations and objective functions.

In addition, we also compared the runtime of model fit across the three packages and their peak RAM usage. Datasets were generated using 500 random seeds following the same procedure described in Appendix 5.2, except that we also varied the number of subjects to be $I = 500$ or $I = 1000$, and $K = 7, 10$, or 14. We still ran four parallel chains of 2000 burn-in and 4000 sampling iterations on brms and stan using the same default control parameters. glmmTMB models were also fit as described above. The four parallel chains of stan and brms were run on separate cores of an Intel Xeon Silver 4309Y CPU (2.80 GHz), and glmmTMB model was run on one core.

##### Appendix 8.1: Performance

We compared the performance of the BBME model under Bayesian and frequentist estimation frameworks using the same dataset simulated in Appendix 5.2. From each model, we recorded parameter estimates along with their uncertainty intervals: for the Bayesian methods we collected the posterior mean and 95% credible intervals, and for the frequentist methods we collected the point estimates and 95% confidence interval. These outputs were used to calculate empirical coverage probabilities for each parameter for each method, defined as the proportion of intervals containing the true value.

Figure A6 and S13 show the posterior means or estimates, 95% credible or confidence intervals, and coverage probabilities of parameters obtained using Stan, brms, and glmmTMB. The performance of brms and Stan was highly similar, with both producing tighter 95% intervals for correlation parameters compared to glmmTMB. Coverage probabilities for Stan and brms were consistently around 95% for all parameters, whereas glmmTMB showed substantially lower coverage for some parameters. In summary, the Bayesian approaches Stan and brms are preferrable in terms of estimation accuracy.

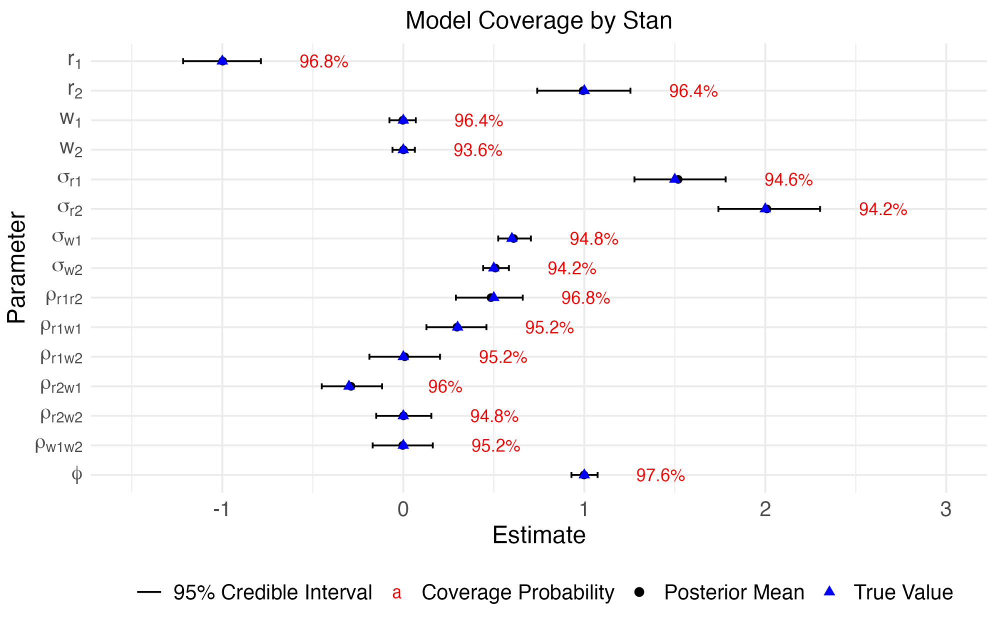

(a)

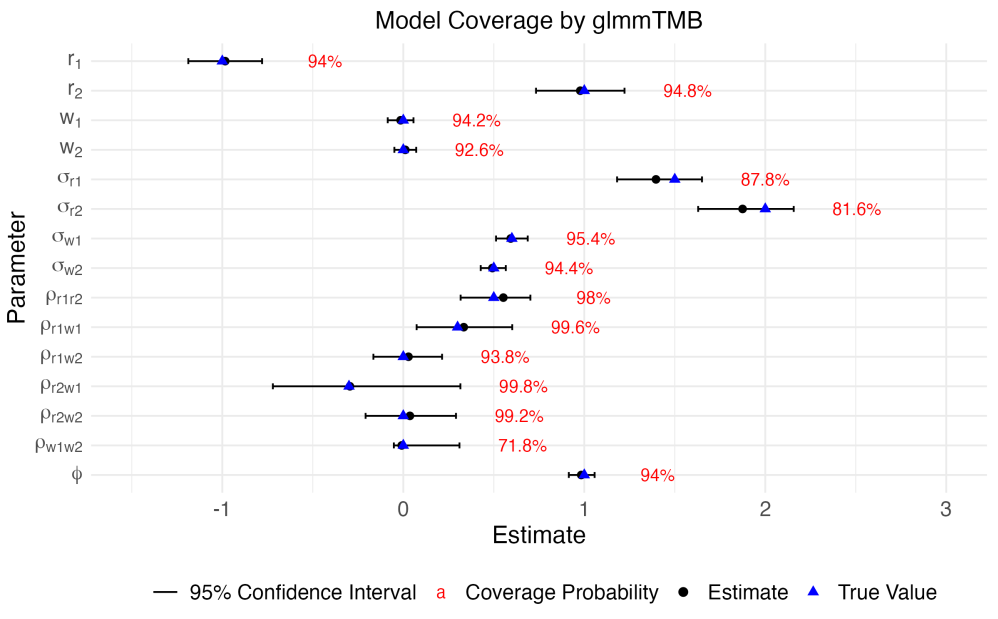

(b)

Figure A6: Dataset were simulated under 500 seeds using the same parameter settings and fitted on (a) Stan [3] and (b) glmmtmb [1]. From each model, 95% credible intervals (Stan) and 95% confidence intervals (glmmTMB) were collected to calculate coverage probabilities across the 500 model fits. We also calculated the mean of posterior mean or estimates, and mean of 95% intervals over the 500 model fits. Black lines indicate intervals, blue triangles mark true parameter values, and red labels denote coverage probabilities.

**Appendix 8.2: Runtime**

Over the 500 replications, glmmTMB had the shortest runtime (mean = 27 seconds, SD = 20 seconds for a dataset of $I = 500, J = 2, K = 14$), but at the cost of reduced coverage. The Bayesian methods required considerably longer runtimes. While accuracy was similar between Stan and brms, Stan scaled better with increasing data size. As shown in Figure A7(a), increasing the number of assessments per time point caused runtime in brms to grow much faster than in Stan. However, this came at the expense of memory usage. Figure A7(b) shows that peak RAM consumption in Stan increased with sample size, whereas brms maintained relatively stable RAM usage.

In summary, Stan is preferred when shorter runtime is desired and sufficient RAM is available. brms is preferable if RAM on the device is limited, despite longer runtime. glmmTMB remains a good choice for obtaining a quick parameter estimates when computational speed is the primary concern and high coverage is not required.

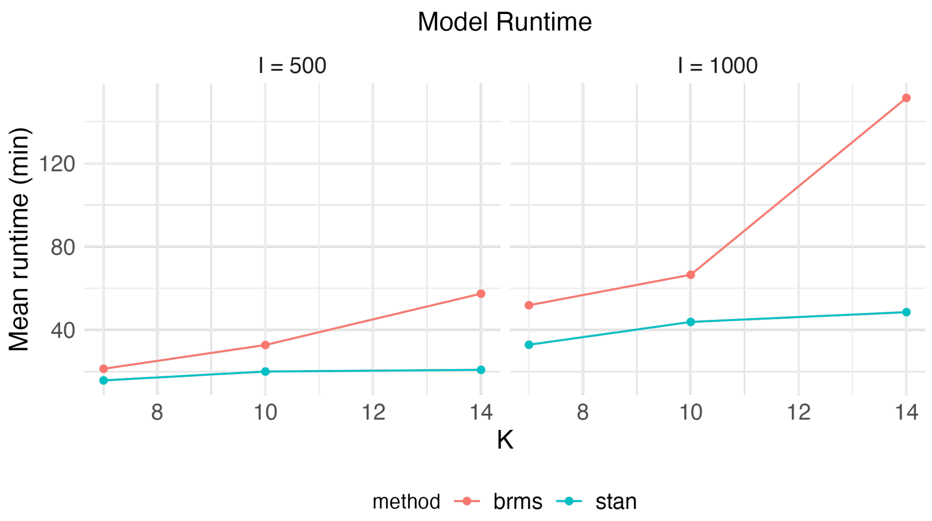

(a)

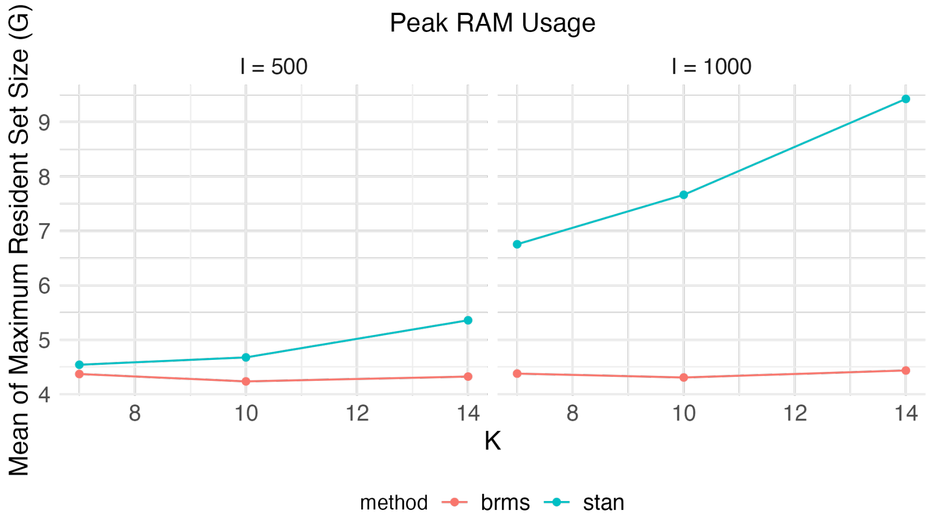

(b)

Figure A7: 500 datasets were simulated with varying numbers of subjects ($I$), $J = 2$ time points, and differing numbers of assessments per time point ($K$), using the same set of parameters but different seeds. Each dataset was fit using Stan and brms. (a) Runtime and (b) peak RAM usage were recorded, and the mean for each sample size and package configuration are shown.

#### Appendix 9: Null Noise Analysis

We performed a model comparison to test whether the BBME model might incorrectly interprets random noise as significant effects. Data was simulated using 500 random seeds following the procedure described in Appendix 5.2. For each of the 500 simulated dataset, we replaced the outcome Y with a random sample drawn from a BB distribution with maximum score $M = 10$, location parameter $\pi = 0.5$, and dispersion parameter $\phi = 1$. This broke the link between the outcome and the explanatory variables. Each of the 500 datasets was fit using four model variants with a common dispersion parameter: a full model; a subject and timepoint specific mixed intercept-only model (no slope model); a mixed intercept-only model; and a fixed intercept-only model (null model), see Appendix 2 for specifications of the latter three models. The fixed intercept only model is the true model according to the data-generating process. We took the mean of posterior mean estimates and 95% credible intervals over the model fits across the 500 datasets. Because the data-generating process assumed no differences across time points or assessments, all 95% credible intervals for fixed effects $(r_1, r_2, w_1, w_2)$ should cover 0 (since $0.5 = \text{expit}(0)$). Failure to do so would indicate that the model is detecting spurious signals. Model fit was assessed using leave-one-out cross validation information criteria (LOOIC) [5], with lower values indicating better out-of-sample fit.

Comparing the different types of models on the 500 simulated datasets, all four models fit the data similarly (Table A2). As shown in Figure A8, the posterior 95% credible intervals for the fixed effects at the two time points covered zero with probabilities close to 95%. These results suggest that the misspecified models did not mistakenly interpret noise in the data as significant effects.

Table A2: Mean LOOIC over 500 model fits, for the full model, no slope model, mixed intercept-only model, and fixed intercept-only model. All four models achieved similar model fits in terms of LOOIC.

| Model | LOOIC |
|---|---|
| Full Model | 29,886 |
| No Slope Model | 29,868 |
| Mixed Intercept-Only Model | 29,587 |
| Fixed Intercept-Only Model (Null Model) | 29,624 |

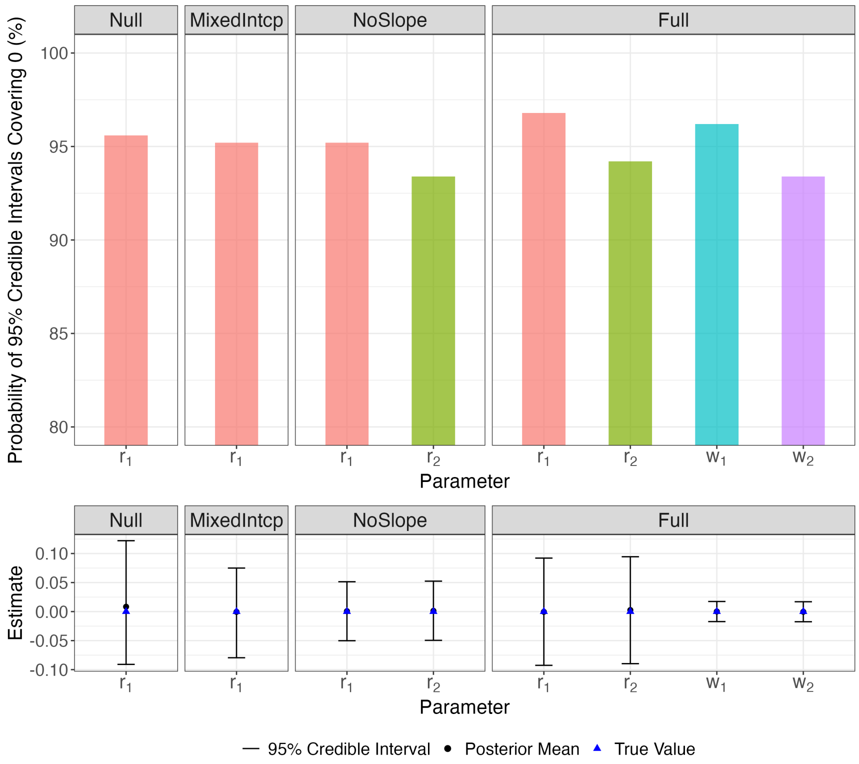

Figure A8: Mean of posterior means and 95% credible intervals of parameters, and probabilities of 95% credible intervals covering 0 (the true value), computed over 500 simulations, for the Null (Fixed Intercept-only), Mixed Intercept-only, No Slope, and Full models. The 95% credible intervals of all fixed effects covered 0 with a probability around 95%, suggesting that the more complex models did not detect random noise as significant effects.

#### Appendix 10: Model Fitting on Real Data

##### Appendix 10.1: FM Study

In FM study, we fit BBME model on the sample of $I = 147$ subjects, $J = 4$ time points, and $K = 45$ pain assessments per time point in Stan [3]. We ran 4 parallel chains of 2000 burn-in iterations, followed by 2000 sampling iterations, saving every second posterior draw for each parameter. Control parameter adapt_delta was set to 0.85, and all other settings remained as default in Stan. All parameters converged with $\hat{R} = 1$.

For comparison to LMM, we fit using syntax

```
PAIN ~ 0 + week + test_id:week + (0 + week + test_id:week | ID)
```

using lmer() function in R package lme4, and

```
PAIN ~ 0 + week + test_id:week + treatment + (0 + week + test_id:week | ID)
```

if adjusting for binary covariate of whether the subject is in treatment group.

##### Appendix 10.2: A2CPS Cuff Pressure Pain Data

In A2CPS study, we fit a sample of $I = 495$ subjects, $J = 2$ time points, and $K = 3$ pain assessments per time point. Note that pain score is first doubled to the range of 0 to 20. We ran 4 parallel chains of 4000 burn-in iterations, followed by 4000 sampling iterations, and saved every second posterior draw for each parameter. Control parameters were set to adapt_delta = 0.85 and max_treedepth = 12; all other settings remained as default values in Stan. All model parameters converged with $\hat{R} = 1$.

For comparison to LMM, we fit using syntax

```
PAIN ~ 0 + period + location:period + cohort + (0 + period + location:period | ID)
```

using lmer() function in R package lme4.

#### Appendix 11: Timepoint-Specific BBME on A2CPS Cuff Pain Dataset

One advantage of BBME is that it can accommodate different dispersion parameters across time points or subjects. To explore this, we fit the same model but allowed seperate dispersion parameters for CUFF1 and REST2. This alternative model achieved a log-likelihood of $-3110.90$ evaluated at the posterior mean parameters, which was very close to that of the BBME with common dispersion parameter. The posterior mean dispersion was 1529.6 for CUFF1 and 391.20 for REST2. Estimates of the other parameters remained close to those from the common-dispersion model. These results suggest that allowing timepoint-specific dispersion did not materially improve fit in this dataset, although it did indicate some differences in variability across the two sessions. In practical terms, the simpler common-dispersion model may therefore be adequate for these data.

Table A3: Posterior estimates for cuff pressure pain in the A2CPS study, including mean of random intercepts ($r$), mean of random slopes (w), standard deviations of random effects ($\sigma$), and timepoint-specific dispersion ($\phi_j$). We ran Four parallel chains of 6000 warm up and 4000 sampling iterations, thinning = 2. All model parameters converged with $\hat{R} < 1.05$.

| Parameter | Posterior mean | SD | Posterior median | 2.5% quantile | 97.5% quantile | ESS |
|---|---|---|---|---|---|---|
| $r_{\text{CUFF1}}$ | -0.87 | 0.08 | -0.86 | -1.06 | -0.72 | 1094 |
| $r_{\text{REST2}}$ | -6.62 | 0.42 | -6.60 | -7.51 | -5.85 | 952 |
| $w_{\text{CUFF1}}$ | 0.34 | 0.03 | 0.34 | 0.28 | 0.39 | 5640 |
| $w_{\text{REST2}}$ | -0.64 | 0.27 | -0.62 | -1.23 | -0.17 | 279 |
| $\sigma_{r_{\text{CUFF1}}}$ | 0.99 | 0.05 | 0.99 | 0.90 | 1.09 | 3866 |
| $\sigma_{r_{\text{REST2}}}$ | 4.03 | 0.36 | 4.01 | 3.40 | 4.79 | 949 |
| $\sigma_{w_{\text{CUFF1}}}$ | 0.45 | 0.03 | 0.45 | 0.39 | 0.50 | 3755 |
| $\sigma_{w_{\text{REST2}}}$ | 0.75 | 0.14 | 0.71 | 0.52 | 1.07 | 308 |
| $\rho_{r_{\text{CUFF1}},r_{\text{REST2}}}$ | 0.27 | 0.07 | 0.27 | 0.13 | 0.41 | 3797 |
| $\rho_{r_{\text{CUFF1}},w_{\text{CUFF1}}}$ | -0.25 | 0.07 | -0.26 | -0.39 | -0.11 | 3575 |
| $\rho_{r_{\text{CUFF1}},w_{\text{REST2}}}$ | -0.21 | 0.13 | -0.21 | -0.47 | 0.03 | 935 |
| $\rho_{r_{\text{REST2}},w_{\text{CUFF1}}}$ | 0.06 | 0.08 | 0.06 | -0.11 | 0.22 | 3132 |
| $\rho_{r_{\text{REST2}},w_{\text{REST2}}}$ | 0.46 | 0.19 | 0.48 | 0.02 | 0.76 | 294 |
| $\rho_{w_{\text{CUFF1}},w_{\text{REST2}}}$ | 0.28 | 0.12 | 0.28 | 0.04 | 0.51 | 1238 |
| $\beta_{\text{TKA}}$ | 0.03 | 0.08 | 0.01 | -0.11 | 0.22 | 782 |
| $\phi_{\text{CUFF1}}$ | 1529.56 | 29870.49 | 311.42 | 99.32 | 5759.99 | 7668 |
| $\phi_{\text{REST2}}$ | 391.20 | 2226.50 | 137.03 | 47.93 | 1702.78 | 5962 |

#### Supplementary Tables

Table S1: True and posterior estimates of simulated data with subject-level dispersion. Data were generated for 500 subjects, each observed at up to 2 time points with 4-10 pain scores per time points. There is a 15% chance subjects only have one time point. The model was fit using 4 parallel chains with 2000 burn-in iterations followed by 4000 iterations, applying a thinning factor of 2. All model parameters converged with $\hat{R} = 1$

| Parameter | True | Posterior mean | SD | Posterior median | 2.5% quantile | 97.5% quantile | ESS |
|---|---|---|---|---|---|---|---|
| $r_1$ | -1.0 | -0.82 | 0.10 | -0.82 | -1.02 | -0.62 | 4463 |
| $r_2$ | 1.0 | 0.96 | 0.12 | 0.96 | 0.72 | 1.20 | 5269 |
| $w_1$ | 0.0 | -0.02 | 0.04 | -0.02 | -0.09 | 0.05 | 5392 |
| $w_2$ | 0.0 | 0.00 | 0.03 | 0.00 | -0.06 | 0.06 | 5392 |
| $\sigma_{r1}$ | 1.5 | 1.58 | 0.12 | 1.58 | 1.35 | 1.83 | 1663 |
| $\sigma_{r2}$ | 2.0 | 2.02 | 0.14 | 2.02 | 1.77 | 2.30 | 2629 |
| $\sigma_{w1}$ | 0.6 | 0.62 | 0.05 | 0.62 | 0.54 | 0.72 | 1906 |
| $\sigma_{w2}$ | 0.5 | 0.52 | 0.03 | 0.52 | 0.46 | 0.59 | 2954 |
| $\rho_{r1,r2}$ | 0.5 | 0.40 | 0.08 | 0.40 | 0.24 | 0.54 | 1914 |
| $\rho_{r1,w1}$ | 0.3 | 0.35 | 0.08 | 0.35 | 0.19 | 0.50 | 1796 |
| $\rho_{r1,w2}$ | 0.0 | -0.05 | 0.08 | -0.05 | -0.21 | 0.10 | 3098 |
| $\rho_{r2,w1}$ | -0.3 | -0.29 | 0.07 | -0.29 | -0.42 | -0.15 | 3293 |
| $\rho_{r2,w2}$ | 0.0 | -0.13 | 0.07 | -0.13 | -0.27 | 0.02 | 3586 |
| $\rho_{w1,w2}$ | 0.0 | 0.13 | 0.07 | 0.13 | -0.01 | 0.27 | 3983 |
| $\mu_{\log(\phi)}$ | 0.0 | 0.01 | 0.07 | 0.01 | -0.13 | 0.14 | 5208 |
| $\sigma_{\log(\phi)}$ | 1.0 | 1.07 | 0.07 | 1.07 | 0.94 | 1.20 | 3932 |

Table S2: The proportion of true random effects $\pi_{ijk}$ and $\eta_{ijk}$ contained within their respective 95% credible intervals (CrI) for each pain score on 0 to 10. Overall, 95.5% of the true values of random effects are contained within the respective 95% credible intervals.

| Pain Scores | Count | Proportions contained in 95% CrI |
|---|---|---|
| 0 | 2471 | 0.955 |
| 1 | 450 | 0.956 |
| 2 | 352 | 0.955 |
| 3 | 287 | 0.962 |
| 4 | 276 | 0.971 |
| 5 | 278 | 0.957 |
| 6 | 268 | 0.940 |
| 7 | 288 | 0.972 |
| 8 | 299 | 0.960 |
| 9 | 364 | 0.967 |
| 10 | 1083 | 0.941 |

Table S3: LMM estimates on the FM study. All correlation parameters are shown in the Figure S14(b).

| Parameter | Estimate | Std Error |
|---|---|---|
| $r_{wk1}$ | 73.89 | 1.54 |
| $r_{wk2}$ | 80.01 | 1.48 |
| $r_{wk3}$ | 63.14 | 2.37 |
| $r_{wk4}$ | 64.86 | 2.34 |
| $w_{wk1}$ | 0.19 | 0.04 |
| $w_{wk2}$ | 0.05 | 0.06 |
| $w_{wk3}$ | 0.07 | 0.04 |
| $w_{wk4}$ | 0.04 | 0.05 |
| $\sigma_{r_{wk1}}$ | 18.24 | |
| $\sigma_{r_{wk2}}$ | 17.38 | |
| $\sigma_{r_{wk3}}$ | 26.85 | |
| $\sigma_{r_{wk4}}$ | 26.41 | |
| $\sigma_{w_{wk1}}$ | 0.44 | |
| $\sigma_{w_{wk2}}$ | 0.49 | |
| $\sigma_{w_{wk3}}$ | 0.42 | |
| $\sigma_{w_{wk4}}$ | 0.42 | |

Table S4: LMM estimates on the A2CPS cuff pressure pain dataset

| Parameter | Estimates | Std Error |
|---|---|---|
| $r_{\text{CUFF1}}$ | 6.3 | 0.28 |
| $r_{\text{REST2}}$ | 0.89 | 0.25 |
| $w_{\text{CUFF1}}$ | 1.39 | 0.11 |
| $w_{\text{REST2}}$ | -0.02 | 0.05 |
| $\sigma_{r_{\text{CUFF1}}}$ | 3.78 | |
| $\sigma_{r_{\text{REST2}}}$ | 2.66 | |
| $\sigma_{w_{\text{CUFF1}}}$ | 2.13 | |
| $\sigma_{w_{\text{REST2}}}$ | 0.59 | |
| $\rho_{r_{\text{CUFF1}},r_{\text{REST2}}}$ | 0.30 | |
| $\rho_{r_{\text{CUFF1}},w_{\text{CUFF1}}}$ | -0.35 | |
| $\rho_{r_{\text{CUFF1}},w_{\text{REST2}}}$ | -0.29 | |
| $\rho_{r_{\text{REST2}},w_{\text{CUFF1}}}$ | -0.03 | |
| $\rho_{r_{\text{REST2}},w_{\text{REST2}}}$ | -0.17 | |
| $\rho_{w_{\text{CUFF1}},w_{\text{REST2}}}$ | 0.20 | |
| $\beta_{\text{TKA}}$ | 0.44 | 0.27 |

Table S5: Proportion of invalid predicted score and intervals by LMM on A2CPS Cuff Pressure Pain dataset. Invalid scores are defined as those falling outside the valid range [0, 20]. Invalid intervals are defined as those with either lower or upper bound falling outside the valid range.

| Cohort | Period | Location | Number of completed assessments | Proportion of invalid score | Proportion of invalid interval |
|---|---|---|---|---|---|
| Knee Arthroplasty | CUFF1 | beginning | 347 | 0 | 0.236 |
| Knee Arthroplasty | CUFF1 | middle | 345 | 0 | 0.099 |
| Knee Arthroplasty | CUFF1 | end | 344 | 0 | 0.166 |
| Knee Arthroplasty | REST2 | beginning | 356 | 0.146 | 0.834 |
| Knee Arthroplasty | REST2 | middle | 356 | 0.053 | 0.834 |
| Knee Arthroplasty | REST2 | end | 356 | 0.171 | 0.840 |
| Thoracic | CUFF1 | beginning | 108 | 0 | 0.194 |
| Thoracic | CUFF1 | middle | 108 | 0 | 0.046 |
| Thoracic | CUFF1 | end | 106 | 0 | 0.123 |
| Thoracic | REST2 | beginning | 106 | 0.217 | 0.887 |
| Thoracic | REST2 | middle | 106 | 0.160 | 0.887 |
| Thoracic | REST2 | end | 106 | 0.292 | 0.906 |

Table S6: Posterior estimates by BBME on the FM study adjusting for treatment versus control, including mean of random intercepts ($r$), mean of random slopes ($w$), standard deviations of random effects ($\sigma$), dispersion ($\phi$), and covariate effect. All parameters converged with $\hat{R} < 1.05$. All correlation parameters are shown in the Figure S16(a).

| Parameter | Posterior mean | SD | Posterior median | 2.5% quantile | 97.5% quantile | ESS |
|---|---|---|---|---|---|---|
| $r_{wk1}$ | 0.26 | 0.07 | 0.26 | 0.12 | 0.41 | 589 |
| $r_{wk2}$ | 0.47 | 0.07 | 0.47 | 0.32 | 0.61 | 562 |
| $r_{wk3}$ | -0.23 | 0.10 | -0.23 | -0.44 | -0.03 | 1018 |
| $r_{wk4}$ | -0.14 | 0.10 | -0.14 | -0.35 | 0.06 | 1119 |
| $w_{wk1}$ | 0.01 | 0.00 | 0.01 | 0.00 | 0.01 | 4024 |
| $w_{wk2}$ | 0.00 | 0.00 | 0.00 | 0.00 | 0.01 | 5212 |
| $w_{wk3}$ | 0.00 | 0.00 | 0.00 | 0.00 | 0.01 | 4613 |
| $w_{wk4}$ | 0.00 | 0.00 | 0.00 | 0.00 | 0.00 | 4464 |
| $\beta_{treatment}$ | 0.00 | 0.06 | 0.00 | -0.12 | 0.13 | 237 |
| $\sigma_{r_{wk1}}$ | 0.63 | 0.04 | 0.63 | 0.56 | 0.72 | 4774 |
| $\sigma_{r_{wk2}}$ | 0.60 | 0.04 | 0.60 | 0.52 | 0.69 | 4248 |
| $\sigma_{r_{wk3}}$ | 1.01 | 0.07 | 1.01 | 0.89 | 1.16 | 5106 |
| $\sigma_{r_{wk4}}$ | 1.01 | 0.07 | 1.01 | 0.89 | 1.16 | 5150 |
| $\sigma_{w_{wk1}}$ | 0.02 | 0.00 | 0.02 | 0.01 | 0.02 | 3289 |
| $\sigma_{w_{wk2}}$ | 0.02 | 0.00 | 0.02 | 0.02 | 0.03 | 4457 |
| $\sigma_{w_{wk3}}$ | 0.02 | 0.00 | 0.02 | 0.01 | 0.02 | 2426 |
| $\sigma_{w_{wk4}}$ | 0.02 | 0.00 | 0.02 | 0.01 | 0.02 | 3198 |
| $\phi$ | 36.53 | 0.60 | 36.53 | 35.35 | 37.70 | 6277 |

Table S7: LMM estimates adjusting for covariate on the FM study. All correlation parameters are shown in the Figure S16(b).

| Parameter | Estimate | Std Error |
|---|---|---|
| $r_{wk1}$ | 74.53 | 2.82 |
| $r_{wk2}$ | 80.66 | 2.72 |
| $r_{wk3}$ | 61.44 | 3.34 |
| $r_{wk4}$ | 63.81 | 3.34 |
| $w_{wk1}$ | 0.16 | 0.05 |
| $w_{wk2}$ | 0.04 | 0.05 |
| $w_{wk3}$ | 0.10 | 0.05 |
| $w_{wk4}$ | 0.03 | 0.05 |
| $\beta_{treatment}$ | -0.42 | 2.90 |
| $\sigma_{r_{wk1}}$ | 18.63 | |
| $\sigma_{r_{wk2}}$ | 16.61 | |
| $\sigma_{r_{wk3}}$ | 26.14 | |
| $\sigma_{r_{wk4}}$ | 26.13 | |
| $\sigma_{w_{wk1}}$ | 0.46 | |
| $\sigma_{w_{wk2}}$ | 0.50 | |
| $\sigma_{w_{wk3}}$ | 0.44 | |
| $\sigma_{w_{wk4}}$ | 0.44 | |

#### Supplementary Figures

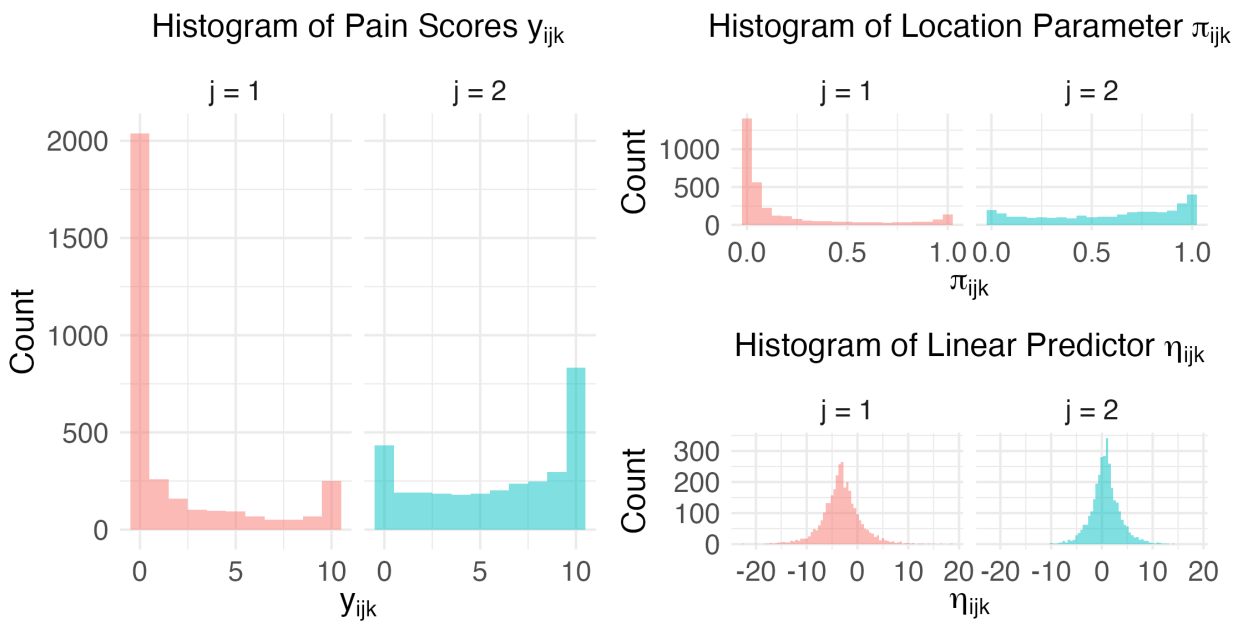

Figure S1: Histograms of simulated data for zero-inflated simulation study in Appendix 6.1. Pain scores, location parameters ($\pi_{ijk}$), and linear predictors ($\eta_{ijk}$) from one simulated dataset of 500 subjects, up to 2 time points, and 4-10 pain scores per time point. Random effects were sampled from $\begin{bmatrix} r_{i1} \\ r_{i2} \\ w_{i1} \\ w_{i2} \end{bmatrix} \sim \text{MVN}_4\left(\begin{bmatrix} -3.2 \\ 0.6 \\ 0 \\ 0 \end{bmatrix}, \begin{bmatrix} 1.5^2 & 0.5\times1.5\times1 & 0.3\times1.5\times0.6 & 0 \\ 0.5\times1.5\times1 & 1^2 & -0.3\times1\times0.6 & 0 \\ 0.3\times1.5\times0.6 & -0.3\times1\times0.6 & 0.6^2 & 0 \\ 0 & 0 & 0 & 0.5^2 \end{bmatrix}\right)$ with timepoint-specific dispersion $\phi_{j=1} = 5$ and $\phi_{j=2} = 10$.

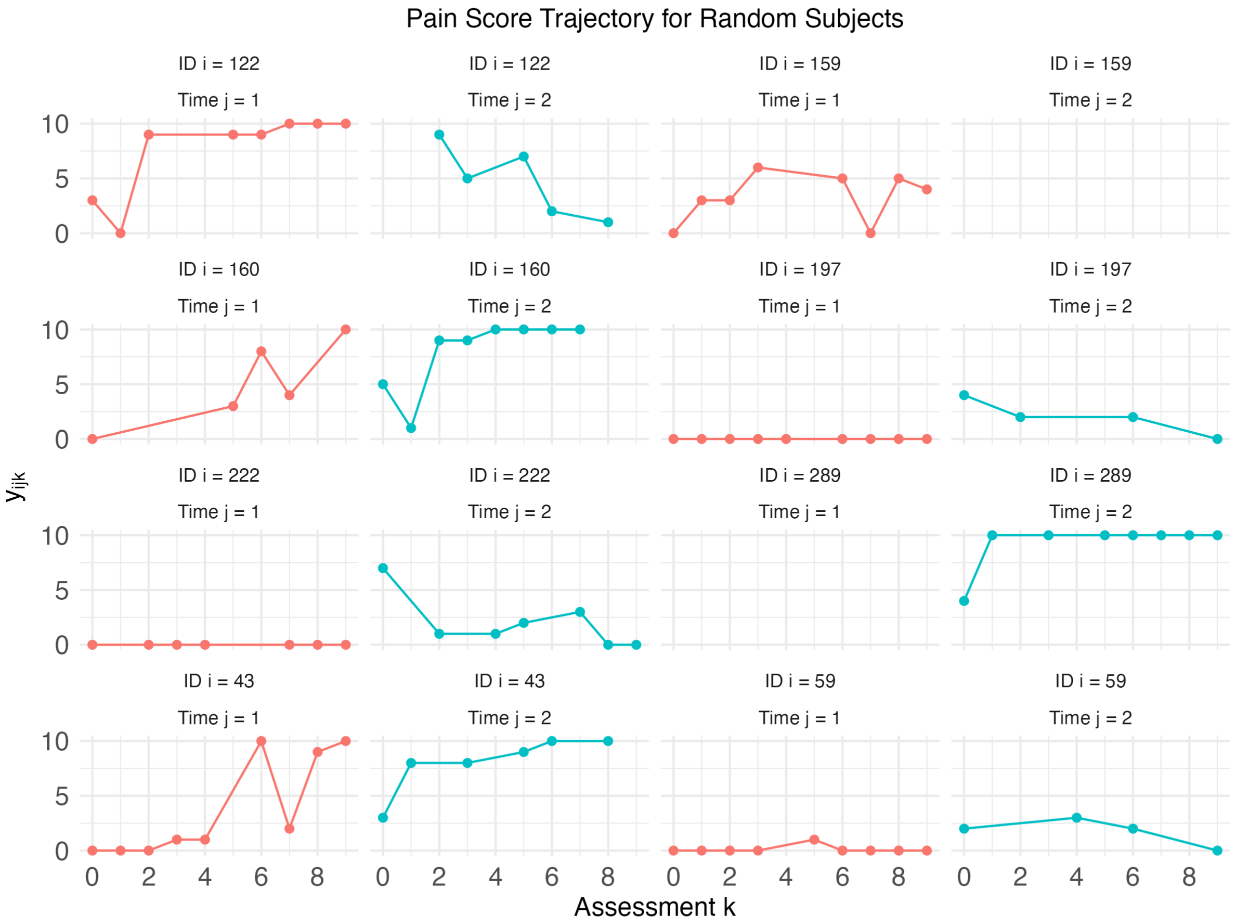

Figure S2: Pain trajectories of randomly sampled subjects from the simulated data in zero-inflation example study in Appendix 6.1. Pain levels are on average lower in timepoint $j = 1$ than in $j = 2$. Parameters used for data simulation are stated in Appendix 5.1.

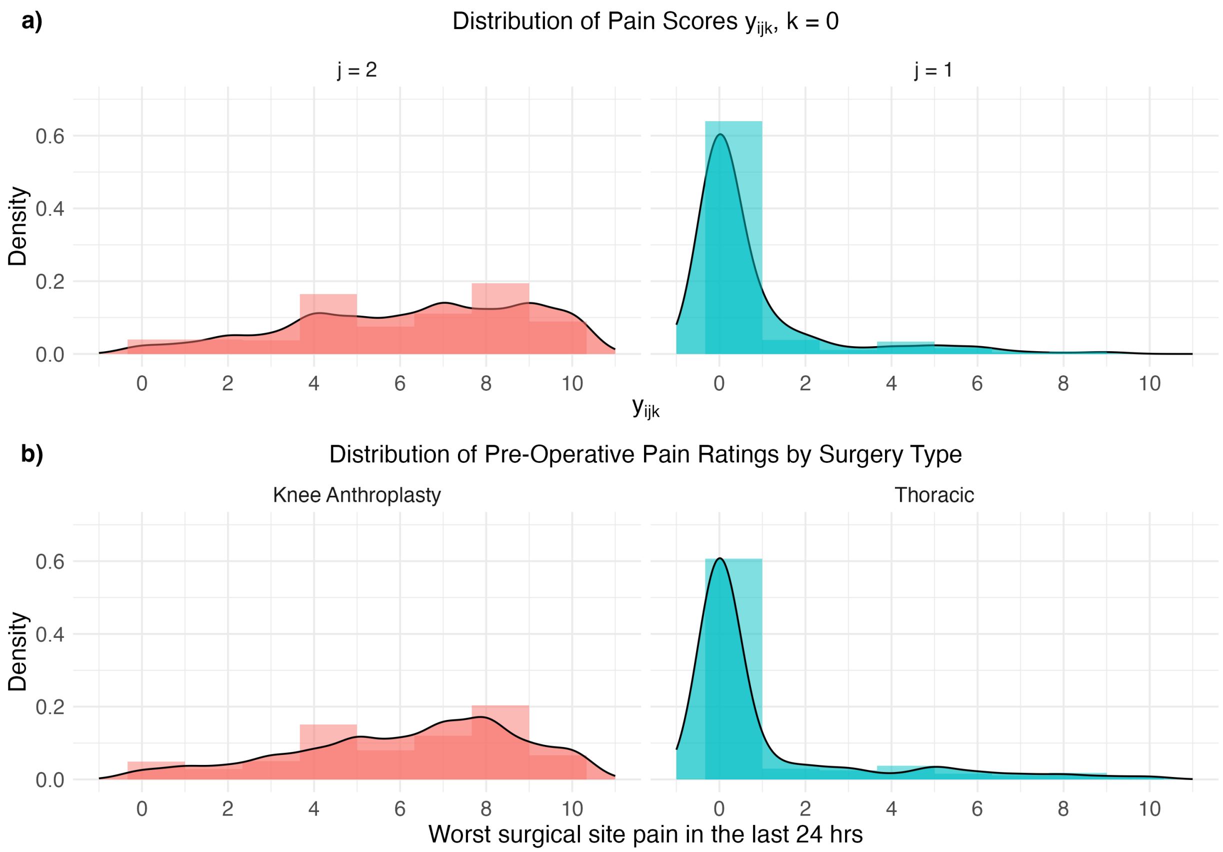

Figure S3: Comparison of the distribution of a) simulated pain scores for zero-inflation simulation in Appendix 6.1 and b) observed A2CPS Brief Pain Inventory (BPI) worst surgical-site pain scores (Figure 2). Simulation parameter values were chosen to mimic the distribution of A2CPS baseline worst surgical-site pain scores as an example of zero-inflated data.

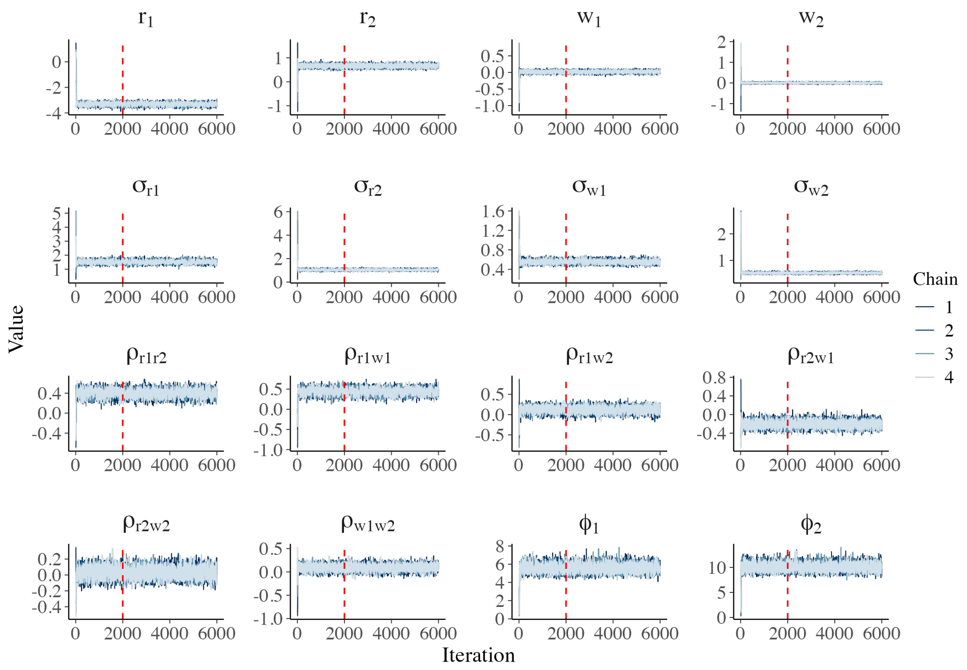

(a)

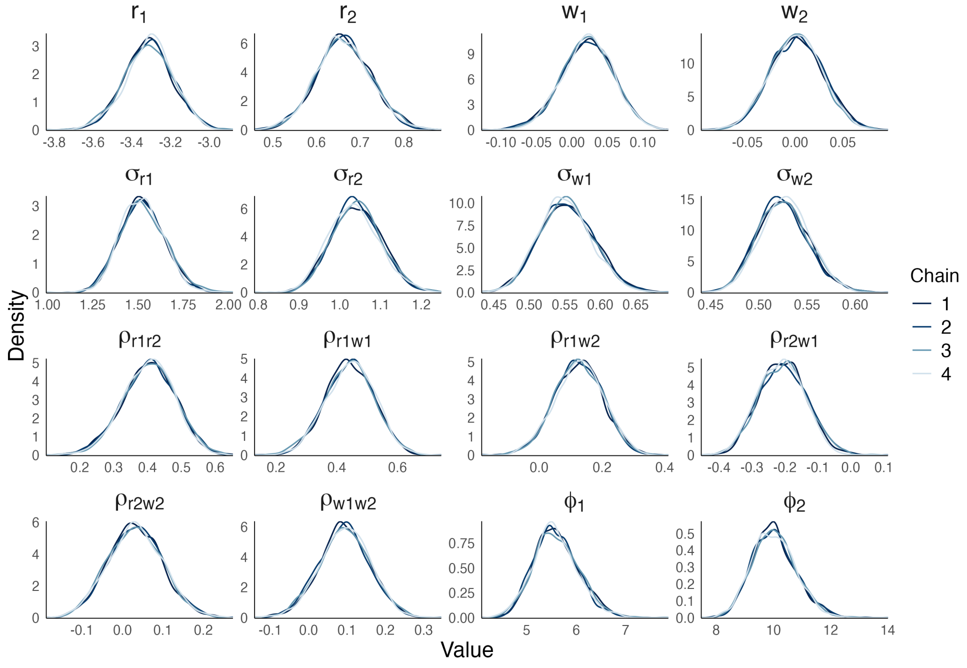

(b)

Figure S4: Simulation results of full BBME model with timepoint-specific dispersion for zero-inflation simulation study in Appendix 6.1. The model was fit using multivariate normal prior with mean **0** and covariance matrix $5^2 I_{2J}$ on the mean of random effects, half-cauchy(0, 5) on standard deviations of random effects, LKJ(1) on correlations, and half-cauchy(0, 20) on dispersions. (a) Trace plots of four MCMC chains (burn-in period shown before the red dashed line, thinning factor = 2). (b) Density plots of the four chains after burn-in, showing good mixing and overlap across chains.

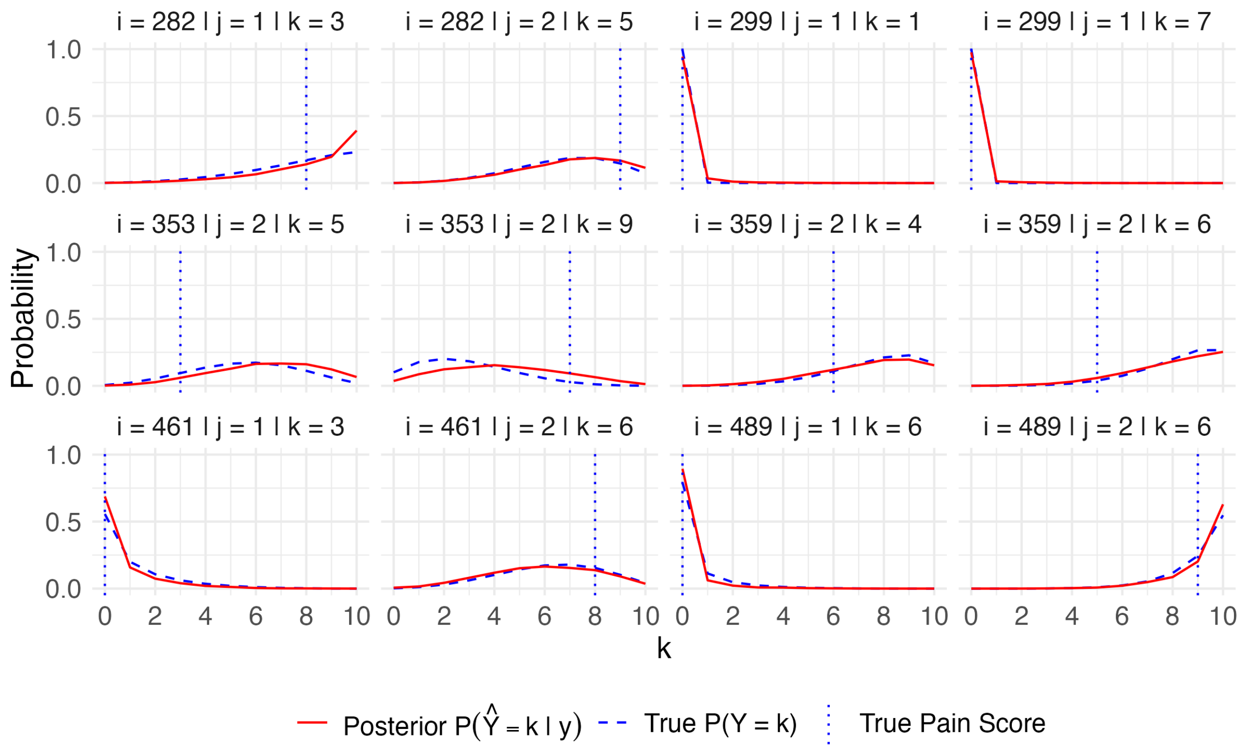

Figure S5: True probability mass function curve and posterior mass at score = $k$ from posterior samples for the pain scores of six randomly selected subjects in zero-inflation simulation study in Appendix 6.1. The posterior curves closely align with the true probability curves, demonstrating good model fit.

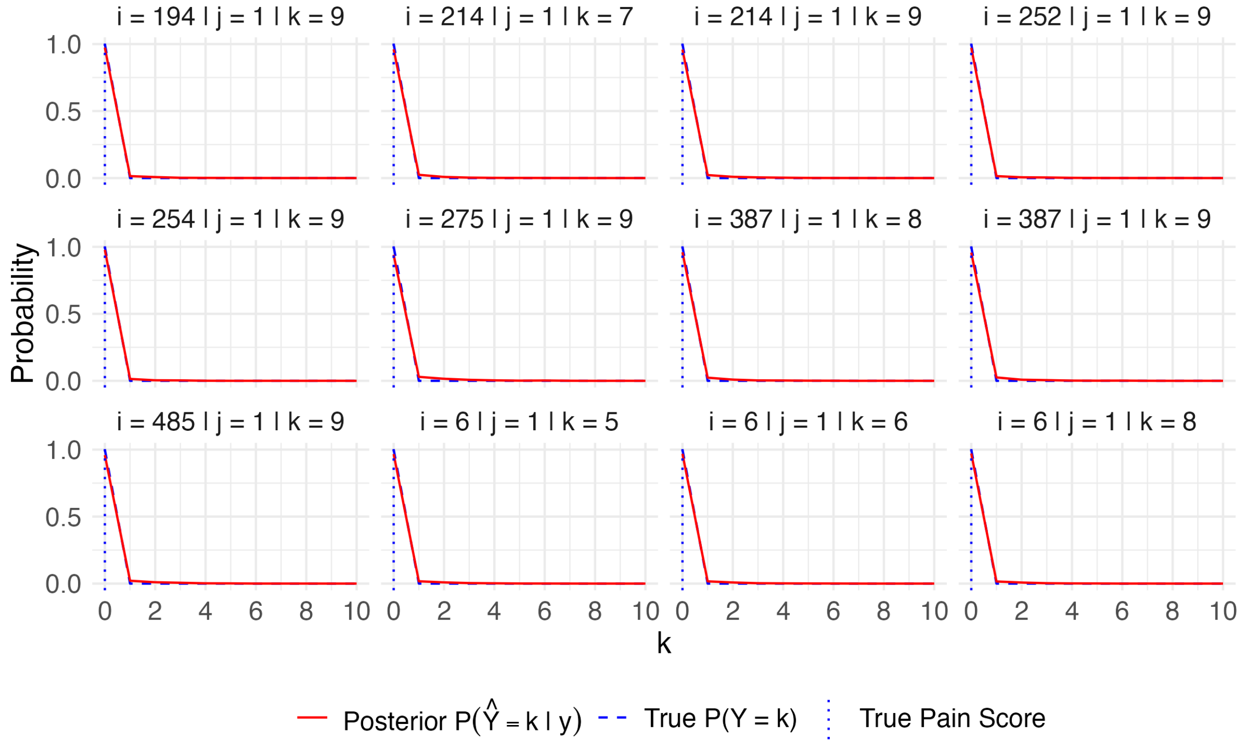

Figure S6: True probability mass function curve and posterior mass at k from posterior samples for the pain scores with the greatest discrepancy between true and predicted linear predictor $\eta_{ijk}$ in zero-inflation simulation study in Appendix 6.1. Despite differences in $\eta_{ijk}$, the posterior curves align closely with the true curves, indicating robustness of the model fit.

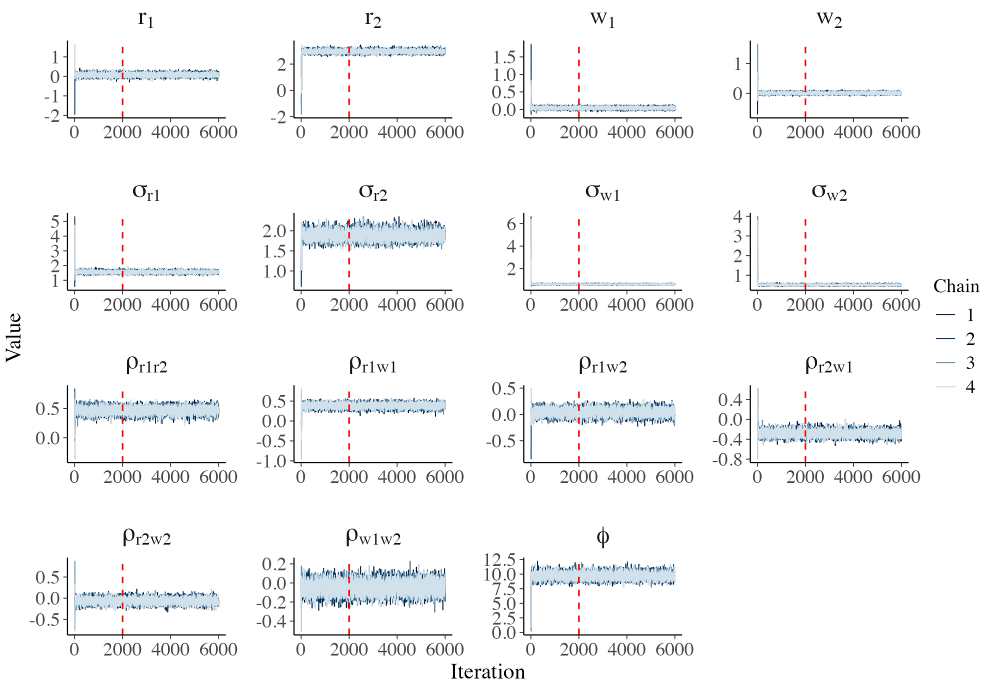

(a)

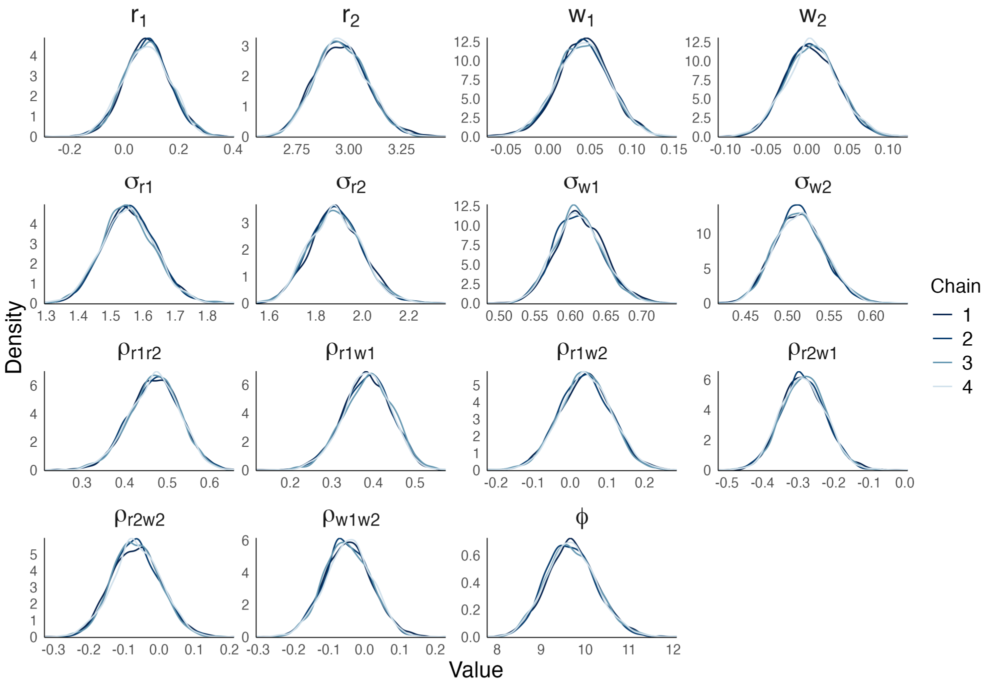

(b)

Figure S7: Simulation results of full BBME model with common dispersion in the one-inflation simulation study in Appendix 6.2. (a) Trace plots of four MCMC chains (burn-in period shown before the red dashed line, thinning factor = 2). (b) Density plots of the four chains after burn-in, showing good mixing and overlap across chains.

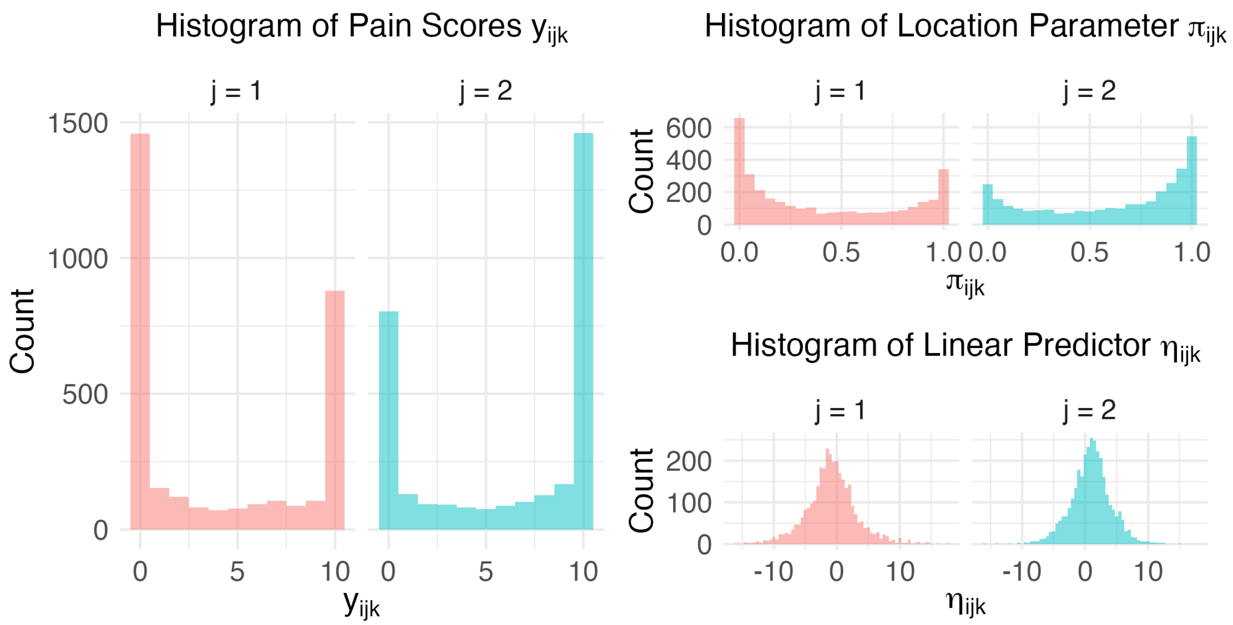

Figure S8: Histograms of pain scores, location parameters, and linear predictors from a simulated dataset of 500 subjects, each observed at up to 2 time points with 4-10 pain scores per time points. Random effects were sampled from $\begin{bmatrix} r_{i1} \\ r_{i2} \\ w_{i1} \\ w_{i2} \end{bmatrix} \sim \text{MVN}_4\left(\begin{bmatrix} -1 \\ 1 \\ 0 \\ 0 \end{bmatrix}, \begin{bmatrix} 1.5^2 & 0.5\times1.5\times2 & 0.3\times1\times0.6 & 0 \\ 0.5\times1.5\times2 & 2^2 & -0.3\times2\times0.6 & 0 \\ 0.3\times1\times0.6 & -0.3\times2\times0.6 & 0.6^2 & 0 \\ 0 & 0 & 0 & 0.5^2 \end{bmatrix}\right)$ with log of subject-level dispersion parameters given by random intercepts sampled from $N(0,1)$.

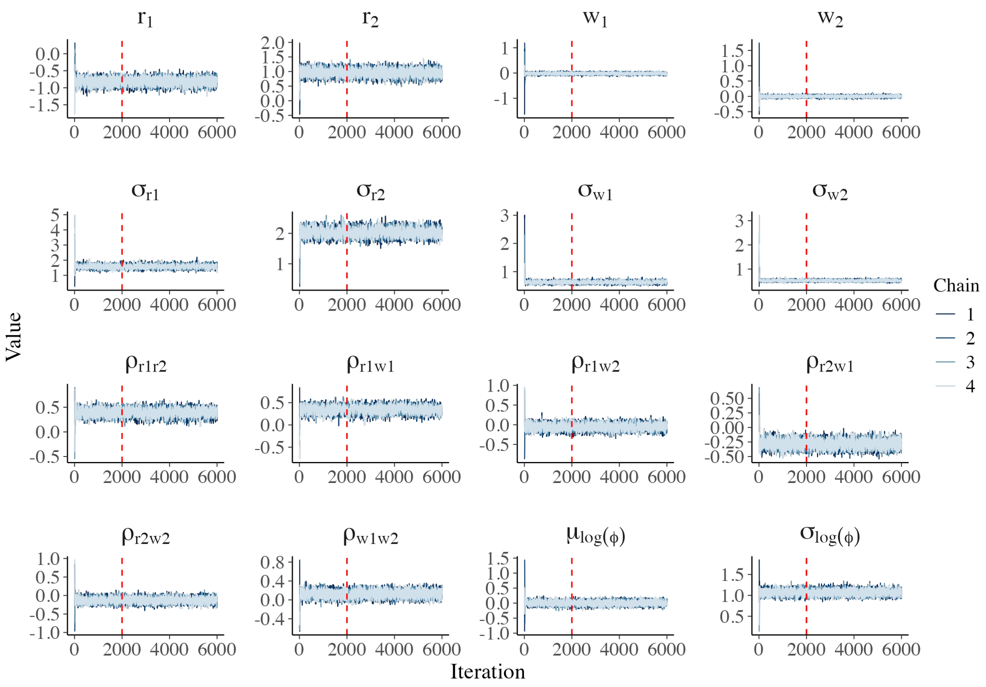

(a)

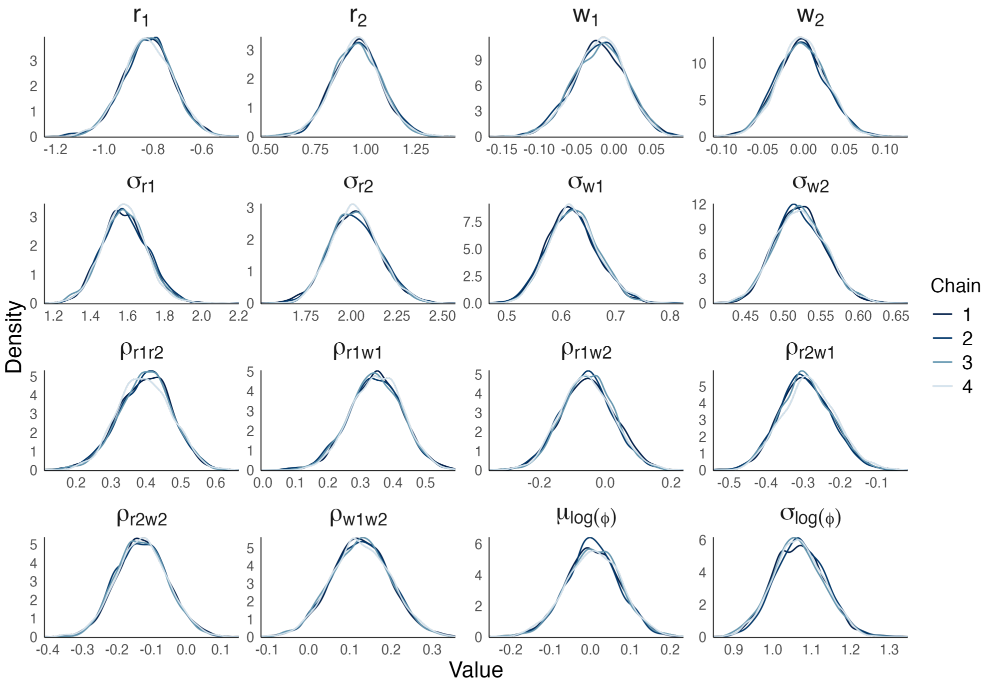

(b)

Figure S9: Simulation results of the BBME model with subject-level dispersion. (a) Trace plots of four MCMC chains with burn-in iterations shown before the red dashed line (thinning = 2). and (b) Posterior density plots of the four chains after burn-in, demonstrating good mixing and convergence.

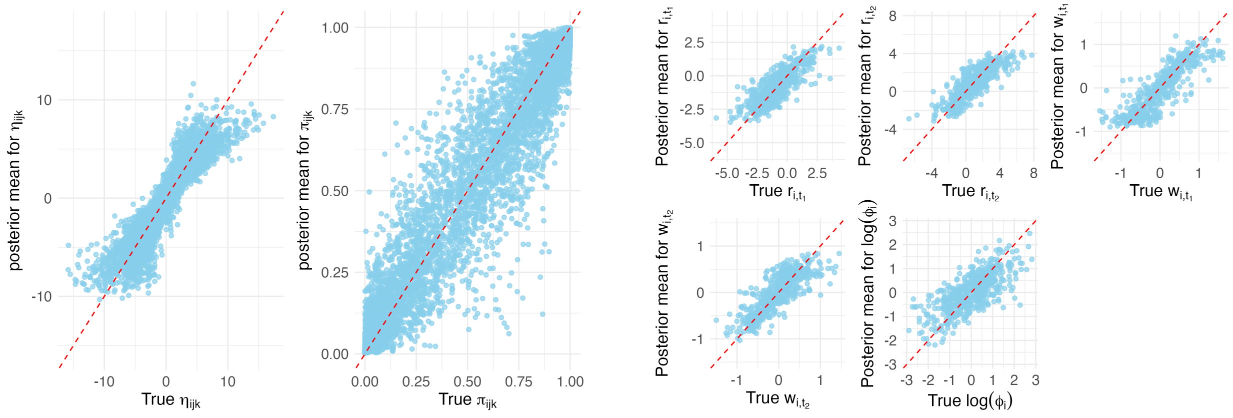

(a) (b)

Figure S10: Simulation results of BBME model with subject-level dispersion. (a) Scatter plot of the true linear predictor and true location parameter against posterior means for each pain score. (b) Scatter plot of posterior means of random effects against their true values. The red line indicates the identity line.

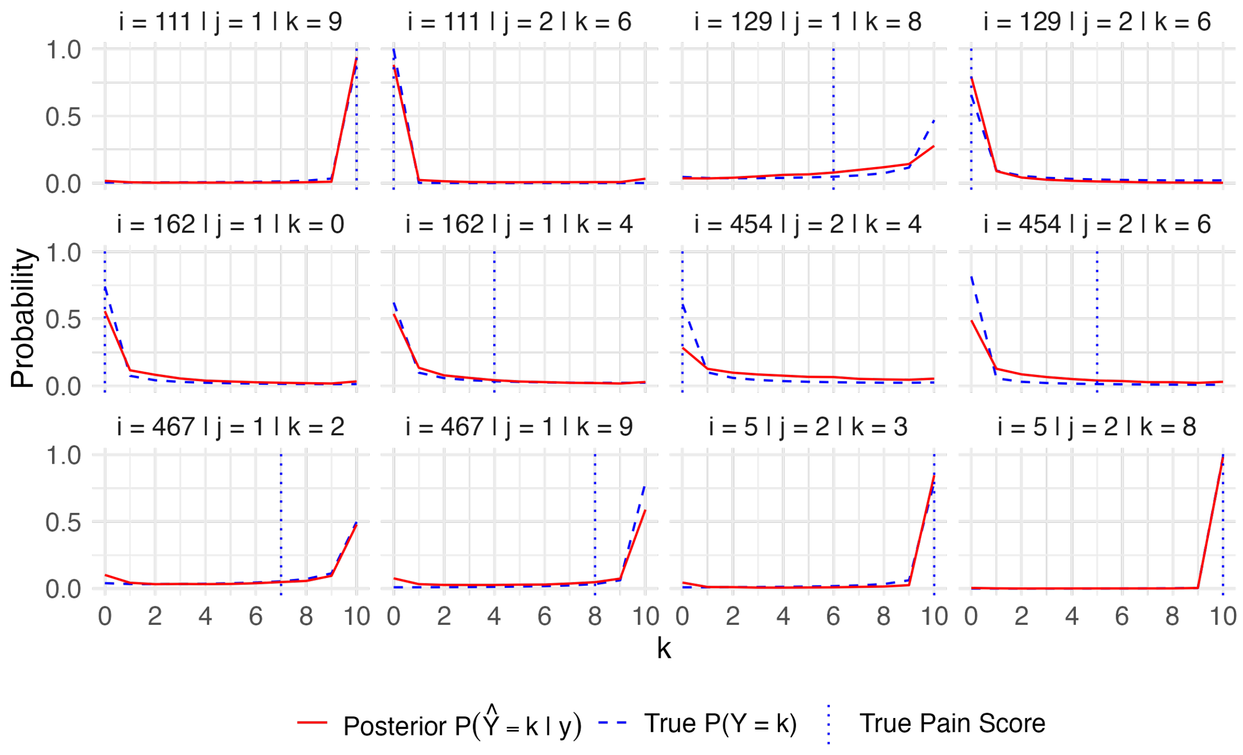

Figure S11: True and posterior mean probability mass function curves of pain scores for six randomly selected subjects, fitted in BBME with subject-level dispersion. The posterior mean curves closely match the true curves.

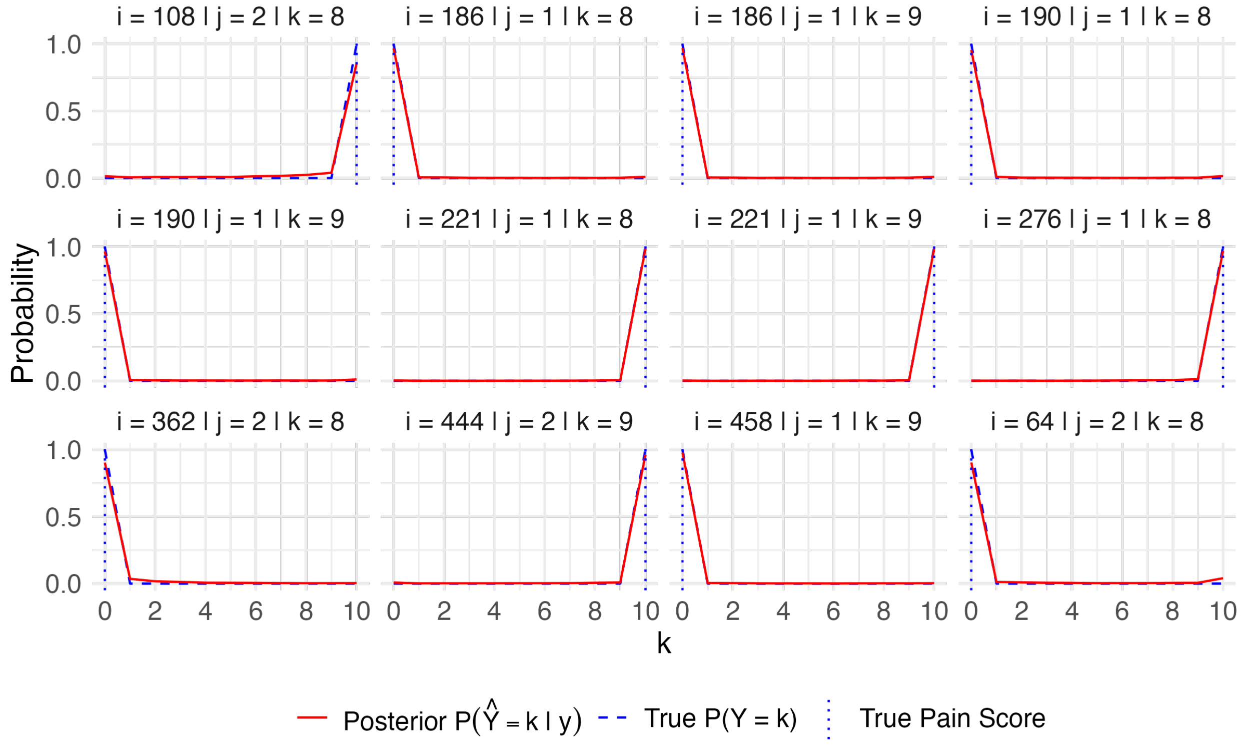

Figure S12: True and posterior mean probability mass function curves of pain scores with the greatest differences between true and predicted linear predictor $\eta_{ijk}$, fitted in BBME with subject-level dispersion. Despite the discrepancies in $\eta_{ijk}$, the posterior probability mass functions remain close to the true curves.

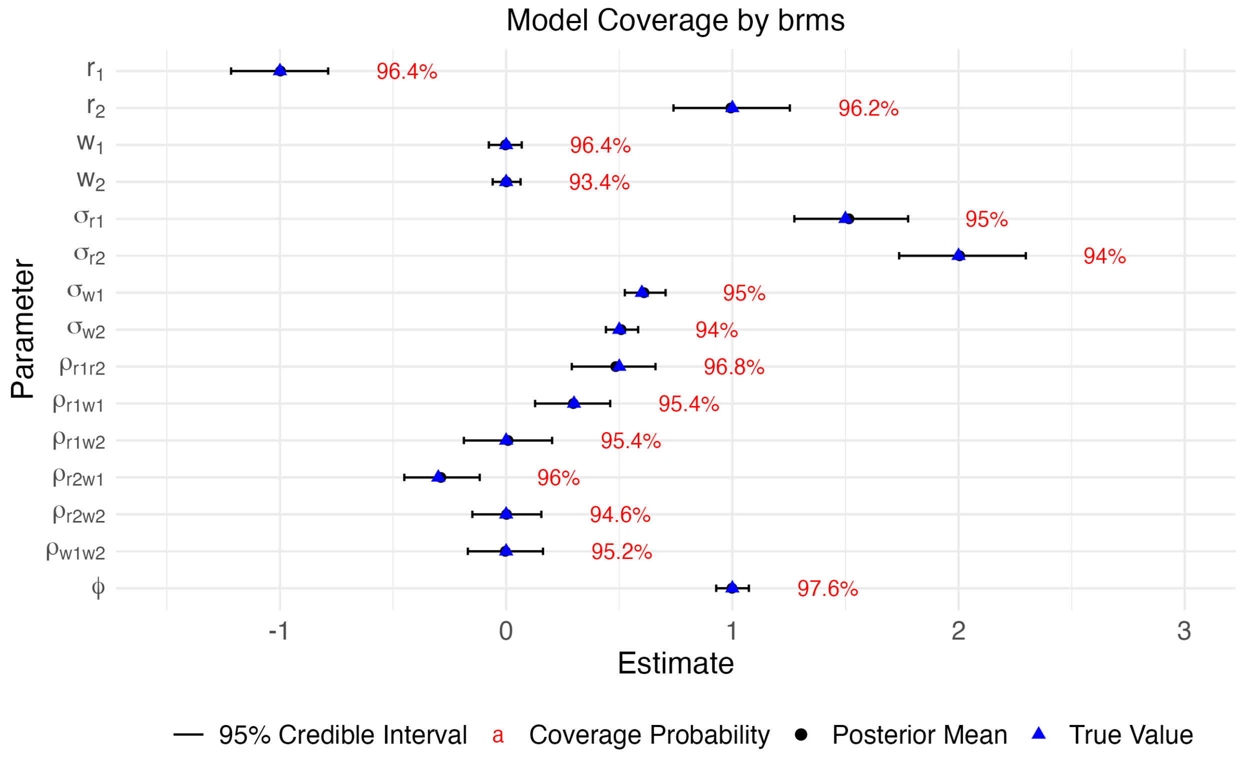

Figure S13: Mean of posterior mean, 95% credible intervals, and coverage probabilities from 500 simulated datasets fitted using brms [2]. Black lines indicate 95% credible intervals, blue triangles denote true parameter values, and red labels show coverage probabilities.

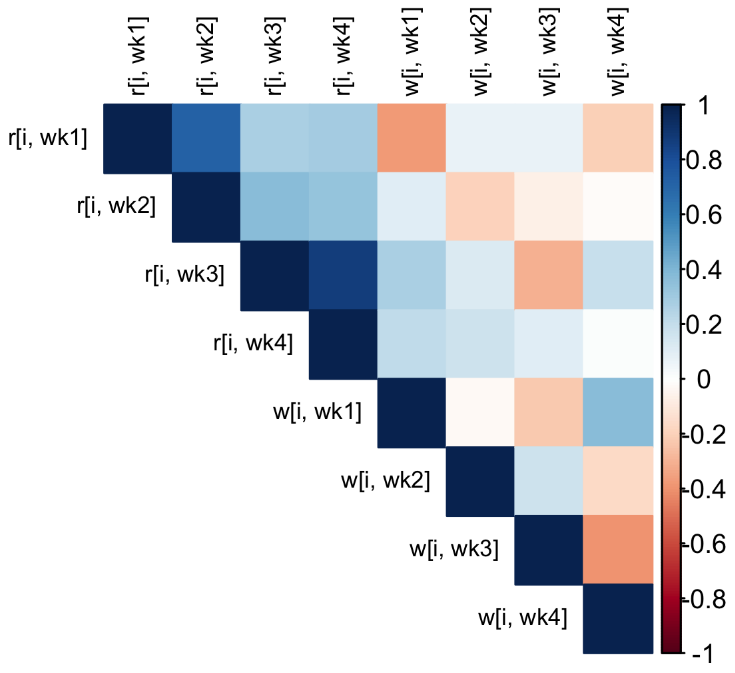

(a)

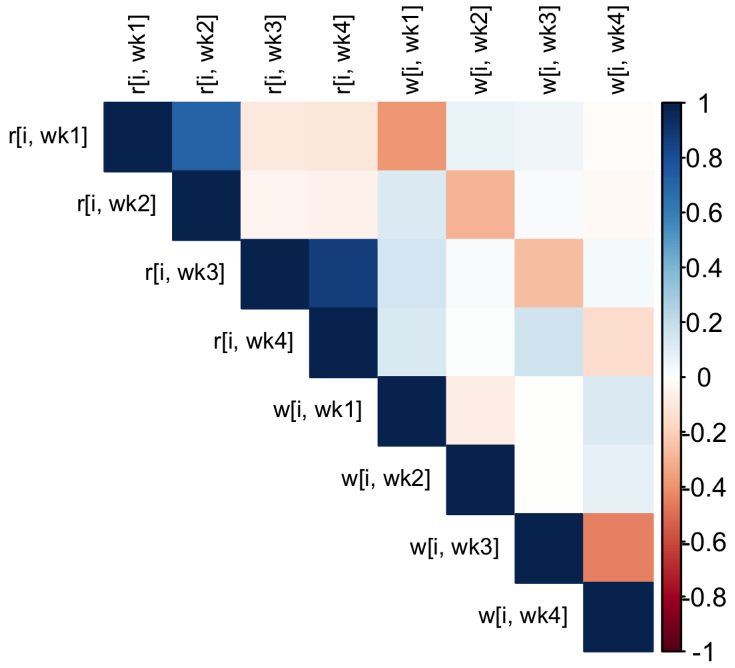

(b)

Figure S14: Correlation estimates for the FM study by BBME and LMM model. a) Posterior estimates of correlations among random effects by BBME with common dispersion. All parameters converged with $\hat{R} = 1$. b) LMM-estimated correlations among random effects.

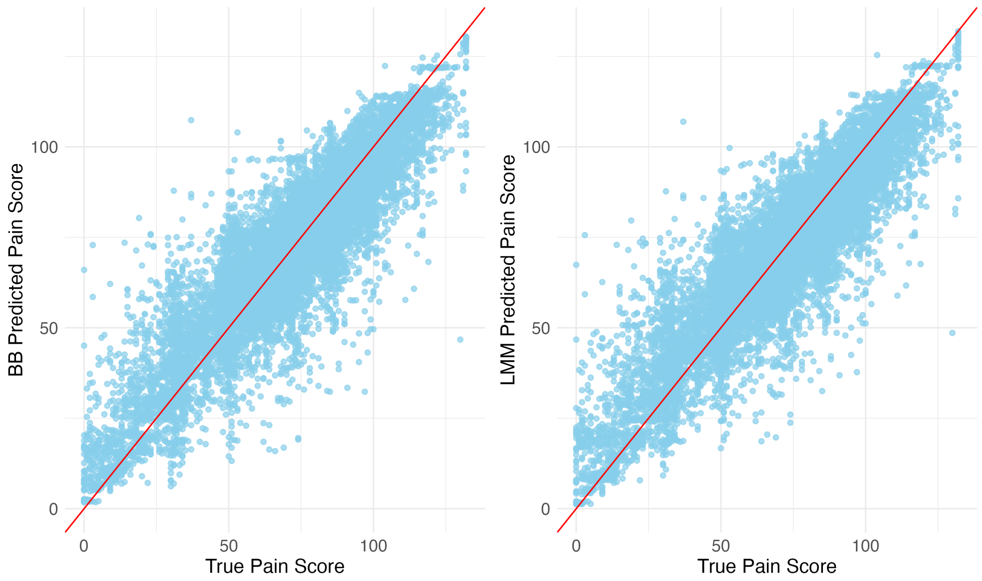

Figure S15: Scatterplot of predicted pain score by BBME and LMM against true pain scores. Predicted scores were similar for the non-inflated dataset in FM study. The red line indicates the identity line.

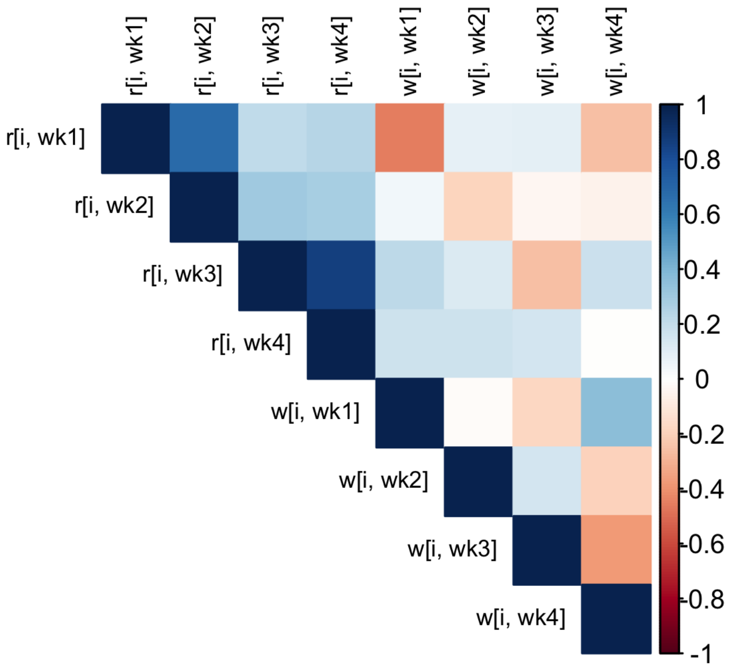

(a)

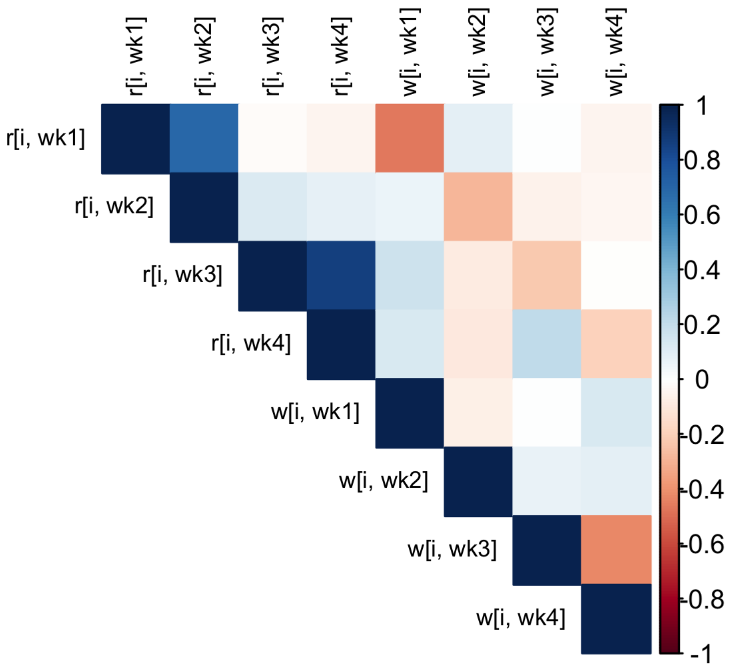

(b)

Figure S16: Correlation estimates for the FM study by common-dispersion BBME and LMM model adjusting for covariate. a) Posterior estimates of correlations among random effects by BBME. All parameters converged with $\hat{R} < 1.05$. b) LMM-estimated correlations among random effects.

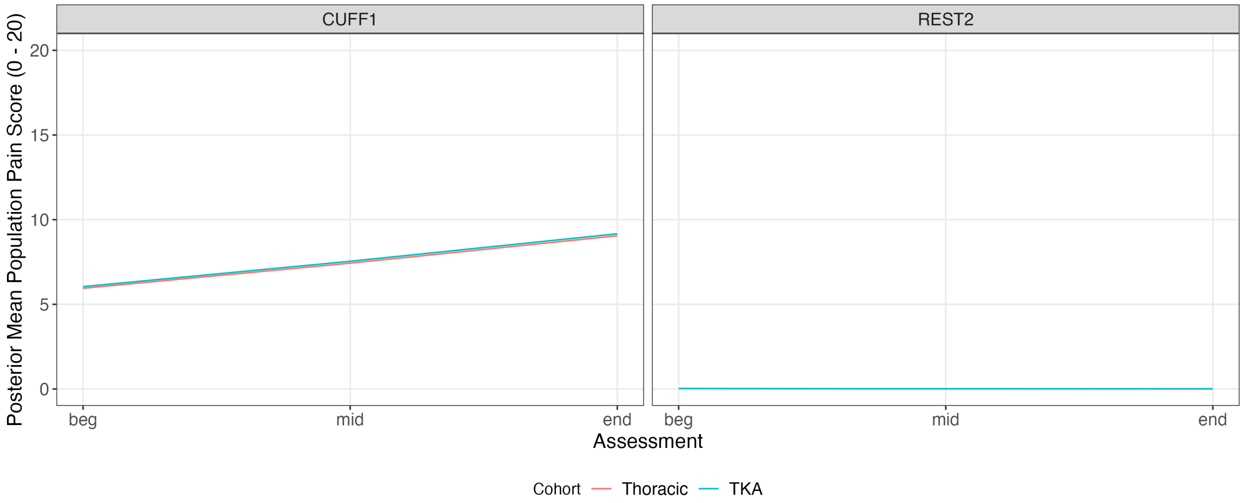

Figure S17: Population level BBME-estimated posterior mean pain scores at CUFF1 and REST2 for thoracic and TKA subjects in A2CPS cuff pain dataset.

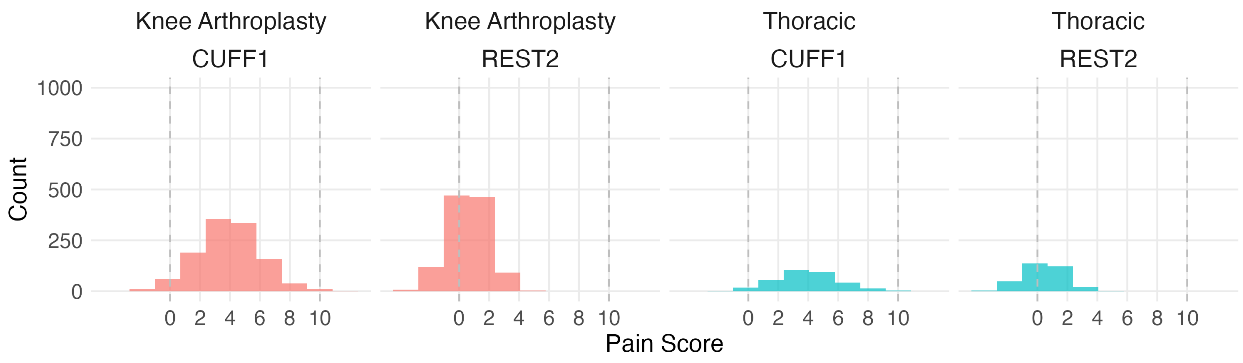

Figure S18: Resampled dataset using LMM population estimates on A2CPS cuff pain dataset.

### References

[1] Brooks M, Kristensen K, van Benthem K, Magnusson A, Berg C, Nielsen A, Skaug H, Mächler M, Bolker B. glmmTMB balances speed and flexibility among packages for zero-inflated generalized linear mixed modeling. The R Journal 2017;9:378–400.

[2] Bürkner P. brms: An R Package for Bayesian Multilevel Models Using Stan. J Stat Soft 2017;80:1.

[3] Carpenter B, Gelman A, Hoffman MD, Lee D, Goodrich B, Betancourt M, Brubaker M, Guo J, Li P, Riddell A. Stan: A probabilistic programming language. Journal of statistical software 2017;76:1–32.

[4] Gelman A, Rubin DB. Inference from iterative simulation using multiple sequences. Statistical Science 1992;7:457–472.

[5] Vehtari A, Gelman A, Gabry J. Practical Bayesian model evaluation using leave-one-out cross-validation and WAIC. Statistics and Computing 2017;27:1413–1432.
